# Environmental fluctuations explain the universal decay of species-abundance correlations with phylogenetic distance

**DOI:** 10.1101/2022.07.12.499693

**Authors:** Matteo Sireci, Miguel A. Muñoz, Jacopo Grilli

**Affiliations:** Departamento de Electromagnetismo y Física de la Materia e Instituto Carlos I de Física Teórica y Computacional. Universidad de Granada. E-18071, Granada, Spain; Quantitative Life Sciences, The Abdus Salam International Centre for Theoretical Physics, 34151 Trieste, Italy

**Author notes:** Electronic address (Matteo Sireci). Electronic address (Miguel A. Muñoz). Electronic address (Jacopo Grilli).

## Abstract

Multiple ecological forces act together to shape the composition of microbial communities. *Phyloecology* approaches —which combine phylogenetic relationships with community ecology— have the potential to disentangle such forces, but are often hard to connect with quantitative predictions from theoretical models. On the other hand, *macroecology*, which focuses on statistical patterns of abundance and diversity, provides natural connections with theoretical models but often neglects inter-speficic correlations and interactions. Here, we propose a unified framework combining both such approaches to analyze microbial communities. In particular, by using both cross-sectional and longitudinal metagenomic data for species abundances, we reveal the existence of a novel empirical macroecological law establishing that correlations in species-abundance fluctuations across communities decay from positive to null values as a function of phylogenetic similarity in a consistent manner across ecologically distinct microbiomes. We formulate three mechanistic models —relying on alternative ecological forces— that lead to radically different predictions. We conclude that the empirically observed macroecological pattern can be quantitatively explained as a result of shared fluctuating resources, i.e. *environmental filtering* and not e.g. as a consequence of species competition. Finally, we also show that the macroecological law is also valid for temporal data of a single community, and that the properties of delayed temporal correlations are reproduced by the model with environmental filtering.

---

Microbial communities are ubiquitous on earth, from human microbiota, to ocean, soil, and glacial environments [1]. Their widespread presence is paralleled by their complex and highly variable composition, both across time and space [2]. Understanding what are the main drivers, or *“ecological forces”*, shaping the coexistence and stability of microbial communities under changing environmental conditions and perturbations is a fundamental challenge of utmost relevance for, e.g., environmental and health sciences.

Ecological forces can emerge from the interactions between species or between species and the environment, including both biotic and abiotic factors. Experiments in simple and controlled laboratory environments have made it possible to trace the effects of various ecological forces on community composition, often reshaping classical ideas on ecological interactions [3–8]. For instance, cross-feeding has emerged as a central player in determining community assembly and species coexistence [9, 10]. However, the precise role of different ecological forces in determining composition and variation in more complex natural communities remains mostly unknown. While detailed information about environmental [11–13] and genetic [14–16] factors shaping interaction and response to environmental conditions is sometimes available, we still lack frameworks to infer their quantitative strength and to disentangle the relative relevance of each of the acting ecological forces from available data [17–19].

Macroecology —i.e. the study of ecological communities through the analysis of global patterns of abundance, diversity, and distribution [21]— stands as a prominent approach to link quantitative ecological models with empirical data of complex and diverse communities [22, 23]. In particular, in the context of microbial communities, a growing body of evidence reveals that the abundance dynamics observed in microbial communities is characterized by distinctive and reproducible statistical patterns, also known as *macroecological laws* [23–27]. Further evidence shows that, despite the complexity of the underlying “microscopic” dynamics, most of such patterns can be reproduced by relatively simple models —such as e.g. the stochastic logistic model (SLM)— capturing salient features of the underlying ecological forces [24–28]. However, such simplified models often neglect interactions between species, treating their abundance fluctuations as independent from each other. Nevertheless, it is noteworthy that including species interactions in models such as the SLM does not significantly affect the shape of single-species macroecological patterns. For instance, generalized Lotka-Volterra equations with environmental stochasticity —which reduce to the SLM in the absence of interactions— predict time-series statistics and patterns similar to those of the SLM [25–27].

On the other hand, it seems clear that the ecological forces shaping community composition and variability can only be unveiled within a macroecological approach by explicitly studying multispecies abundance patterns. For instance, empirically-determined pairwise correlations between species abundances can be partially explained by consumer-resource models with resource fluctuations [28].

One challenge in connecting empirical macroecological patterns with simple yet biologically-grounded models is that not all statistical patterns are equally informative. For instance, it is well known that, in many ecological systems, the empirical shape of the species abundance distribution (SAD) —i.e. one of the most prominent macroecological patterns— can be reproduced by models with very different underlying biological assumptions (such as, e.g. neutral and niche theories [29, 30]). Similarly, multiple mechanisms are expected to determine the observed correlations between species abundance fluctuations. Pairwise correlations are in fact the result of multiple ecological forces, such as competition, cooperation, cross-feeding, but also of indirect effects through a network of interactions [31]. Analyzing the phylogenetic structure of community composition [32, 33] is a standard approach to disentangling the effects of these alternative assembly mechanisms. This type of approach is generally applied to analyze species (co-)occurrence. For example, shared environmental fluctuations (called “*environmental filtering*” hereon) produce phylogenetic clustering (i.e., similar species share a tendency to be simultaneously present or absent [34]), while exclusion by limiting similarity determines phylogenetic overdispersion (i.e., similar species tend not to be simultaneously present). This type of phylogenetic approaches has been widely applied in plant communities as well as in other systems [35–37] including, in particular, microbial communities [38]. More generally, phyloecology, which combines phylogenetic relationships with community ecology, has the potential to reveal the processes determining community composition [39, 40]. However, with few notable exceptions focusing on testing neutral models [41, 42], a connection between empirical observations of community ecology based on phylogeny and quantitative predictions of theoretical models is still missing.

Here, our goal is to develop such a connection under the lens of macroecology. In particular, we first elucidate the existence of a new empirical macroecological law that associates pairwise abundance correlations with species phylogenetic similarity. To rationalize such a finding, we formulate three alternative theoretical models —each relying on different ecological forces— all of which reproduce previously studied single-species macroecological patterns [25–27], but lead to radically different predictions for phylogenetic-dependent correlation patterns. These analyses allow us to conclude that only *environmental filtering* (and not, e.g., species competition) explains the empirically-observed pattern of decaying correlations with phylogenetic distance. Last but not least, we analyze temporal data for a fixed community, showing that the macroecological law also holds quantitatively in this context and that delayed temporal correlations are naturally reproduced by our model with environmental filtering.

## Results

### The averaged correlation of abundance fluctuations decays with phylogenetic distance in a consistent fashion

We consider the phylogenetic (or cophenetic) distance, *d_G,ij_* (where the subindex *G* stands for “genetic”) for each pair of operational taxonomic units (OTUs) (*i, j*), by using publicly-available results from 16*S* ribosomic RNA analyses [20, 43]. This genetic distance exhibits a broad variability across OTU pairs (see Methods and Fig. S1). For each pair of OTUs, we measure the correlation between the corresponding abundance fluctuations *η_ij_* (see Fig. 1(a) and Methods) across samples. Fig. 1(b) illustrates the value of the pairwise correlation *η*, averaged over the pairs of OTUs at a given phylogenetic distance (where distances are grouped into discrete intervals or bins) for diverse biomes. Remarkably, the resulting averaged correlation is found to decay with the phylogenetic distance, *d_G_*, in a robust way across environments and datasets. In particular, phylogenetically close OTUs (small values of *d_G_*) display a significant positive correlation while, for distant OTUs, the average correlation decreases to zero.

**FIG. 1:**
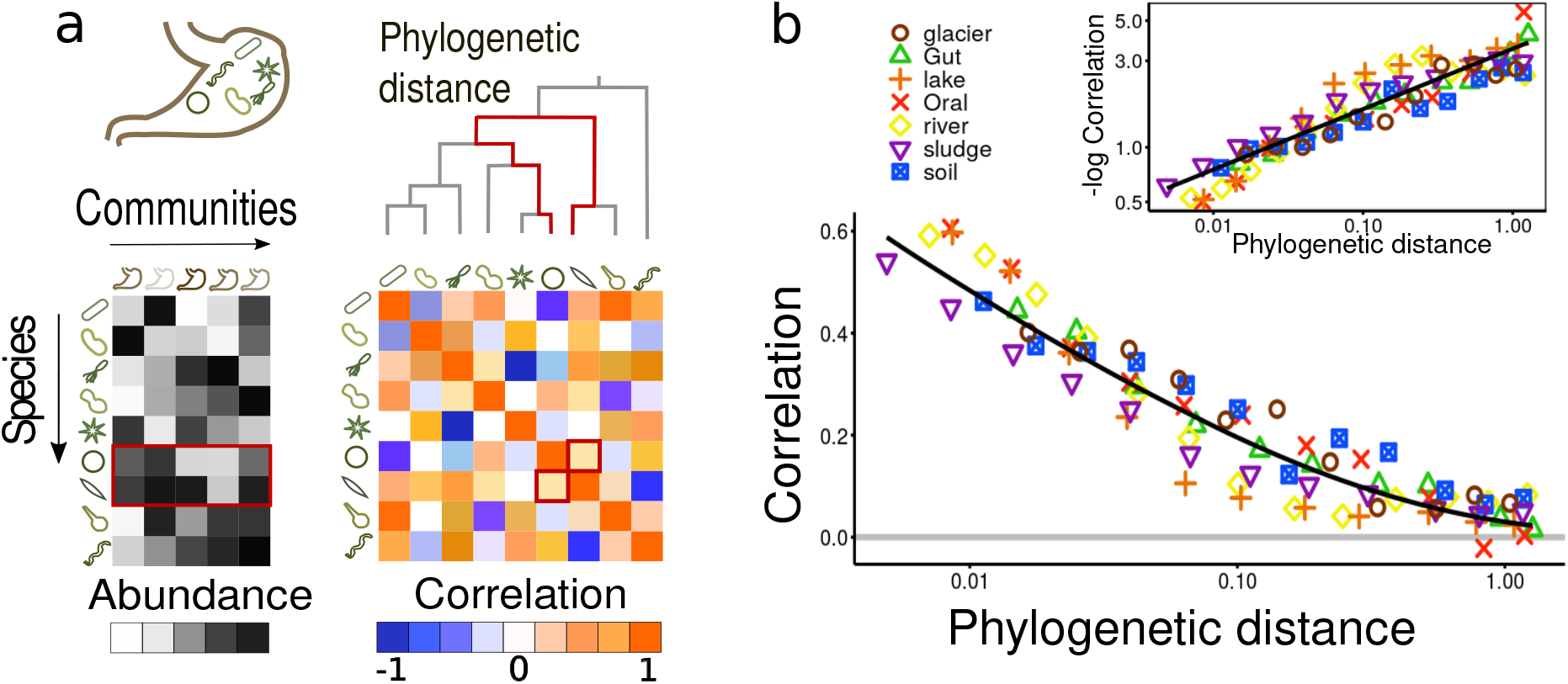
**(a) Pictorial illustration of the data organization and statistical analyses**. Abundances of different species (i.e. OTU at 97% similarity [20]), for different communities of the same biome (e.g gut of different hosts) are collected, respectively in rows and columns of the left table. The grey scale in the matrix entries stands for the level of abundance with darker shades corresponding to more abundant species. The (symmetric) species-abundance correlation matrix (color coded) is obtained by calculating for each pair of existing species the correlation of abundance fluctuations across communities. Finally, the phylogenetic distance is computed for all possible pairs of species by reconstructing the phylogenetic tree; and than associated with the couple correlation. The abundances, correlation, and phylogenetic distance of two given species are emphasized in red color. **(b) Macroecological law for pairwise correlations as a function of the phylogenetic distance for different biomes**. The correlation of abundance fluctuation averaged over all couples within a given discretized distance bin (colored symbols) decays with the phylogenetic distance (log scale) for all the considered microbiomes (see legend). In particular, each bin in the x-axis includes all couples with a phylogenetic distance within such a discrete bin (each one including at least 10^3^ couples for each of the 8 considered biomes; as shown in the SI sec. S2,S3.A, the pairs are not uniformly distributed across phylogenetic distances: the vast majority of couples lie in the rightmost bins, with large distances and small pairwise correlation values). The black line represents a stretched-exponential decay, Eq.(1) with *λ* ≈ 3.5. To emphasize the functional dependence, the inset shows the same data but for the negative log of the correlations represented in double logarithmic scale (i.e., a plot in which stretched exponential functions become straight lines; in this case with slope 1/3).

We compare this observation with randomized data, obtained by shuffling the position of OTUs on the phylogenetic tree. Such a randomization tree preserves both the statistical properties of the abundances and the property of the tree while removing the relation between the two. The comparison with the randomizations allows us to show that the positive correlations at low phylogenetic distances are significantly higher than what expected by change. We confirmed the robustness of this empirical observation by changing the metric to quantify abundance pairwise correlations, obtaining in all cases similar decaying correlation patterns (see “Supplementary Information” (SI), Fig. S2-S3).

At a more quantitative level, the reported decay of the correlation function is well captured on average by a stretched-exponential function [44]:

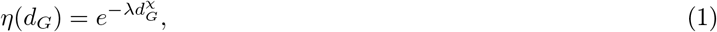

where *χ* ~ 1/3, as shown in Fig. 1(b). The value of *χ* and the goodness of fit of the functional form of Eq. 1 have both a small degree of variation across biomes. The best fits of the exponent *χ* across all the considered biomes —always in the range 0.2 − 0.4— are reported in sec.S3.B of the SI. Interestingly, the decay of the correlation function is slower than exponential: the logarithm of the correlations exhibits an algebraic (power-law) scaling with the phylogenetic distance *d_G_* as explicitly illustrated in the inset of Fig. 1(b).

In order to scrutinize whether the observed pattern is consistent across the phylogenetic tree, we repeated the same type of analyses at the coarser level of taxa, comparing correlations within and between taxonomic orders. Fig. S11 shows that species from different taxa (i.e. at large phylogenetic distances) tend to have, on average, vanishing correlations, while the averaged correlations within the same taxa decay from positive to zero with phylogenetic distance, recovering the pattern in Fig. 1 in a consistent way in the vast majority of the observed taxa (see Fig. S8-S10). Small deviations to this general pattern appear to be due to specific taxa. In the SI we explore the case of soils where a couple of orders are the main drivers of the observed departure from the macroecological law (see Fig. S9).

These results suggest that the observed correlation pattern showing a stretched-exponential decay with phylogenetic distance is a universal one, not depending on the considered ecological context nor on a particular taxa. Whatever ecological forces are at the origin of such species-abundance correlations, they manifest themselves regularly and consistently across taxa and environments.

### Ecological forces in preference space: three alternative scenarios produces three alternative predictions

Which ecological forces are responsible for the described pattern of abundance correlations across communities? In general, ecological interactions in a network of species could *a priori* create both positive and negative correlations, which stem from both direct pairwise interactions and network effects.

To unravel these conflicting mechanisms, we consider a general population-dynamic model where species grow and may compete for resources in a fluctuating environment. To be more specific, we consider a set of *N* species whose growth is coupled with a set of *R population-dependent* factors (e.g., resources) as well as on *M* other *population-independent* factors (such as, typically, very abundant resources, temperature, pH, salinity, etc.). The key difference between these two sets, is that population-dependent factors are affected by population abundances through consumption and, thus, resource availability explicitly depends on population densities. This dependence (which can be derived from consumer-resource models, as shown in SI section S4.G) induces an effective competition. On the other hand, population-independent factors display fluctuations which are independent of population densities. Furthermore, we assume that all these *R* + *M* factors, both population-dependent and population-independent, are characterized by temporal fluctuations (independent across factors). Population-dependent factors can be interpreted as nutrients, which are depleted by consumption. Population-independent factors could be abiotic factors or nutrients whose dynamics is not governed by population densities. For instance, if populations are limited by other factors (e.g. other resources or phages) it could happen that some resources are depleted by non-biological processes (e.g., dilution) before they are affected by consumption. In this scenario, resource abundance is independent of population density.

We consider that the effect of the environment on a given species can be represented as a vector in an abstract preference space, representing its preferences for the *R* available population-dependent resources and growth responses to the *M* population-independent factors. Thus, the set of resource preferences characterizing each given species *i* is represented by a vector ***b**^i^* in an abstract *R*-dimensional space of population-dependent factors and by another vector ***a**^i^* in a *M*-dimensional space of population-independent factors (see Fig. 2(a) for a pictorial illustration). The precise way in which these vectors are generated is described Methods and in Sec.S4.A in the SI, where diverse algorithms are designed for the cases of high and low-dimensional preference spaces, respectively.

**FIG. 2:**
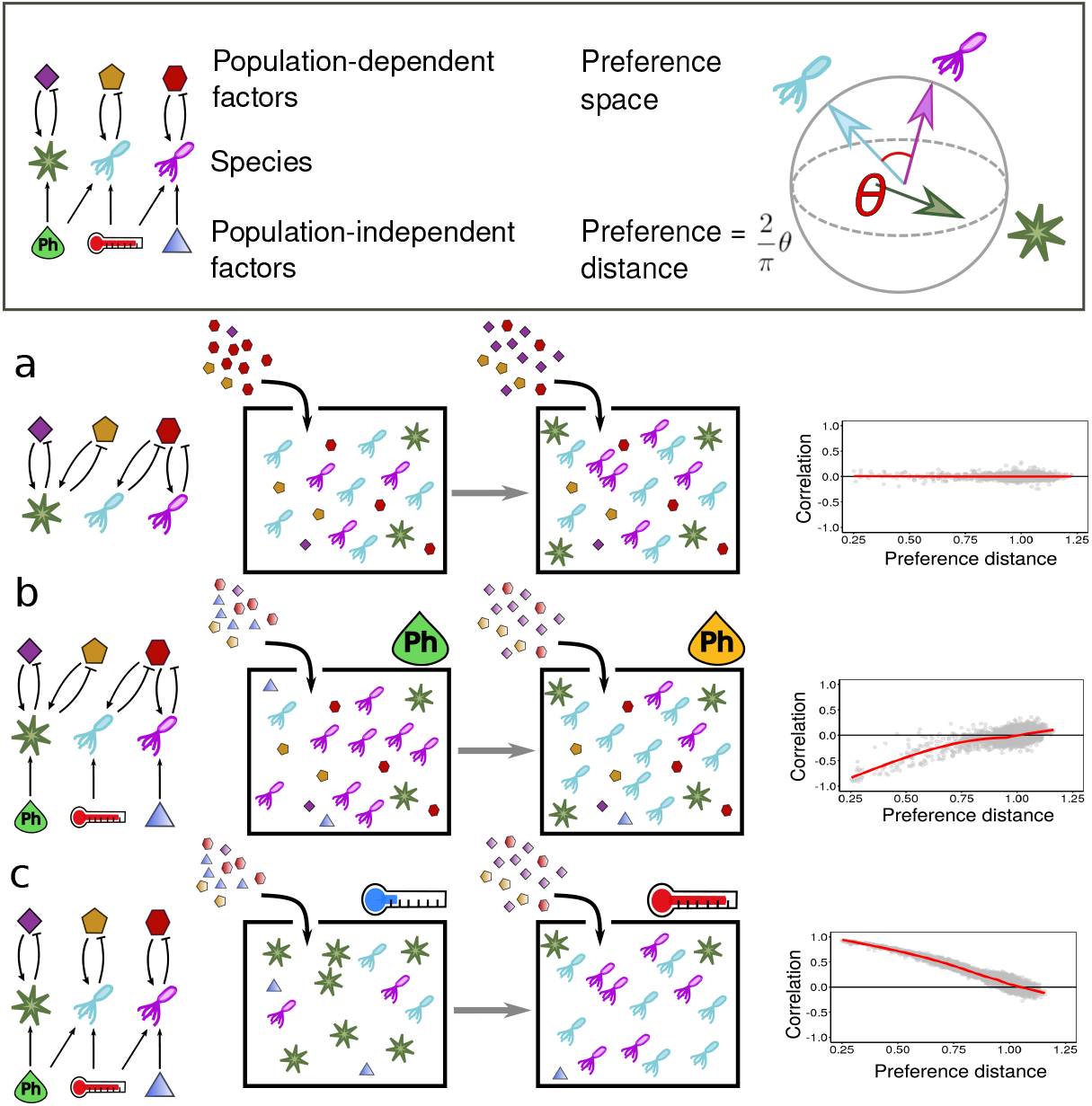
**(Top) Sketch of the elements of the model**. *Left*: Bacterial species depend upon both *population-dependent factors* such as resource abundances (polygons) and *population-independent factors*,e.g. abiotic variables like temperature, pH, light intensity, etc., and very abundant resources (shady polygons). The normal arrows stand for species preferences, while the inhibition arrows symbolize the feedback between population-dependent factors and populations. *Right*: Species preferences are represented as radial vectors in a sphere: positive/negative projections represent positive/negative influence on growth. The (pairwise) preference distance is quantified by the angle between vectors (multiplied by 2/*π*, see methods). The red and blue species are similar to each other while the green one is more different from them. **(Bottom) Schematic illustration for the three considered scenarios** (models A, B and C) of: (*left*) population-dependent and population-independent factor preferences; (*centre*) model dynamics, and (*right*) stationary correlations as a function of preference distance (with gray dots standing for simulation results and red lines for averages/theory). **(a) Shared fluctuating population-dependent resources**. When species are subjected to the combination of both forces, their effects cancel out leading to an “effective” neutral situation with no correlations. **(b) Shared population-dependent resources and non-overlapping fluctuating population-independent factors**. when two species sharing some resource preference experience an environmental fluctuation, one outcompetes the other, causing negative correlations, increasing monotonically to zero as similarity decreases. **(c) Shared fluctuating population-independent factors with fixed non-overlapping resources**. If two species share some preference for population-independent factors, but not for resources, the follow in a similar way environmental fluctuations, causing positive correlations which decrease with preference distance.

The *per-capita* growth rate of each species is influenced by population-dependent and population-independent factors, both weighted by the corresponding preference vector leading to the following general model

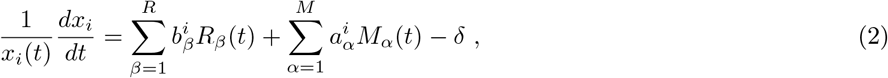

where *x_i_*(*t*) is the abundance of species *i* at time t, *R_β_*(*t*) is the value of population-dependent factor *β* at time *t*, *M_α_*(*t*) is the value of the population-independent factor *α*, and *δ* is a constant death rate. We assume the population-independent factors to be subject to stochastic fluctuations

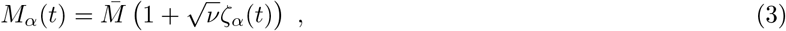

where 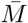 represents a baseline level, *ζ_α_*(*t*) is a (zero-mean unit-variance) Gaussian white noise and the parameter *ν* quantifies the strength of fluctuations. Similarly, the abundance *R_β_* of each population-dependent resource *β* fluctuates in time and is reduced by consumption, i.e.,

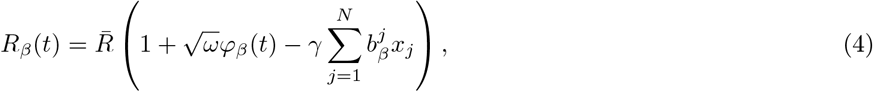

where 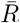 is a baseline level, *γ* the consumption timescale, *φ_β_*(*t*) a (zero-mean unit-variance) Gaussian white noise, and the parameter *ω* quantifies the amplitude of fluctuations. In sec.S4.G. of the SI, we sketch how to derive Eq. (2) from a more standard consumer-resource model [45, 46]. Note also that in the case of considering abiotic factors, such as temperature or pH, as population-independent factors, they could affect the growth rate in a multiplicative way, as often assumed in metabolic theory [47]. In Sec.S4.F of the SI we show that such a multiplicative variant of the model, that we name Model D, is qualitatively equivalent to Eq.(2) but mathematically much more involved.

The model defined by Eq.(2) can be effectively written as a generalized Lotka-Volterra model with species competition and fluctuating growth rates:

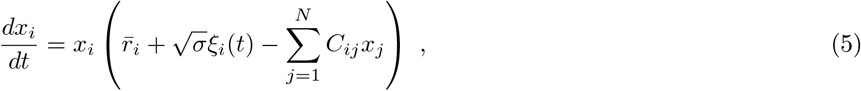

where 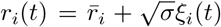 is a time-dependent and fluctuating growth rate with mean value 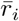; the entries of the competition matrix, *C_ij_*, are determined by the overlap in population-dependent factor preferences of species *i* and *j*

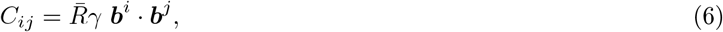

and, finally, the noise covariances are 〈*ξ_i_*(*t*)*ξ_j_*(*t*′)〉 = *ρ_ij_δ*(*t* − *t*′) with

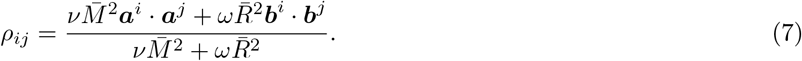

The explicit dependence of 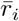, *σ* and *ξ_i_* on the original parameters is described in the Methods section. Observe that growth-rate correlations depend both on population-dependent and population-independent factors (***b*** and ***a***) while the effective competition is mediated only by shared population-dependent ones (***b***).

Thus, both types of coupling terms in the effective Lotka-Volterra population dynamics —i.e. the elements of the species competition matrix and those of the growth-rate fluctuation covariance matrix— crucially depend on the set of species similarities in preference space. However, depending on the relative strength of both types of couplings, as well as on the distribution of preferences vectors, one can define three different models, depending on which are the dominating ecological forces (see Fig. 2 a/b/c):

A. *Shared fluctuating population-dependent resources*. If population-independent fluctuations are negligible (i.e. *ν* = 0), species interactions are determined by a combination of the effect of competition (encoded in the entries *C_ij_*) and resource-abundance fluctuations (encoded in the entries *ρ_ij_*), which are both proportional to the species resource-preference overlap: ***b**^i^* · ***b**^j^*.
B. *Shared population-dependent resources and non-overlapping fluctuating population-independent factors*. If resource fluctuations are negligible (i.e., *ω* = 0) and population-independent factors preferences are all orthogonal to each other, species experience independent growth rate fluctuations (*ρ_ij_* = *δ_ij_*) and competition for fixed resources (through the coupling matrix *C_ij_*).
C. *Shared fluctuating population-independent factors with fixed non-overlapping resources*. If population-dependent factor preferences are all orthogonal to each other, there are, essentially, no shared resources. In this case, species experience correlated growth rate fluctuations and no inter-specific competition 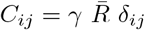. We refer to this force as “*environmental filtering*”.

Observe that more general and complex models involving both correlated population-independent factors and resource fluctuations, as well as combinations of the previous limiting cases, could also be constructed, but, for the sake of clarity, here we focus of these three (simplest) ones.

Using extensive numerical simulations (see Methods) we investigate the relationship between pairwise abundance correlations and preference similarities for the three models. In particular, one can define a preference distance, *d_P_* (where the sub-index *P* stands for “preference”) proportional to the angle between preference vectors for each pair of species (with *d_P_* = 0 for coinciding vectors and *d_P_* = 1 for orthogonal ones). In model (A) and (B) such a distance is calculated over the resource preference ***b***, while the vectors of population-independent factors preferences ***a*** need to be considered in model (C).

As illustrated in Figure 2(a/b/c) the three models give raise to three qualitatively distinct patterns of correlation as a function of preference distance *d_P_* : (A) Shared fluctuating population-dependent resources induce an effective *neutral* behavior, with nearly vanishing correlations across the spectrum of pairwise preference distances. (B) Shared resources and non-overlapping fluctuating population-independent factors produce *negative* correlations at small distances that increase to near-zero values in a monotonic way. (C) Shared fluctuating population-independent factors with fixed non-overlapping resources leads to correlations that decay from *positive* to vanishing values with distance.

### Environmental filtering reproduces the correlation decay with distance

In order to make a more quantitative comparison between the previous results and the empirically-determined universal pattern of decaying correlations, it is necessary to specify the relation between the preference distances *d_P,ij_*—on which the models rely— and the empirically-determined phylogenetic similarity of actual species, as quantified by their genetic distance *d_G,ij_*. For this purpose, it seems natural to assume that *d_P_* and *d_G_* are positively correlated, i.e., that phylogenetically close species typically have more similar preferences than distant ones. Under this assumption, the overall trend of the decay in Fig. 2 implies that environmental filtering is the process responsible for the empirically-observed decay of correlations (Fig. 1). Competition for constant and/or shared fluctuating resources can instead be discarded as the leading mechanism on the basis of the empirically observed pattern. This does not imply that competition is not present, but rather that it does not generate a signal detectable at a phylogenetic level within the present level of resolution.

To make further quantitative progress in the connection between the previous mechanistic modelling approaches—in particular, model C or “environmental filtering”— and available phylogenetic data, one needs to define a more precise mapping between preference similarity in the model and empirically-determined phylogenetic distance, i.e. to characterize the functional dependence between *d_P_* on *d_G_*, using information on pairwise correlations.

The previous task is not straightforward: species are coupled to each other within a network of interactions, so that pairs of species cannot be simply analyzed one at the time and, on the other hand, the full set of coupled non-linear equations is intractable. Fortunately, however, as explicitly shown in the Methods sect., one can make further progress by explicitly mapping model C into a *correlated stochastic logistic model* (CSLM) :

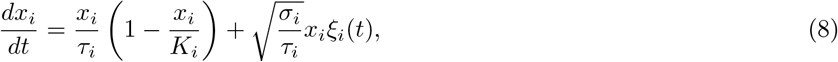

where 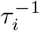 is the growth rate, *K_i_* an effective carrying capacity, *σ_i_* the amplitude of environmental fluctuations, and *ξ_i_*(*t*) is a Gaussian white noise, with correlations proportional to the preference distance,

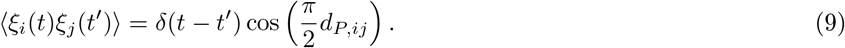

For the sake of simplicity, in the derivation (see Methods), we assumed that the preference space has a large dimensionality, i.e. *M* ⨠ 1, but this can be shown not to limit the generality of the forthcoming results (see SI, Sec. S4.E for more details).

This mapping is particularly illuminating as the resulting CSLM extends the standard stochastic logistic model (SLM) [25], as it includes correlated growth-rate fluctuations that stem from shared environmental fluctuating resources and that induce non-trivial species correlations. Moreover, it is important to stress that —if species-abundances trajectories are observed individually— there are no statistical differences between the CSLM and the standard SLM. This implies that the CSLM also reproduces (as the SLM does) the three macroecological patterns put forward in [25–27] (see Methods). Thus, the CLSM constitutes an improvement of existing modelling approaches to microbial macroecological laws.

A crucial advantage of Eq.(8) (together with and Eq.(9)) with respect to the generalized Lotka-Volterra equation is that is can be treated analytically to obtain a mathematical expression linking pairwise species-abundance correlations with their preference distance, *d_P,ij_* (see Methods). The resulting analytical relationship can be exploited to estimate the preference distance matrix from empirical correlation data, thus allowing us to establish the desired relation between preference distance *d_P_* and phylogenetic distance *d_G_* for every pair of species (see Methods):

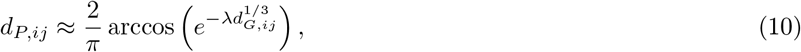

where *λ* is a constant. Observe, that Eq.(10) is highly non-linear, implying that, as the phylogenetic distance grows, preference distances rapidly saturate to values close to 1; in other words, even phylogenetically similar species tend to have a large preference dissimilarity (i.e. their preference vectors tend to be orthogonal to each other).

By implementing the relation given by Eq (10) in the definition of noise correlations Eq. (9), we obtain a version of the CSLM, directly relating ecological processes and phylogeny, which allows us to relate the species-abundance pairwise correlations to their empirically measured genetic similarity, *d_G,ij_*. Actually, given that the macroecological pattern we intend to reproduce is for the averaged correlation at a given (binarized) phylogenetic distance, we dropped the sub-index *ij* in Eq.(10) and use it as a relation between averages (see Methods, Eq (46)). In particular, by combining Eq. (46) with Eq. (37), one obtains exactly Eq. (1), i.e. the empirically observed relation between correlation and phylogenetic distance (see Methods).

Fig. 3 shows that, for the particular case of the human gut microbiome, a computational simulation of the final version of the model captures quite well the averaged decay of pairwise correlations with phylogenetic distance and that the analytical predictions describe accurately such an averaged behavior.

**FIG. 3:**
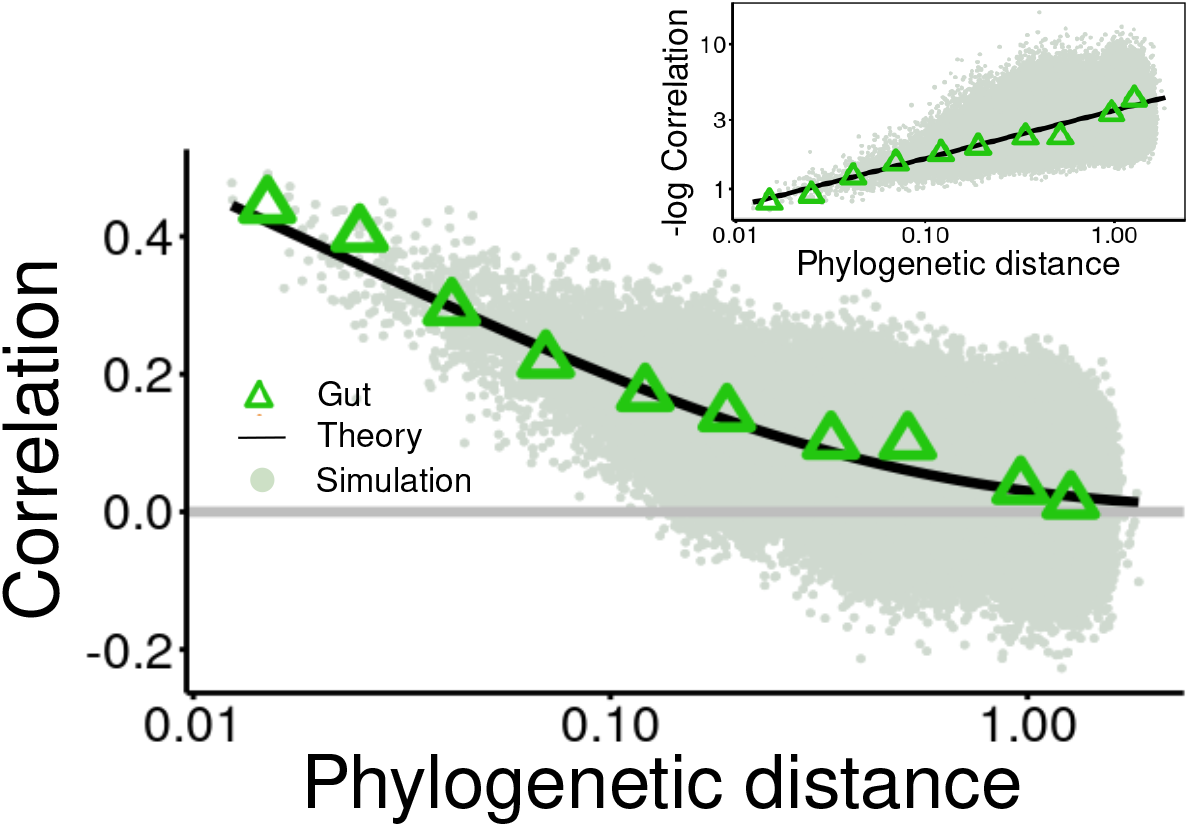
The model with environmental filtering reproduces the empirical law. Correlation values are plotted as a function of the phylogenetic distance both for the gut microbiome data (green triangles for each binarized value) and the simulated computational model (green clouds of points). The analytical expression, Eq.(1) with *λ* = 3.5 is also plotted (black line). Simulations of the model have been performed, using *N* = 300 species and considering as an input the empirical phylogenetic distance matrix of the gut microbiome, randomly sampling from it the *N* species. Inset: −log Correlations as a function of phylogenetic distance in double-logarithmic scale, empirically and from the mode, same data as the main figure. For more simulations details see Methods

### The macroecological law holds for temporal (longitudinal) data

One important prediction of Eq. (8) is that the decay of abundance correlations with phylogenetic distance is caused by shared temporal fluctuations. In order to further test the predictions of Eq. (8), we consider longitudinal data from the human microbiome. In particular, we analyzed three human body sites (gut, oral cavity, and hand palms) of two hosts [43]. From these data, we calculate the correlation of species abundance fluctuation *η_ij_* as above, but now averaging over time, rather than across individuals (see Fig. 4(a)). In particular, Figure 4(b) illustrates —for the specific case of the human gut— that the macroecological law of decaying correlation holds also for such temporal data, and that delayed correlations rapidly decay to zero. In particular, the correlations as a function of phylogenetic distance decay on average as a stretched exponential with an exponent close to 1/3, as observed in cross-sectional data.

**FIG. 4:**
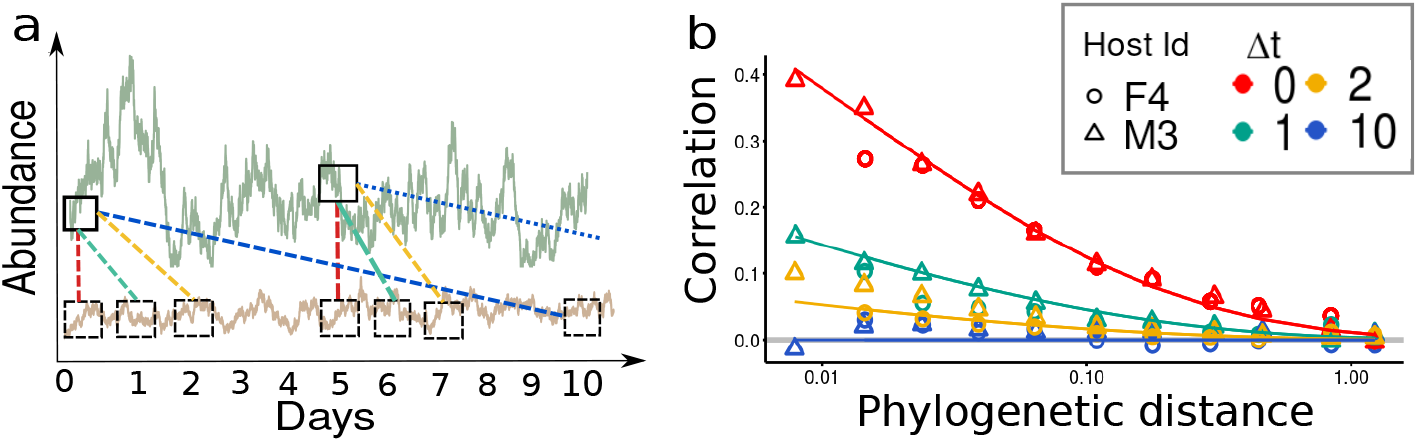
**(a) Sketch of the time-dependent (longitudinal) correlation data analyses**. Typical time-series for two species (green and brown, respectively) along 10 days. The dashed lines illustrate how equal-time (red) and 1, 2, 10 days delayed correlations (green, yellow and blue respectively) are computed, see Methods for more details. **(b) Macroecological law for temporal data**. Equal time (red), one-day delay (light blue), two-days (yellow) and ten-day delay (blue) symbols represent correlations as a function of the discretized phylogenetic distance (logarithmic scale) for the gut microbiomes of two different hosts (labelled with circles (F4) and triangles (M3), respectively). Solid lines stand for the prediction from the CSLM, Eq. (45), averaged over hosts, with timescale parameter *τ_i_* = 1, for *i* = 1, .., *N* and *λ* = 4.5.

To further test the CSLM model in its ability to reproduce time-dependent features of species correlations, we also computed delayed pairwise correlations, *η_ij_*(Δ*t*) defined as the correlation between the abundance fluctuations of species *i* at time *t* with the abundance fluctuations of species *j* at time *t*+Δ*t* (see Methods and Fig. 4 (a) for a graphical illustration). Let us remark that, in principle, the value of such a delayed correlation is, in general, not trivially linked to the correlation computed at the same time, as it depends of the specific properties of the dynamics giving rise to species inter-dependencies. Remarkably, as shown in Figure 4(b) the CSLM with no additional modification quantitatively reproduces also the temporal delayed correlations for different values of the delay (see SI Sec. S3.E for additional details and analyses) only by setting the growth time scale *τ_i_* = 1 for all species.

## Discussion

We have considered both cross-sectional (across communities) and longitudinal (across time) empirical data for the species abundances in microbial communities from many different environments and studied their species-abundance pairwise correlations as a function of pairwise phylogenetic distance, revealing the emergence of an universal macroecological law. This empirical law states in quantitative terms that the average correlation function decays from positive to null values as the phylogenetic distance (or dissimilarity) increases, following a stretched-exponential decay function.

We explored the possible ecological forces shaping species correlations from a theoretical standpoint. In particular, by scrutinizing different ecological models, each one implementing a diverse set of ecological forces between species, we found that the universal correlation pattern cannot possibly be reproduced by competition or exclusion principles. Instead, temporal environmental filtering —i.e. the presence of correlated noise stemming from shared fluctuating factors— as modeled by a correlated stochastic-logistic model (CSLM), explains quantitatively empirical data. Furthermore, time-dependent (delayed) correlations in longitudinal data are also well reproduced by the model.

This novel ecological pattern gives a quantification at the level of phylogenetic signals detectable in taxa-taxa abundance correlation. The pattern, as also shown in Fig. S5, S6, and S7, does not recapitulate the full range of correlations observed in natural communities. In this context, our work complements the research aiming at inferring ecological interactions from correlations, by showing how phylogenetic similarity can be used to disentangle the effects of environmental fluctuations and interactions (such as, e.g., competition).

These results are based on multiple assumptions and their limitations give opportunities for extensions of the current work. First, at a theoretical level, the CSLM reproduces the average correlation at each discrete phylogenetic distance, but not the full distribution around such a mean value (see Fig. S29 in SI). This is because, to be able to connect genetic and preference similarities, we enforced a “mean-field” type of relationship, Eq. (10), neglecting variability across pairs of species in the phenotypic-distance-to-preference-distance mapping. On the other hand, in figure S5 of the SI, we show that the variance of the distribution of the empirically-measured pairwise correlations within each distance bin seems to follow a weak decaying power-law pattern with phylogenetic distance, with a diverse decaying exponent characteristic for each analyzed biome. Possibly, these patterns could be used to generate the preference vectors of the model in a more general way, allowing for more variability. Empirical data are not informative enough at the moment to proceed in this direction, and further analyses are required.

It is also important to stress that the origin of the power law behavior and its exponent value close to a value 1/3 in the universal pattern of correlations (i.e., Eq.(1)) remains unexplained. This type of scaling could be determined by the scale-invariant, i.e. fractal, structure of phylogenetic trees [48–51], but further investigations, beyond the scope of the present work, are needed to shed light onto this empirical finding. Furthermore, it is known that a vast class of competitive models can lead to species clustering in trait space [52, 53]. Even if such models produce an “oscillating” pattern of positive and negative correlation, and hence are not sufficient to explain the behavior here reported, their possible extension could be relevant for explaining the phylogenetic distance distribution observed in data (see Fig. S1 in SI).

Although environmental filtering has been found to dominate the pattern of species-abundance correlations, the above-mentioned variability could be the result of the complex interplay of other ecological forces. To identify which further forces are relevant and to discriminate their effects it will be important to analyze time-dependent data in a more detailed way as well as to analyze differences in carrying capacities, and correlations between different hosts [27]. Furthermore, an exhaustive analysis of the variations of the correlation pattern across environments and phyla is also needed. Interestingly, Fig. S8-S10 in the SI show that some phyla (e.g. bacteroidetes) follow robustly the pattern, while some others, such as actinobacteria, exhibit wild fluctuations. Indeed, the non-monotonic deviation in the soil biome around distance 0.1 seems to be caused by the actinobacteria phylum and, in particular, by the actinomycetales and gaiellales orders (see Fig. S9). Hence, we leave for future work the promising study of deviations across taxa, that could reveal more information on additional interactions responsible for the observed residual correlations. Another relevant caveat is that our analyses here are limited to the taxonomic resolution of OTUs, clustering together individuals with more than 97% similarity. Recent results suggest that ecological dynamics starts to decouple at much finer phylogenetic resolutions [54]. Moreover, strains seem to still obey the three macroecological laws of variation and diversity valid at species level [55]. These results leave open the question of how ecological forces shape the variation of community composition at finer phylogenetic scales.

On the other hand, from a complementary viewpoint, we analyzed the behavior of correlations at the coarse-grained resolution of phyla. In particular, Fig. S11, in the SI illustrates that by considering just inter-phyla correlations one cannot observe the stretched exponential decay, that is determined by intra-phyla OTU pairs. Analogously, by extending our analyses to finer phylogenetic resolutions it could be possible to reveal the nature of intra-specific interactions, eventually elucidating the emergence of competition as a key player in determining correlations. Actually, in our view, one should not fix a characteristic taxonomic resolution to have a complete description of complex communities, but, instead, start from individuals (or functional units) and progressively cluster them together at larger and larger coarse-grained scales (i.e. moving across observational scales as customarily done in physics using “renormalization group” tools in statistical physics [56, 57]) as different ecological forces may shape communities at diverse resolution levels [58].

### Data and Code

All the data sets analyzed in this work have been previously published and were obtained from the EBI Metagenomics database [43]. Previous publications of some of us have reported on the details of the experiments and the corresponding statistical analyses [25]. In order to test the robustness of the macroecological laws and the modeling framework presented in this work, we considered 7 data sets that differ not only on the considered biome, but also on the sequencing techniques and the pipelines used for data processing which underscores the consistency of our results. Data sets were selected to represent a wide set of biomes. We considered only data sets with at least 50 samples with more than 10^4^ reads. No data set was excluded *a posteriori*. The main code used for analysis is available here.

### Correlation analysis

In each community a, with *a* = 1, .., *M*, the count of the *i*-th species, with *i* = 1, …, *N*, is called 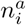 and only sufficiently abundant species are considered, i.e., 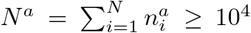. The relative abundance of species *i* in community *a* is calculated as:

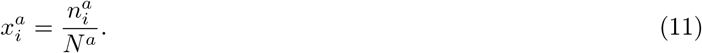

Community averages are defined as: 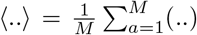, such that the mean and the variance of a species relative abundance are:

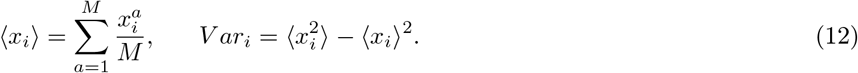

Another important quantity is the rank of species *i* in community *a*, 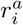, where the most abundant species has rank 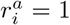, the second most abundant 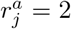, and so on. Using these ingredients, one can construct the following (five) different quantities, that gauge fluctuations in species abundance, or simply “*fluctuation quantifiers*”:

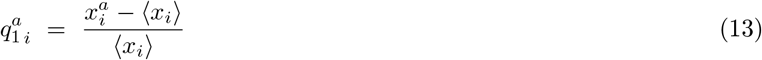

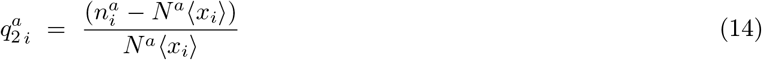

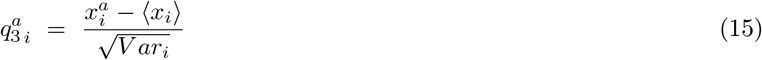

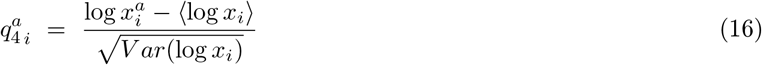

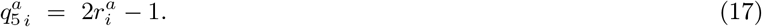

Similarly, one can estimate the correlation between species abundance fluctuations by using any of these quantifiers:

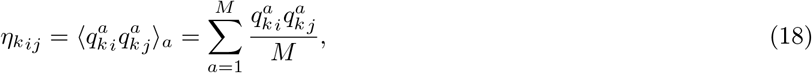

for *k* = 1, 2, …5. Finally, one can average over all pairs of species with a distance falling within a certain “bin” of phylogenetic distance.

#### Temporal analyses

The analysis of temporal (longitudinal) data is analogous to that for cross-sectional data in the preceding section, but instead of studying fluctuations and correlations between different communities, one considers a single community a data along a time-series (e.g. samples from different days of the time series, *t* = 1, .., *T*). All the quantities are defined as above but replacing the community average by a time average 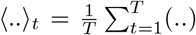. In particular, the equal-time pairwise correlations are defined by:

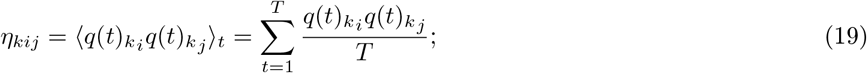

for species *i* an *j*. Similarly, the Δ*t* delayed correlation is:

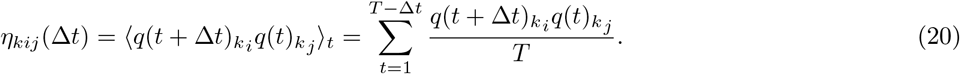

### Models in preference space and evolutionary algorithm

In the preference space model, each single species is represented by a *R*-dimensional population-dependent factor (resource) preference vector ***b*** and a *M*-dimensional population-independent factor one ***a***. Without loss of generality, environmental factors are assumed to be equivalent and, in order to fix a tradeoff, the preference vectors of all species are assumed to have the same squared module, fixed to 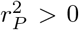, so that they can be characterized by a point in a *R*-dimensional sphere of radius *r_P_*, i.e.: 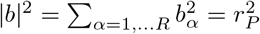 (respectively on a *M*-dimensional sphere with same radius in population-independent space).

Using the explicit expressions for the dynamics of population-dependent and population-independent factors, the general model as defined by Eq.(2), can be rewritten as the generalized Lotka-Volterra equation, Eq.(5), with deterministic growth rate and interaction matrix given by

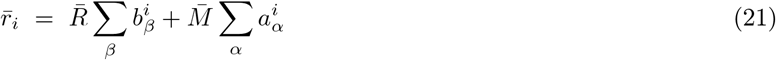

and

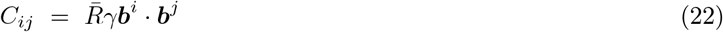

respectively, and with an effective zero-mean Gaussian noise:

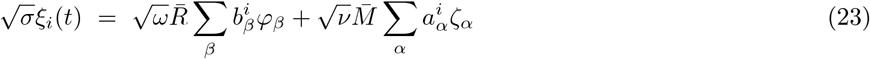

with a covariance matrix given by Eq.(7) and a noise amplitude 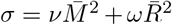; see Sec. S4 in the SI for the detailed derivation.

In all the variants of the model considered here (A, B and C) only one set of preference vector is needed (either population-dependent for models A and B, or population-independent for C). Thus, one can quantify the preference similarity or, simply, the “*preference distance*” between species *i* and *j*, as the cosine distance between their relevant preference vectors (for simplicity, in the following, we restrict the notation to model C for which population-independent preferences are relevant). The preference distance is defined as:

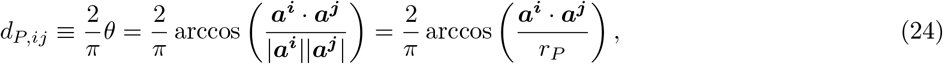

where the sub-index *P* stands either for “preference”.

One can generate the set of *M* preference vectors ***a*** by sampling their component from a Gaussian with mean *m/M* (*m* small and positive) and standard deviation 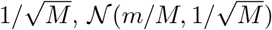, such that the radius is constant and close to unity for large values of *M* :

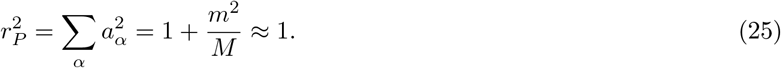

However, as a consequence of the central limit theorem, for sufficiently large numbers of environmental factors, *M*, the random vectors ***a^i^*** tend to be orthogonal to each other, i.e., *d_P,ij_* ≈ 1 ∀*i, j*), hindering the possibility of generating similar species by simple random sampling. In order to circumvent this difficulty, we devised a simple evolutionary algorithm, such that, starting from an initial random distribution of vectors 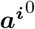 —and implementing and evolutionary branching process— generates as an outcome a set of vectors ***a^i^*** which are distributed across a broad range of possible cosine-distance values. The algorithm includes the following steps:

1. Sample at random two species *i, j*, *j* dies and *i* reproduces, making a copy (labeled *j*) of itself with some variation.
2. The preference vectors of the new species are obtained from the old one with some variation:

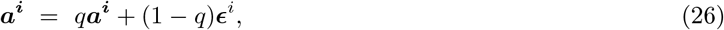

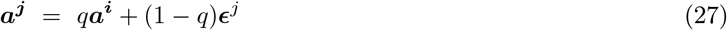

where the parameter *q* ∈ [0, 1] is the fidelity of reproduction and *ϵ*^*i,j*^ are vectors sampled from 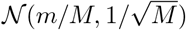 (note that the resulting vectors are kept within the sphere).
3. Iterate Z times.

By considering a sufficiently large number of iterations *Z* and a value *q* = 0.9, the population develops a pool of similar individuals, with small pairwise distances, which was absent in the initial condition and covers, even if in an heterogeneous way, all the spectrum of possible distances (see Fig. S19-S20 in the supplementary material). On the other hand, if the dimension *M* of population-independent factors cannot be considered large, e.g. in the presence of just a few abiotic factors such as temperature or pH, we have devised an alternative algorithm that can produce a long-tail distance distribution even when *N* ⋙ *M* (see the SI Sec. S4.A.2. for more details).In any case, the previous evolutionary algorithms are just efficient procedures used to generate communities with a broad distribution of phylogenetic distances.

### Correlated stochastic logistic model

#### Derivation

The CSLM is obtained from Eq. (2) in the case where each species consumes only one resource with baseline 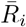 at rate *γ_i_* and this resource is not consumed by any other species (model C). In particular, by taking the limit *M* ⨠ 1, one can easily find Eq.(8) with the following definitions of the involved parameters:

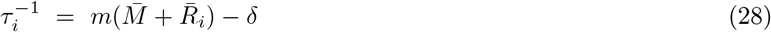

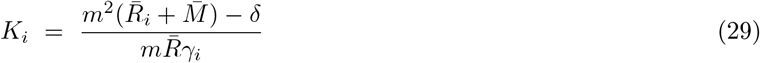

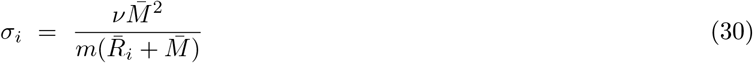

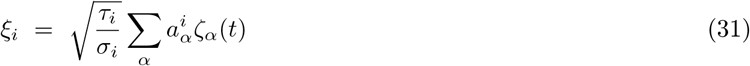

The new environmental noise *ξ_i_* is still Gaussian, because it is the weighted sum of Gaussian variables, with moments:

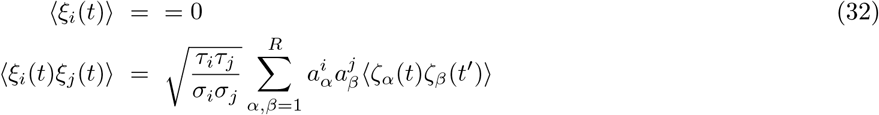

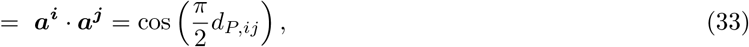

where we have used the parameter definition Eq.(28), the normalization condition |***a^i^***|^2^ = 1 and the definition of preference distance. In the case of *N* ⨠ *M* one can still write the CSLM but., in order to keep it equivalent to the Model C also the modulus of preference vectors need to be taken into consideration, see Sec. S4.E in the SI.

#### Macroecological laws and marginal properties

The CSLM, in the discretization scheme, has a Gamma stationary *marginal* distribution [25, 59]:

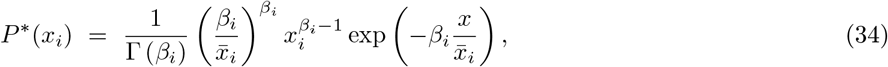

where the average abundance 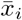 and the squared inverse coefficient of variation *β_i_* read:

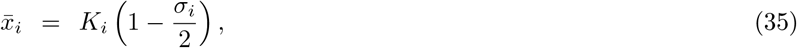

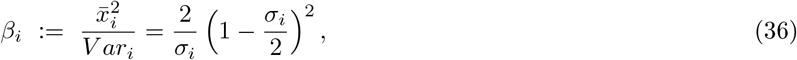

respectively, coinciding with the ones obtained for the standard SLM [25]. Hence, the CSLM is able to reproduce the three macroecological laws for diversity and fluctuation, namely:

1. The stationary marginal distribution of species abundances is a Gamma distribution.
2. By fixing *σ_i_* = *σ*, for all species the Taylor law relating the mean and variances across species is recovered.
3. The mean abundances are distributed as a log-normal just by imposing that the *K_i_*’s are log-normally distributed too.

#### Correlations

The joint probability cannot be calculated analytically for the CSLM, and hence an expression for the pairwise correlation functions cannot be derived in an exact way. Nevertheless, one can rely on a linear-noise approximation around the fixed point (see SI. Sec. S4.E.1 for details) leading to:

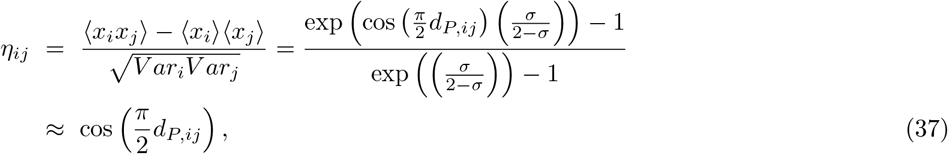

which is the expression employed in the main text to relate correlations with preference distances.

On the other hand, the Langevin equation of the model can be solved exactly, leading to:

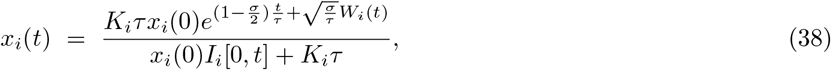

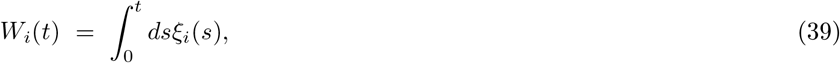

where *I_i_*[0, *t*] the integral of the associated “geometric Brownian motion” [60]:

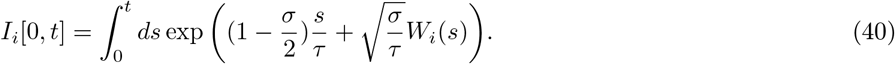

The exact integral Eq.(38) can be used to understand the effect of delays on pairwise correlations. Indeed, assuming that the system has reached the stationary state for sufficiently large times (*t* → ∞), one can compute heuristically the relation between the abundance at time *t* and at a later time *t* + Δ*t*:

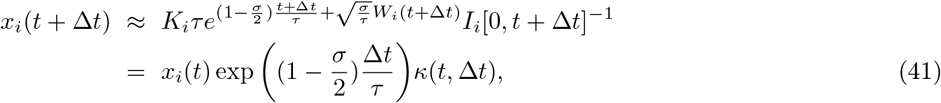

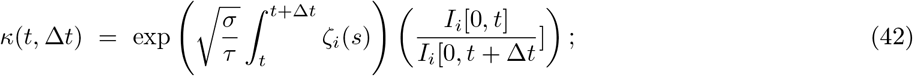

the function *κ* in the limit *t* ⨠ Δ*t* converges to 1, so that

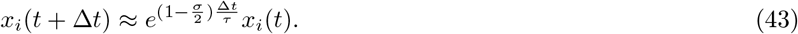

Plugging this expression into the Pearson coefficient formula one readily obtains:

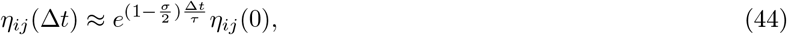

that combined with the linear approximation, Eq.(37), leads to:

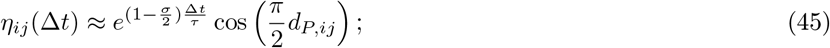

see SI. Sec. S4.E.2 for more details.

#### Inferring preference distances from data

To tune the CSLM to reproduce the observed empirical pattern it is necessary to infer the relation between preference and phylogenetic distances. Note that the empirical pattern we aim at reproducing is between average correlation and averaged phylogenetic distance within each bin, i.e., it suffices to find a relation between the (average) distance *d_P_* and *d_G_* (in other words: we are not interested in the full probability distribution of correlations in one bin, but just on its mean value).

The preference distance of species can be now explicitly calculated by inverting the formula for the correlation Eq. (37) separately for each species pair and by taking averages over the couples within each bin of phylogenetic distance:

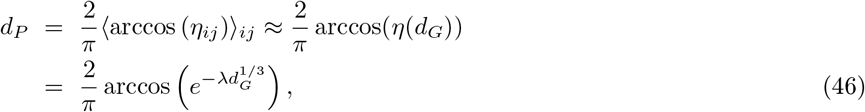

where the variance of *η_ij_* within each bin of phylogenetic distance has been neglected, i.e. a so-called *“mean-field approximation”*. A plot and a discussion of Eq (46) can be found in the SI, Sec. S4.H Thanks to Eq.(46) it is possible to generate a preference-distance matrix, and hence the matrix of noise pairwise correlations from phylogenetic data:

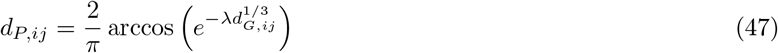

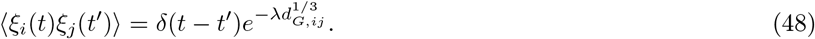

Clearly, this simple version of the CSLM cannot reproduce correlation variability as a function of phylogenetic similarity (see Discussion for possible extensions).

### Computational simulations

The different models in preference space, Eq.(5) as well as the CSLM have been simulated in the Itô discretization scheme using the Milshtein algorithm [61] In Fig. 2, gray points stand for the Pearson’s correlation coefficients at the stationary state for 10 realizations with *N* = 200 species and *M* = *R* = 300; the averages are obtained over 10^3^ samples at stationarity, at time separated by *δ_t_* = 10. Red lines are obtained by averaging the correlation over pairs. In each simulation the initial populations are sampled from a Gaussian distribution *N* (0.5, 0.01); other parameters are *N* = 200, *R* = *M* = 300, *m* = 0.1, 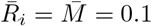, *γ_i_* = 1, *ν_α_* = *ω_α_* = 0.1, *q* = 0.9, *Z* = 50*N*, *t_fin_* = 10^4^.

In Fig. 3 dark-green points stand for the Pearson’s correlation coefficient at the stationary state of 10 realizations with *N* = 300 species, the averages are over 10^3^ abundances sampled during the stationary time series every *δ_t_* = 10*τ*. In each realization, we use the phylogenetic distances of N species sampled at random from the phylogenetic distance matrix of a random community of the considered biome to construct the species noises correlation, Eq.(47). The model parameters are set to reproduce the species marginal properties and delayed correlations, following the prescriptions from the previous section, in Methods, and in [25]. Carrying capacities are generated log-normally by taking the exponential of random variables sampled from a Gaussian distribution 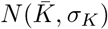, *τ_i_* = *τ* and *σ_i_* = *σ* for *i* = 1, .., *N*. Parameter values: *τ* = 1, 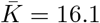, *σ_K_* = 3.8, *σ* = 1.42, *λ* = 3.5, *t_f_* = 10^4^.

MAM and MS acknowledge the Spanish Ministry and Agencia Estatal de investigación (AEI) through Project of I+D+i Ref. PID2020-113681GB-I00, financed by MICIN/AEI/10.13039/501100011033 and FEDER “A way to make Europe", as well as the Universidad de Granada and Consejería de Conocimiento, Investigación Universidad, Junta de Andalucía and European Regional Development Fund, Project B-FQM-366-UGR20 for financial support. We also thank W. Shoemaker for a careful reading of the manuscript and J. Iranzo, J. Cuesta, J. Camacho Mateu, R. Rubio de Casas, and L. Seoane for valuable discussions.

## References

[1] Whitman, W. B., Coleman, D. C. & Wiebe, W. J. Prokaryotes: The unseen majority. Proceedings of the National Academy of Sciences 95, 6578–6583 (1998). URL https://www.pnas.org/content/95/12/6578. https://www.pnas.org/content/95/12/6578.full.pdf.

[2] Mandal, S. et al. Analysis of composition of microbiomes: a novel method for studying microbial composition. Microbial ecology in health and disease 26, 27663 (2015).

[3] Frentz, Z., Kuehn, S. & Leibler, S. Strongly deterministic population dynamics in closed microbial communities. Phys. Rev. X 5, 041014 (2015). URL https://link.aps.org/doi/10.1103/PhysRevX.5.041014.

[4] Ratzke, B. J., Christoph & Gore, J. Strength of species interactions determines biodiversity and stability in microbial communities. Nature Ecology Evolution 4, 376–383 (2020). URL https://doi.org/10.1038/s41559-020-1099-4.

[5] Gralka, M., Szabo, R., Stocker, R. & Cordero, O. X. Trophic interactions and the drivers of microbial community assembly. Current Biology 30, R1176–R1188 (2020). URL https://www.sciencedirect.com/science/article/pii/S0960982220311611.

[6] Friedman, J., Higgins, L. M. & Gore, J. Community structure follows simple assembly rules in microbial microcosms. Nature Ecology &Evolution 1 (2017).

[7] Szabo, R. E. et al. Ecological stochasticity and phage induction diversify bacterioplankton communities at the microscale. bioRxiv (2021). URL https://www.biorxiv.org/content/early/2021/09/27/2021.09.27.461956. https://www.biorxiv.org/content/early/2021/09/27/2021.09.27.461956.full.pdf.

[8] Hu, J., Amor, D. R., Barbier, M., Bunin, G. & Gore, J. Emergent phases of ecological diversity and dynamics mapped in microcosms. Science 378, 85–89 (2022).

[9] Goldford, J. E. et al. Emergent simplicity in microbial community assembly. Science 361, 469–474 (2018).

[10] Kehe, J. et al. Positive interactions are common among culturable bacteria. Science Advances 7, eabi7159 (2021).

[11] Ley, R. E., Lozupone, C. A., Hamady, M., Knight, R. & Gordon, J. I. Worlds within worlds: evolution of the vertebrate gut microbiota. Nature Reviews Microbiology 6, 776–788 (2008).

[12] Thompson, L. et al. A communal catalogue reveals earth’s multiscale microbial diversity. Nature 551, 457–463 (2017).

[13] Lozupone, C. A. & Knight, R. Global patterns in bacterial diversity. Proceedings of the National Academy of Sciences 104, 11436–11440 (2007). URL https://www.pnas.org/content/104/27/11436. https://www.pnas.org/content/104/27/11436.full.pdf.

[14] Arumugam, M. et al. Enterotypes of the human gut microbiome: [plus]corrigendum [plus]addendum. Nature 473, 174–180 (2011).

[15] Grieneisen, L. et al. Gut microbiome heritability is nearly universal but environmentally contingent. Science 373, 181–186 (2021). URL https://www.science.org/doi/abs/10.1126/science.aba5483. https://www.science.org/doi/pdf/10.1126/science.aba5483.

[16] Martiny, J. B. H., Jones, S. E., Lennon, J. T. & Martiny, A. C. Microbiomes in light of traits: A phylogenetic perspective. Science 350, aac9323 (2015).

[17] Prosser, J. I. et al. The role of ecological theory in microbial ecology. Nature Reviews Microbiology 5, 384–392 (2007).

[18] Marquet, P. A. et al. On Theory in Ecology. BioScience 64, 701–710 (2014). URL https://doi.org/10.1093/biosci/biu098. https://academic.oup.com/bioscience/article-pdf/64/8/701/14094079/biu098.pdf.

[19] Gilbert, J. A. & Dupont, C. L. Microbial metagenomics: Beyond the genome. Annual Review of Marine Science 3, 347–371 (2011). URL https://doi.org/10.1146/annurev-marine-120709-142811. PMID: 21329209, https://doi.org/10.1146/annurev-marine-120709-142811.

[20] Johnson, J. et al. Evaluation of 16s rrna gene sequencing for species and strain-level microbiome analysis. Nature Communications 10 (2019).

[21] Brown, J. H. et al. Macroecology (University of Chicago Press, 1995).

[22] Shade, A. et al. Macroecology to unite all life, large and small. Trends in Ecology Evolution 33, 731–744 (2018). URL https://www.sciencedirect.com/science/article/pii/S0169534718301861.

[23] Shoemaker, W. R., Locey, K. J. & Lennon, J. T. A macroecological theory of microbial biodiversity. Nature Ecology &Evolution 1 (2017).

[24] Ji, B. W., Sheth, R. U., Dixit, P. D., Tchourine, K. & Vitkup, D. Macroecological dynamics of gut microbiota. Nature microbiology 5, 768–775 (2020).

[25] Grilli, J. Macroecological laws describe variation and diversity in microbial communities. Nature communications 11, 1–11 (2020).

[26] Descheemaeker, L. & de Buyl, S. Stochastic logistic models reproduce experimental time series of microbial communities. eLife 9, e55650 (2020). URL https://doi.org/10.7554/eLife.55650.

[27] Zaoli, S. & Grilli, J. A macroecological description of alternative stable states reproduces intra- and inter-host variability of gut microbiome. Science Advances 7, eabj2882 (2021).

[28] Ho, P.-Y., Good, B. H. & Huang, K. C. Competition for fluctuating resources reproduces statistics of species abundance over time across wide-ranging microbiotas. Elife 11, e75168 (2022).

[29] O’Dwyer, J. P. & Chisholm, R. A mean field model for competition: from neutral ecology to the red queen. Ecology letters 17, 961–969 (2014).

[30] Grilli, J., Barabás, G., Michalska-Smith, M. J. & Allesina, S. Higher-order interactions stabilize dynamics in competitive network models. Nature 548, 210–213 (2017).

[31] Wootton, J. T. Indirect effects in complex ecosystems: recent progress and future challenges. Journal of Sea Research 48, 157–172 (2002).

[32] Webb, C. O., Ackerly, D. D., McPeek, M. A. & Donoghue, M. J. Phylogenies and community ecology. Annual review of ecology and systematics 33, 475–505 (2002).

[33] HilleRisLambers, J., Adler, P. B., Harpole, W. S., Levine, J. M. & Mayfield, M. M. Rethinking community assembly through the lens of coexistence theory. Annual review of ecology, evolution, and systematics 43, 227–248 (2012).

[34] Cadotte, M. W. & Tucker, C. M. Should environmental filtering be abandoned? Trends in ecology & evolution 32, 429–437 (2017).

[35] Emerson, B. C. & Gillespie, R. G. Phylogenetic analysis of community assembly and structure over space and time. Trends in ecology & evolution 23, 619–630 (2008).

[36] Poulin, R., Krasnov, B. R., Pilosof, S. & Thieltges, D. W. Phylogeny determines the role of helminth parasites in intertidal food webs. Journal of Animal Ecology 1265–1275 (2013).

[37] Krasnov, B. R. et al. Co-occurrence and phylogenetic distance in communities of mammalian ectoparasites: limiting similarity versus environmental filtering. Oikos 123, 63–70 (2014).

[38] Pérez-Valera, E. et al. Fire modifies the phylogenetic structure of soil bacterial co-occurrence networks. Environmental Microbiology 19, 317–327 (2017).

[39] Cavender-Bares, J., Kozak, K. H., Fine, P. V. & Kembel, S. W. The merging of community ecology and phylogenetic biology. Ecology letters 12, 693–715 (2009).

[40] Gaulke, C. A. et al. Ecophylogenetics clarifies the evolutionary association between mammals and their gut microbiota. MBio 9, e01348–18 (2018).

[41] Jeraldo, P. et al. Quantification of the relative roles of niche and neutral processes in structuring gastrointestinal microbiomes. Proceedings of the National Academy of Sciences 109, 9692–9698 (2012).

[42] O’Dwyer, J. P., Kembel, S. W. & Green, J. L. Phylogenetic diversity theory sheds light on the structure of microbial communities. PLoS computational biology 8, e1002832 (2012).

[43] Mitchell, A. et al. Ebi metagenomics in 2017: Enriching the analysis of microbial communities, from sequence reads to assemblies. Nucleic acids research 46 (2017).

[44] Laherrére, J. & Sornette, D. Theoretical microbial ecology without species. The European Physical Journal B 2, 525â539 (1998). URL https://doi.org/10.1007/s100510050276.

[45] MacArthur, R. Species packing and competitive equilibrium for many species. Theoretical Population Biology 1, 1–11 (1970). URL https://www.sciencedirect.com/science/article/pii/0040580970900390.

[46] Posfai, A., Taillefumier, T. & Wingreen, N. S. Metabolic trade-offs promote diversity in a model ecosystem. Phys. Rev. Lett. 118, 028103 (2017). URL https://link.aps.org/doi/10.1103/PhysRevLett.118.028103.

[47] Brown, J. H., Gillooly, J. F., Allen, A. P., Savage, V. M. & West, G. B. Toward a metabolic theory of ecology. Ecology 85, 1771–1789 (2004). URL https://esajournals.onlinelibrary.wiley.com/doi/abs/10.1890/03-9000. https://esajournals.onlinelibrary.wiley.com/doi/pdf/10.1890/03-9000.

[48] Burlando, B. The fractal dimension of taxonomic systems. Journal of Theoretical Biology 146, 99–114 (1990). URL https://www.sciencedirect.com/science/article/pii/S0022519305800463.

[49] Hernandez-Garcia, E., Tugrul, M., Herrada, E., EguÃluz, V. & Klemm, K. Simple models for scaling in phylogenetic trees. International Journal of Bifurcation and Chaos 10, 805–811 (2010).

[50] Xue, C., Liu, Z. & Goldenfeld, N. Scale-invariant topology and bursty branching of evolutionary trees emerge from niche construction. Proceedings of the National Academy of Sciences 117, 7879–7887 (2020). URL https://www.pnas.org/content/117/14/7879. https://www.pnas.org/content/117/14/7879.full.pdf.

[51] OâDwyer, J. P., Kembel, S. W. & Sharpton, T. J. Backbones of evolutionary history test biodiversity theory for microbes. Proceedings of the National Academy of Sciences 112, 8356–8361 (2015).

[52] Scheffer, M. & Van Nes, E. H. Self-organized similarity, the evolutionary emergence of groups of similar species. Proceedings of the National Academy of Sciences 103, 6230–6235 (2006).

[53] Ramos, F., López, C., Hernández-García, E. & Munoz, M. A. Crystallization and melting of bacteria colonies and brownian bugs. Physical Review E 77, 021102 (2008).

[54] Goyal, A., Bittleston, L. S., Leventhal, G. E., Lu, L. & Cordero, O. X. Interactions between strains govern the eco-evolutionary dynamics of microbial communities. bioRxiv (2021). URL https://www.biorxiv.org/content/early/2021/01/04/2021.01.04.425224. https://www.biorxiv.org/content/early/2021/01/04/2021.01.04.425224.full.pdf.

[55] Wolff, R., Shoemaker, W. & Garud, N. Ecological stability emerges at the level of strains in the human gut microbiome. bioRxiv (2021). URL https://www.biorxiv.org/content/early/2021/10/01/2021.09.30.462616. https://www.biorxiv.org/content/early/2021/10/01/2021.09.30.462616.full.pdf.

[56] Wilson, K. G. Problems in physics with many scales of length. Scientific American 241, 158–179 (1979).

[57] Efrati, E., Wang, Z., Kolan, A. & Kadanoff, L. P. Real-space renormalization in statistical mechanics. Reviews of Modern Physics 86, 647 (2014).

[58] Tikhonov, M. Theoretical microbial ecology without species. Phys. Rev. E 96, 032410 (2017). URL https://link.aps.org/doi/10.1103/PhysRevE.96.032410.

[59] Faust, K. et al. Signatures of ecological processes in microbial community time series. Microbiome 6, 1–13 (2018).

[60] Dufresne, D. The integral of geometric brownian motion. Advances in Applied Probability 33, 223–241 (2001).

[61] Toral, R. & Colet, P. Stochastic Numerical Methods: An Introduction for Students and Scientists (Wiley-Vch, 2014).

