## Supplementary material for "Environmental fluctuations explain the universal decay of species-abundance correlations with phylogenetic distance": main.pdf

### Environmental fluctuations explain the universal decay of species-abundance correlation with phylogenetic distance-Supplementary Information

Matteo Sireci,[1,*] Miguel A. Muñoz,[1,†] and Jacopo Grilli[2,‡]

[1]*Departamento de Electromagnetismo y Física de la Materia e Instituto Carlos I de Física Teórica y Computacional. Universidad de Granada. E-18071, Granada, Spain*

[2]*Quantitative Life Sciences, The Abdus Salam International Centre for Theoretical Physics, 34151 Trieste, Italy*

Section S1 presents the datasets that have been used as well as the criteria considered in the empirical analyses. In section S2 we report on the methods used for constructing and analyzing phylogenetic trees. Section S3 presents the different methods employed to measure correlations (A) showing that in cross-sectional the decay is quite robust and that it disappears when the phylogenetic tree is randomized. Furthermore, we also report on the fitting procedure used for the stretched exponential decay (B), its variability in and across biomes (C), and its analysis at the larger scale of phyla (D). The same analyses are repeated for longitudinal/temporal data together with the study of delayed correlations (E). Finally, Section S4 is devoted to the study of various mechanistic models. We show how to produce a coherent distance preference distribution (A), study various candidate models introduced in the main text both in preference (B,C,D,E) and phylogenetic space (H). In addition, we report on a purely abiotic version (F) model and on how to derive these general class of models from a consumer-resource framework (G). Supplementary figures and tables are also included.

##### Contents

**S1. Datasets** 3

**S2. Phylogenetic Analysis** 3

**S3. Correlation analysis** 4

A. Cross-sectional data 4

B. Fit to stretched exponential 8

C. Variability 8

D. Taxonomic Analysis 9

E. Time-series 16

**S4. Models** 20

A. Evolutionary algorithms to generate a wide distribution of preference distances. 22

*Electronic address: ``

†Electronic address: ``

‡Electronic address: ``

1. Algorithm 1: high-dimensional preference space 22
2. Algorithm 2: low-dimensional preference space 22
B. Model 0: Fluctuating and non overlapping factors. 23
C. Model A:Shared fluctuating population-dependent factors. 24
D. Model B: Shared resources and non-overlapping fluctuating population-independent factors. 25
E. Model C: Shared fluctuating population-independent factors with fixed non-overlapping resources. 26
1. Linear approximation around the fixed point 28
2. Temporal behavior 33
F. Model D: Purely abiotic fluctuations 34
G. Model derivation from a consumer-resource framework 35
H. CSLM in phylogenetic space 37

**References** 38

#### S1. DATASETS

All the datasets analyzed in this work were obtained from EBI Metagenomics (now Magnify) and have been previously published [4]. Raw data were processed under different version of EBI Metagenomics pipelines [4] . The consistency of results across studies and pipelines strongly support the robustness and generality of the conclusions here. Supplementary Table S1 reports the references to the original works, description of the Magnify pipeline, and other relevant informations about each dataset. Note that the pipeline version 4.1 uses the algorithm SILVA [6] to assign an OTU classification. Observe also that here we use the term "species" to refer to OTUs, defined accordingly to the methods referred above. Datasets were selected to represent a wide set of diverse biomes: "gut" (human gut), "oral" (human mouth), "lake" and "river" (aquatic ecosystems), "activated sludge", "soil" and "glacier". We considered only datasets with at least 50 samples with more than $10^4$ reads. No dataset was excluded a-posteriori.

| Biome | Type | EBI ID | Magnify ID | Pipeline Version | NCBI ID | Reference | # Samples | [Range Tot, # Reads $N_s$] |
|---|---|---|---|---|---|---|---|---|
| Gut | c | SRP056641 | MGYS00001056 | 2.0 | PRJNA275349 | [1] | 66 | [13842, 102971] |
| Oral | c | SRP056641 | MGYS00001056 | 2.0 | PRJNA275349 | [1] | 62 | [10006, 138172] |
| River | c | ERP012927 | MGYS00001669 | 3.0 | PRJEB11530 | [5] | 188 | [76042, 352675] |
| Lake | c | ERP012927 | MGYS00001669 | 3.0 | PRJEB11530 | [5] | 198 | [57408, 350877] |
| Soil | c | SRP052295 | MGYS00000905 | 2.0 | PRJNA272333 | - | 112 | [11352, 58219] |
| Sludge | c | ERP009143 | MGYS00001064 | 2.0 | PRJEB8105 | [5] | 575 | [22255, 912713] |
| Glacier | c | ERP017997 | MGYS00001292 | 3.0 | PRJEB16145 | [1] | 30 | [79765, 1104214] |
| Gut F4 | l | ERP021896 | MGYS00002184 | 4.1 | PRJEB19825 | [2] | 131 | [21008, 51986] |
| Gut M3 | l | ERP021896 | MGYS00002184 | 4.1 | PRJEB19825 | [2] | 334 | [15047, 58463] |
| Oral F4 | l | ERP021896 | MGYS00002184 | 4.1 | PRJEB19825 | [2] | 135 | [5683, 12651] |
| Oral M3 | l | ERP021896 | MGYS00002184 | 4.1 | PRJEB19825 | [2] | 331 | [1052, 2'3567] |
| Skin L-palm F4 | l | ERP021896 | MGYS00002184 | 4.1 | PRJEB19825 | [2] | 134 | [12298, 34607] |
| Skin L-palm M3 | l | ERP021896 | MGYS00002184 | 4.1 | PRJEB19825 | [2] | 365 | [144, 48475] |
| Skin R-palm M3 | l | ERP021896 | MGYS00002184 | 4.1 | PRJEB19825 | [2] | 358 | [135, 91953] |

Supplementary Table S1: Description and references for the datasets used in this work. In column âTypeâ, c refers to cross-sectional (across communities) and l to longitudinal (across time).

#### S2. PHYLOGENETIC ANALYSIS

All the statistical analysis has been carried out with the tools of the *phyloseq* R library [3]. The phylogenetic tree of each community is obtained by removing the absent species from the three with all possible species. Then, the distance $d_{G,ij}$ between species i and j is calculated as the cophenetic distance [3]. Furthermore, each distance is categorized in one of $n_b = 17$, possible logarithmic bins, where the bin $b$ to which a diven distance is assigned is given by

$$b = int\left(\frac{\log(d) - \min(\log(d))}{[\max(\log(d)) - \min(\log(d))]n_b}\right). \tag{S1}$$

Note that the min and max are calculated for each community/sample independently.

In each bin, the mean distance is calculated by averaging over all the species pairs with a distance within such a bin:

$$d_G(b) = \langle d_{G,ij} \rangle_b = \sum_{i,j \in b}^{N_b} \frac{d_{G,ij}}{N_b}, \tag{S2}$$

where $N_b$ is the number of species couples within bin $b$. Figure S1 shows the histograms of the phylogenetic average distance $d_G$ for the set of considered biomes. Even if the distributions show quantitatively different patterns for each biome, they share the fact that they exhibit a maximum at large distance (slightly below 1) and a monotonous decay to zero with a long tail.

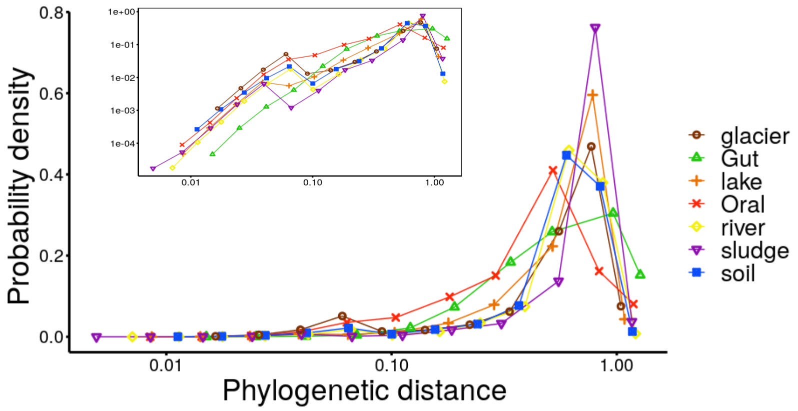


Supplementary Figure S1: **Phylogenetic distance distribution (log scale)** Histogram of the average phylogenetic distances $d_G$for different biomes (colors). The frequency in each bin (points) is calculated using the discretization given by Eq.(S1). We only considered bins with at least $10^3$ pairs of species. Interestingly, for all the considered biomes there is a peak around 1 and slowly-decaying left tail. The inset shows the same data but in log-log scale.

#### S3. CORRELATION ANALYSIS

##### A. Cross-sectional data

In each community $a$, with $a = 1, ..., M$, the count of species $i$, with $i = 1, ..., N$, is represented by $n_i^a$; only sufficiently abundant samples are considered, i.e. the total reads for a community a must be larger than $10^4$: $N^a = \sum_{i=1}^N n_i^a \geq 10^4$.

The relative abundance of species $i$ in community $a$ is calculated as:

$$x_i^a = \frac{n_i^a}{N^a}, \tag{S3}$$

and the average over communities is defined as:

$$\langle .. \rangle = \frac{1}{M} \sum_{a=1}^{M} (..), \tag{S4}$$

such that one can calculate the mean and the variance of a species relative abundance:

$$\langle x_i \rangle = \sum_{a=1}^{M} \frac{x_i^a}{M}, \tag{S5}$$

$$Var_i = \langle x_i^2 \rangle - \langle x_i \rangle^2. \tag{S6}$$

Another important observable is the rank of species $i$ in community $a$, $r_i^a$: the most abundant species has rank $r_i^a = 1$, the second most abundant $r_j^a = 2$, and so on.

Using these ingredients one can construct a set of different observables quantifying species-abundance fluctuations:

$$q_{1\,i}^{a} = \frac{x_i^a - \langle x_i \rangle}{\langle x_i \rangle}, \tag{S7}$$

$$q_{2\,i}^{a} = \frac{n_i^a - N^a \langle x_i \rangle}{N^a \langle x_i \rangle}, \tag{S8}$$

$$q_{3\,i}^{a} = \frac{x_i^a - \langle x_i \rangle}{\sqrt{Var_i}}, \tag{S9}$$

$$q_{4\,i}^{a} = \frac{\log x_i^a - \langle \log x_i \rangle}{\sqrt{Var(\log x_i)}}, \tag{S10}$$

$$q_{5\,i}^{a} = 2r_i^a - 1. \tag{S11}$$

By multiplying a couple corresponding to different species $i$ and $j$, of the same-type observables, one can estimate pairwise correlations of species-abundance fluctuations:

$$\eta_{k\,ij} = \langle q_{k\,i}^{a} q_{k\,j}^{a} \rangle_a = \sum_{a=1}^{N} \frac{q_{k\,i}^{a} q_{k\,j}^{a}}{N}, \tag{S12}$$

with $k = 1, 2, 3, 4$ or 5. Finally, by averaging over all possible pairs of species with a mutual distance within a certain bin $b$, one can compute the averaged correlation in abundance fluctuations as a function of phylogenetic distance:

$$\eta_k(d_G) = \langle \eta_{k,ij} \rangle_{d_{G,ij} \in b} = \sum_{i,j=1, d_{P,ij} \in b}^{N} \frac{\eta_{k,ij}}{N_b}, \tag{S13}$$

where $N_b$ is the number of species couples in bin $b$.

To disentangle actual effects of phylogeny from possible spurious ones, we compared the measured correlations with those emerging from two alternative null models.

A) The first null model consists in calculating $\eta_k$ on a randomized tree, preserving its high-order structure, meaning preserving the transitivity relation, i.e. if $|a - b| < d$ and $|b - c| < d$ also $|a - c| < d$.

B) In the second null model one calculates correlations using a randomization of phylogenetic distances between species, i .e. the distance between a couple is exchanged randomly with another one.

These two alternative null models are equivalent for abundant samples. Figures S2-S3 show that the decay of abundance-fluctuation correlations with phylogenetic distance is qualitatively independent of the chosen observable, eq.(S7), and that the null models show constant, almost vanishing, correlations. These results strongly support that the observed decay results from the actual structure of the phylogenetic tree.

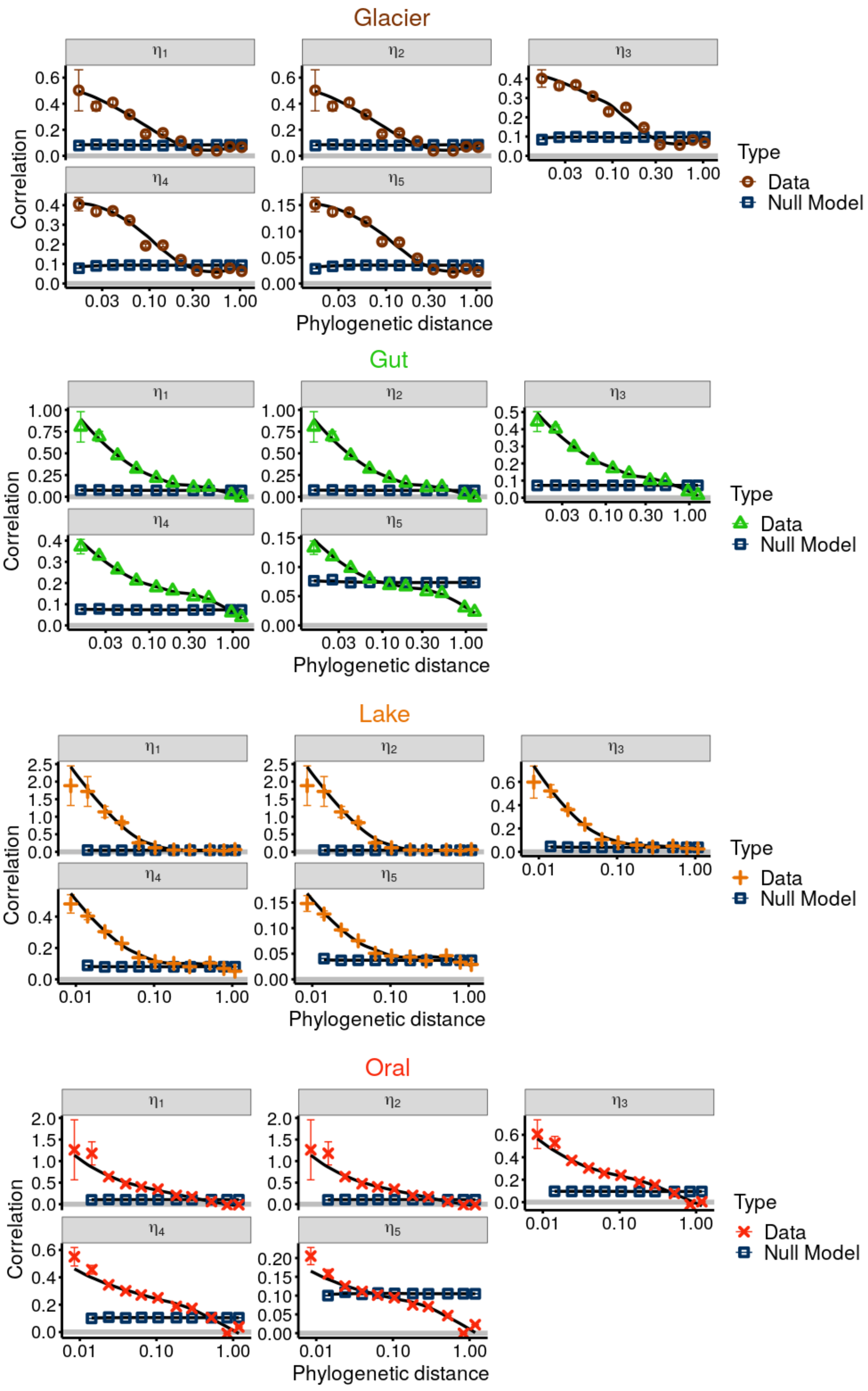


Supplementary Figure S2: **Correlation decay as a function of phylogenetic distance for different observables ($\eta$'s) in four different biomes, namely: glacier, gut, lake and oral.** Colored points stand for empirical correlation data, blue squares for the null model and the black line for the average of points weighted by the number of couples in each bin. Bars represent standard errors in each bin.

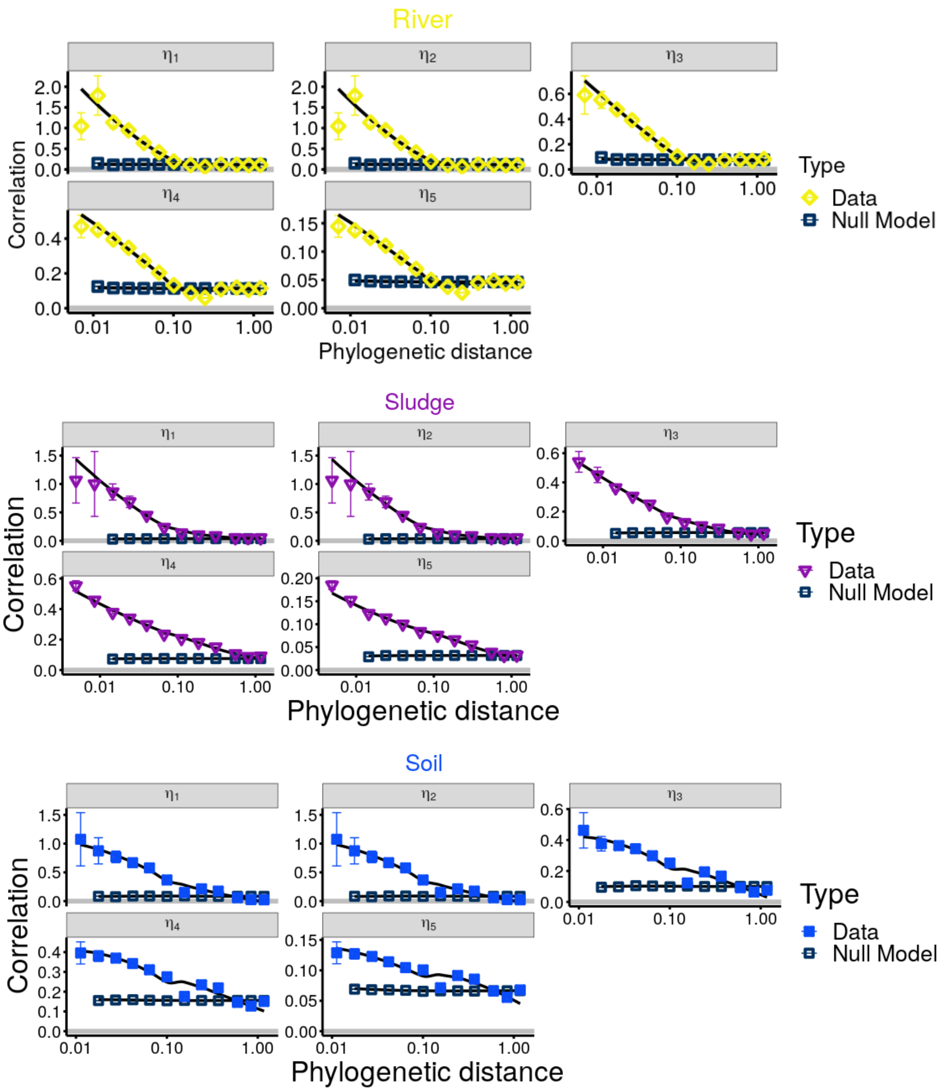


Supplementary Figure S3: **Correlation decay as a function of phylogenetic distance for different observables in three different biomes: river, sludge and soil**. As in the previous figure but for a different set of biomes.

#### B. Fit to stretched exponential

Here, we present the fitting procedure used to obtain the stretched exponential decay. For each biome, we fitted the $\eta_3$ estimator with a stretched exponential function of the form:

$$\eta = e^{-\lambda d_G^{\chi}}, \tag{S14}$$

by conducting a linear fit between $log(-log(\eta))$ and $\log(d_G)$, i.e. :

$$\log\left(-\log\eta\right) = \log\lambda + \chi\log\left(d_G\right). \tag{S15}$$

The different points are weighted by the number of couples within the bin. In Fig.S4 and in Table S2 we present the best parameters $\lambda$ and $\chi$, including errors, and the $R^2$ coefficient of determination for cross-sectional data and plot the resulting fitted curves together with data (plots).

As it can be seen, the vast majority of biomes follow a stretched exponential with excellent approximation, i.e. $R^2 \geq 0.9$, except for the lake and river ones where $R^2$ is smaller, but still larger than 0.8. Furthermore, there is a consistent variability of both the exponent $\chi$ and the scale parameter $\lambda$. Moreover, by considering all the points together independently of the biome, we still obtain a good fit to the stretched exponential curve ($R^2 = 0.86$), with the best fit parameters reported in the main text $\chi = 0.33 \sim 1/3, \lambda = 3.5$, see Fig. 1 in the main text. Let us note that while the fit for each biome is weighted by the number of couples in each bin, the total one is unweighted to not over represent any single biome.

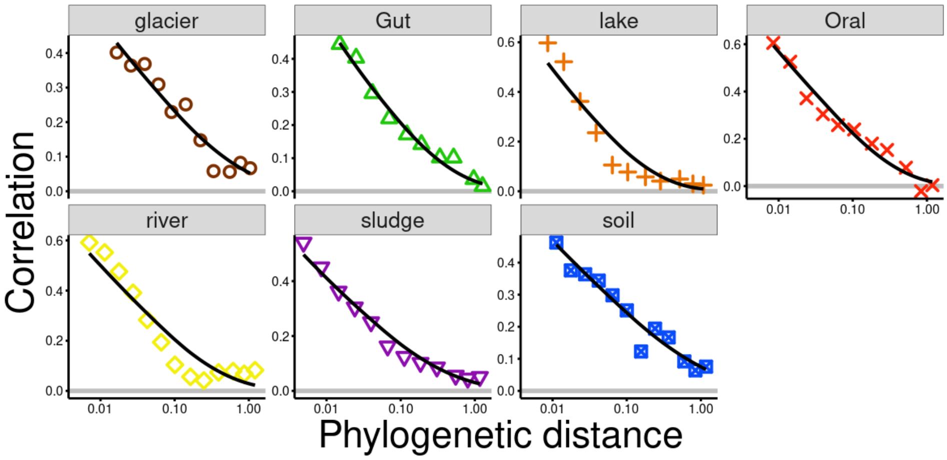


Supplementary Figure S4: **Stretched exponential fit for cross-sectional data.** Plots of the stretched exponential fit together with data. All the biomes follows with very good approximation a stretched exponential decay eq.(S14), with a poorer fit in the lake and river. Variability of both $\chi$ and $\lambda$ can be appreciated both in Table S2 and in the plot. Nevertheless, the "total" dataset still follows a stretched exponential with $\lambda = 3.41 sim 3.5$ and $\chi = 0.33 \sim 1/3$ with good approximation.

#### C. Variability

The results exposed in the preceding sections deal with the decay of the mean Pearson correlation (i.e. mediated in each bin) with phylogenetic distance. On the other hand, here we report on the properties of the full correlation

| Biome | $\chi$ | $\lambda$ | $R^2$ |
|---|---|---|---|
| Gut | $0.34 \pm 0.02$ | $3.42 \pm 0.16$ | 0.97 |
| Oral | $0.41 \pm 0.03$ | $3.9 \pm 0.40$ | 0.94 |
| River | $0.35 \pm 0.05$ | $3.52 \pm 0.47$ | 0.83 |
| Lake | $0.40 \pm 0.05$ | $4.62 \pm 0.62$ | 0.88 |
| Soil | $0.26 \pm 0.02$ | $2.6 \pm 0.14$ | 0.94 |
| Sludge | $0.29 \pm 0.02$ | $3.44 \pm 0.19$ | 0.96 |
| Glacier | $0.31 \pm 0.03$ | $3.00 \pm 0.22$ | 0.91 |
| Total | $0.33 \pm 0.01$ | $3.41 \pm 0.13$ | 0.86 |

Supplementary Table S2: Stretched-exponential fit parameters for each biome.

distribution within each bin. For the sake of simplicity, we consider the $\eta_3$ estimator only, but the results are similar for other quantifiers. Fig.S5 shows that the correlation variance exhibits a tendency to decay with the phylogenetic distance as a power law $d^{-\gamma}$, with the exponent $\gamma \in [1/6, 1/2]$ different for each biome. The power-law decay is not perfect as it shows some "oscillations", similar to what happens discrete fractals and could reveal a discrete scale invariance in the underlying tree. We leave for a future research the complete study of this pattern. For the sake of completeness, in Fig.S6 and Fig.S7, we plot the $\eta_3$ correlation coefficient distribution within each logarithmic bin, including at least $10^3$ couples, for all the considered biomes.

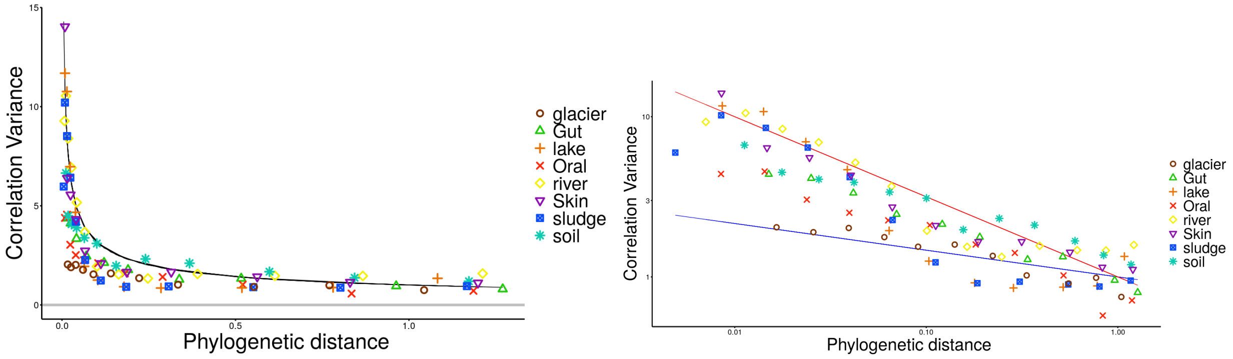


Supplementary Figure S5: **Quasi-universal pattern of correlation variance.** Points stand for the variance of the correlation distribution in each bin. Left: Correlation variance versus phylogenetic distance in natural scale, where the black line is a power-law fit $d^{-\gamma}$, with $\gamma = 1/2$. Right: the same data but represented in a log-log scale. There seems to be a tendency to behave like a power-law but each biome shows a different exponent $\gamma_i$ (red and blue lines stand for power-law fits with maximum and minimum possible exponents $\gamma_{max} = 1/2$, $\gamma_{min} = 1/6$) and there seem to be oscillations.

#### D. Taxonomic Analysis

To verify that the behavior of correlations is uniform across the phylogenetic tree, i.e. that particularly abundant phylum is not determining the decay, we study the correlation pattern at the larger taxonomic scale of phyla. First, we report on the correlations between species of the same phylum (*intra*-phylum correlation). By averaging separately the correlation between species of the same phylum, we obtain how the correlation of abundances fluctuations decays

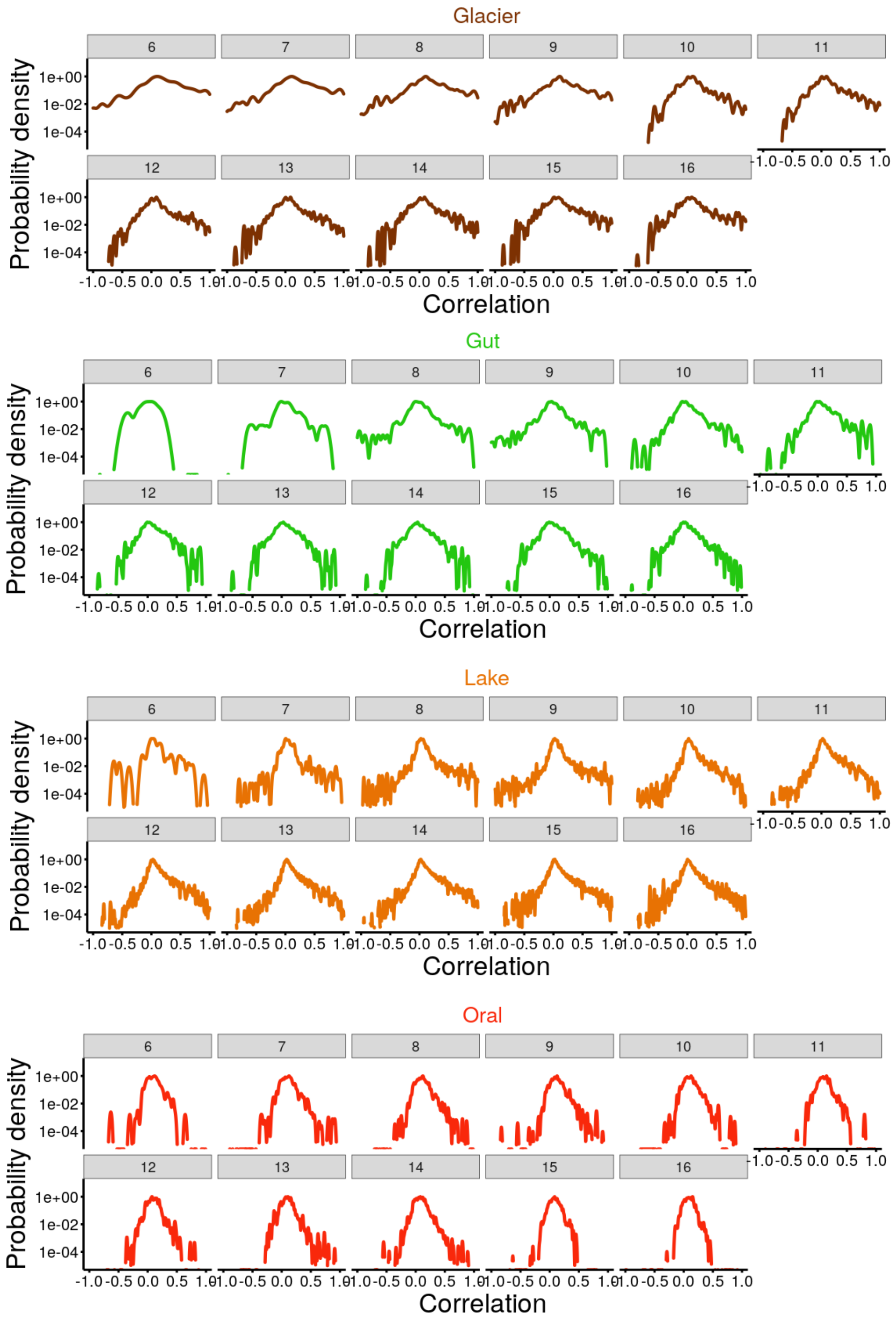


Supplementary Figure S6: $\eta_3$ correlation histogram in each bin (colored lines) in log-scale. Numbers on top of plots indicate the corresponding bin, from 1 to 17. Only bins with at least $10^3$ couples are considered.

with the phylogenetic distance for each single phylum. In Fig. S8 we plotted the intra-phylum correlation for each biome and verify that majority of them decay coherently with general pattern, while a few of them, e.g. actinobacteria, exhibit some deviations. Such taxa-dependent deviations could be responsible for deviations from the typical stretched exponential pattern for specific biomes. To test this claim, we consider as an example the soil biome, where a small

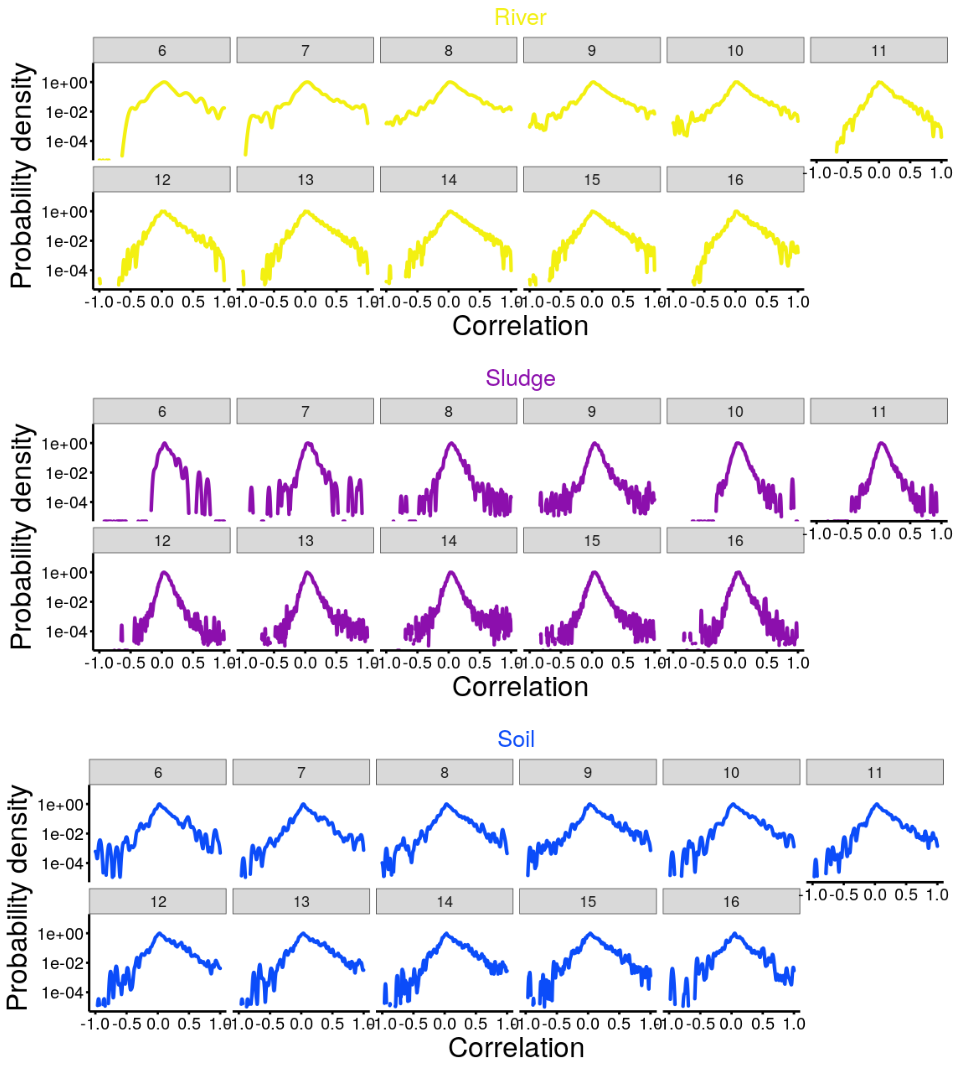


Supplementary Figure S7: $\eta_3$ correlation histogram in each bin (colored lines) in log-scale. Numbers on top of plots indicate the corresponding bin, from 1 to 17. Only bins with at least $10^3$ couples are considered.

non-monotonicity is present in the decay, as illustrated by a characteristic bump around $d = 0.1$ (cfr. with fig.S3). In Fig. S9) (Top), we show that such a deviation is probably due to the behavior of actinobacteria, that present small negative correlation at intermediate distances. Furthermore, in the bottom of Fig.S9), we show that, by zooming into the actinobacteria phylum and considering correlations at the finer level of orders, the negative correlations around distance $d_G = 0.1$ is caused by the actinomycetales and gaiellales. The other orders contribute mainly with positive correlation. These results seem to suggest that deviation from the typical pattern are driven by few bacterial orders. This analysis could be carried forward by going at even finer taxonomic resolution to identify the drivers of this deviation, with the goal of understanding which are the corresponding ecological traits which could produce such decreasing correlation. All these analyses open exciting routes for future research.

To complement the precedent analysis, in Fig.S10 we plotted the intra-phylum correlation (same data as before) for each phylum, representing with colors the different biomes. The decaying pattern is found consistently in the most abundant phylum, like acidobacteria, bacteriotedes, proteobacteria and firmicutes. On the other hand, in less abundant phyla correlations are still positive but not showing a universal behavior. We leave for a future work the study of how different order deviate from the general pattern and why.

Finally, in order to understand how the universal pattern changes at the larger scale of phylum and to discover how it emerges from the different phylum and what changes by coarse-graining in Fig.S11 we report both intra-phylum correlations (red points) and inter-phylum ones (i.e. between species of different phylum) separately. The correlation pattern for these taxonomic relation show two totally different behaviors. Inter-correlations concentrate symmetrically around zero and correspond to very large phylogenetic distances. On the other hand, as showed in the preceding figures, intra-taxonomic correlations decay from positive value to zero with with distance. By averaging over different phyla the stretched-exponential pattern (black line) emerges both in each single biome and in the total case.

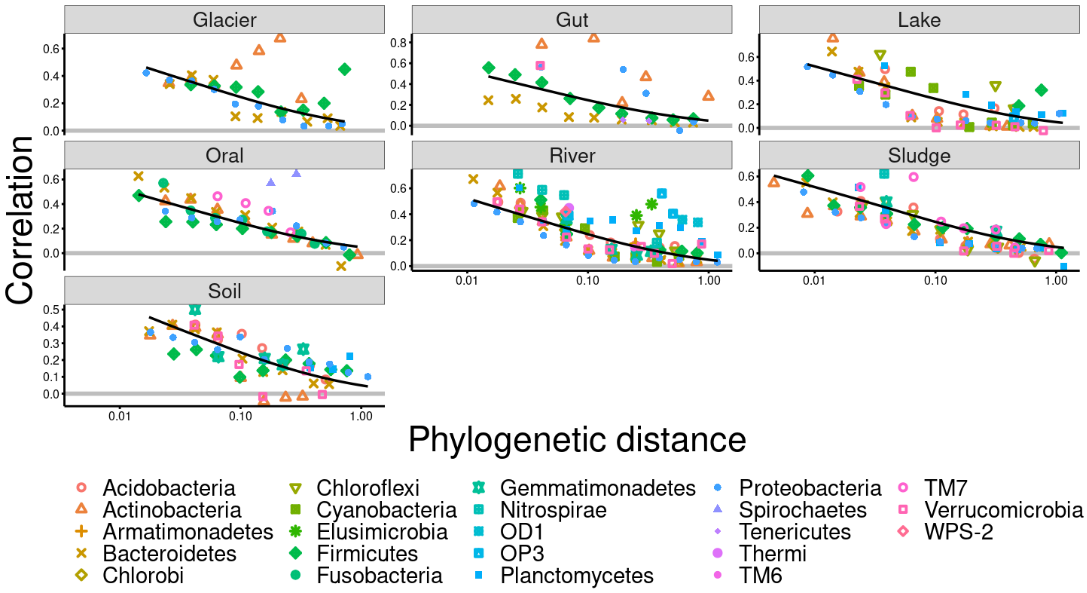


Supplementary Figure S8: Taxonomic Analysis. Intra-phylum correlations versus phylogenetic distance for each biome (see color code and point shapes in legend). Black lines represent the fit with the average parameters (see Fig.S3 B and Table S2) while the symbol sizes represent the log of the number of couples within the bin. The colored points are obtained by averaging only species of the same phylum.

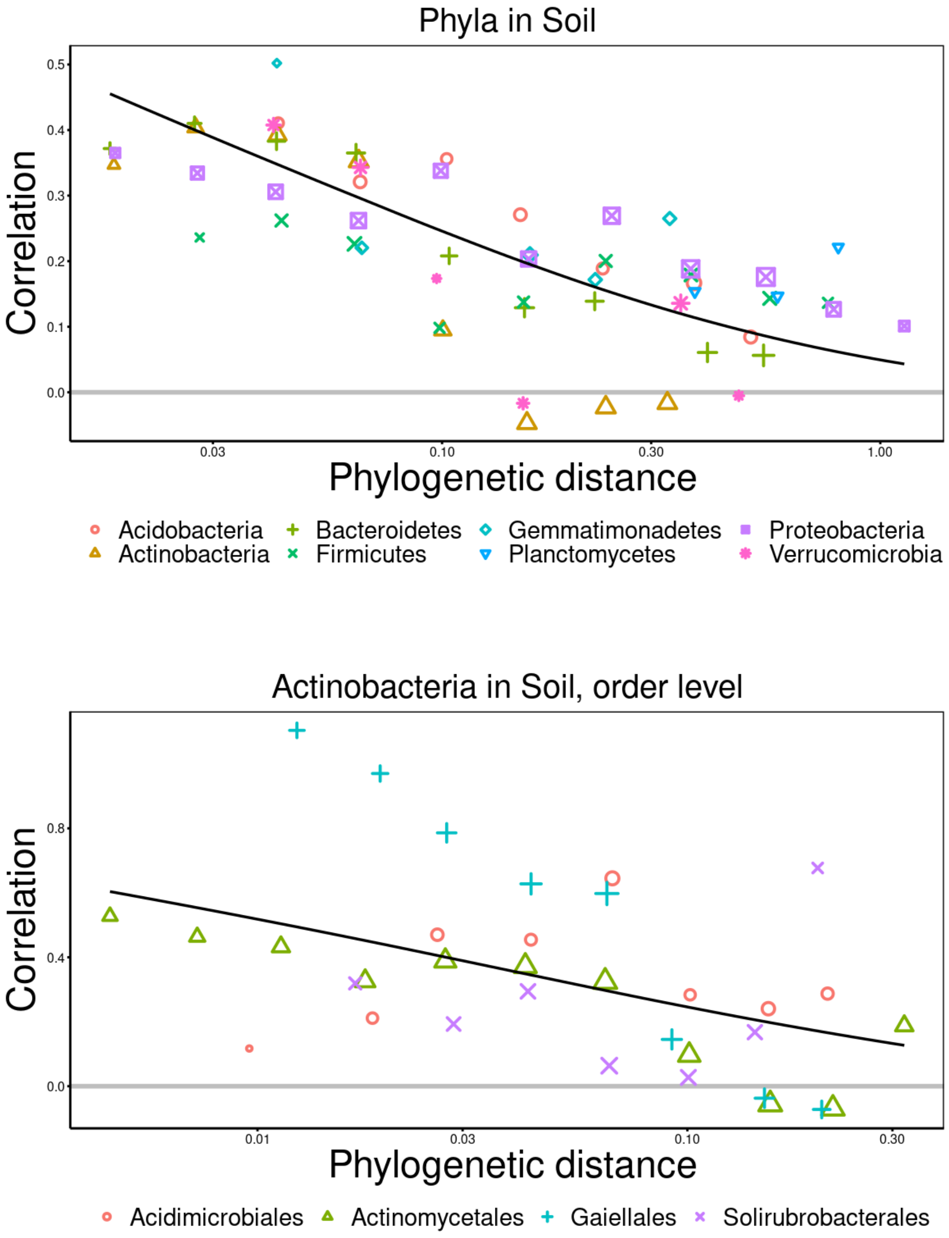


Supplementary Figure S9: **Taxonomic analysis in the soil biome.** Top: intra-phylum correlation pattern in soil; symbol colors and shapes stand for each different phylum (see legend), while symbol sizes stand for the logarithm of the number of couples within each bin. The phylum actinobacteria is an example of a taxa with large deviations from the main pattern, in particular they show negative correlations around phylogenetic distance of 0.1. Also verrucomicrobia exhibit some negative correlation, but are not less abundant than actinobacteria. Bottom: intra-order correlation pattern in the phylum actibacteria in soil biome. By considering the correlation between species of the order inside the actinobacteria phylum we show that the observed deviation is mostly due to actinomycetales and gaiellales.

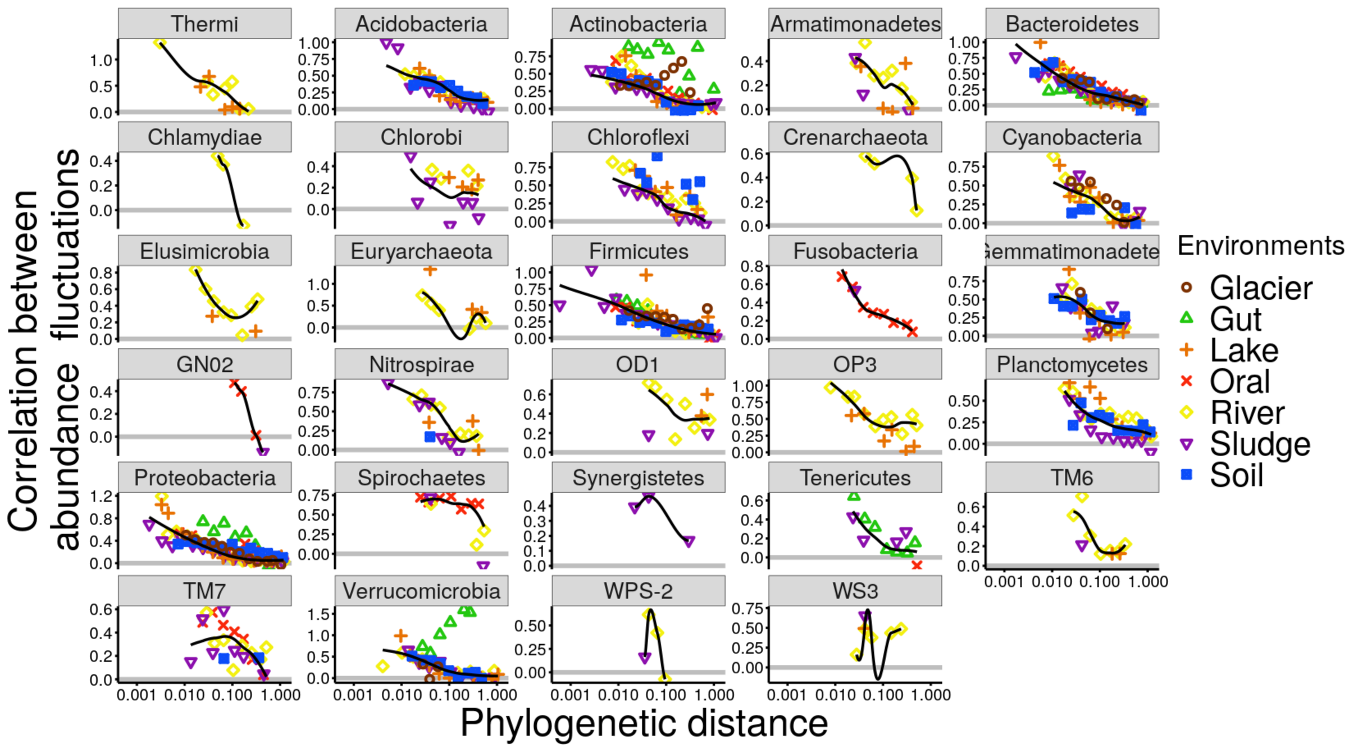


Supplementary Figure S10: **Correlation versus phylogenetic distance in each phylum for different biomes.** Colored points indicate the corresponding biomes and the black line the averaged behavior over biomes weighted by abundances. Correlation of phyla that are resent in many biomes — such as acidobacteria, bacteriotedes, proteobacteria and firmicutes— tend to follow a positive to null decay. On the other hand, less abundant phyla present large deviations for such an overall trend.

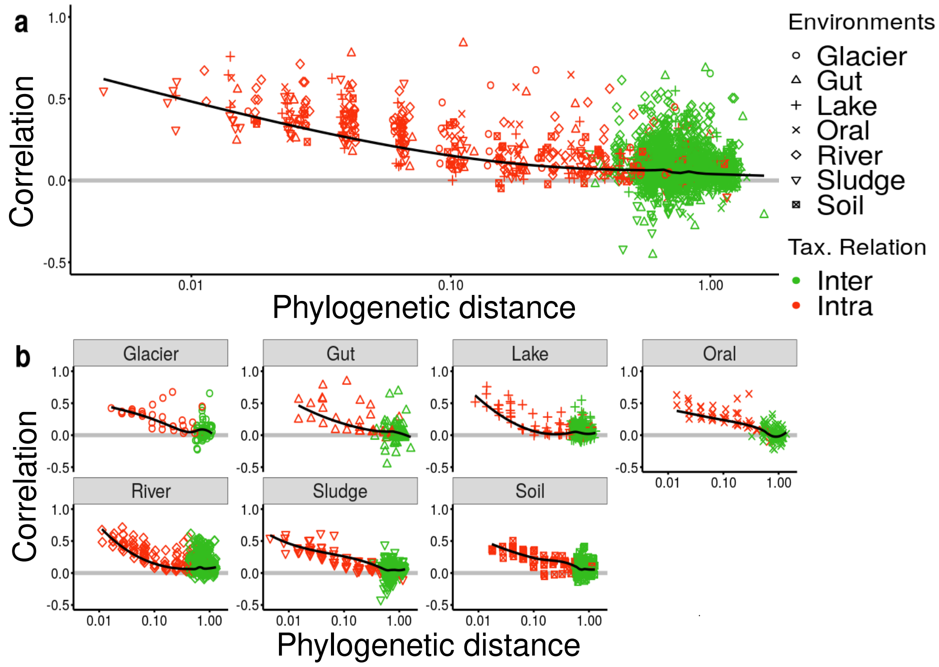


Supplementary Figure S11: **Taxonomic Analyses.** a: Correlation between abundance fluctuations versus phylogenetic distance for intra-phylum (red points) and inter (green points) phyla. In total, 29 phyla are considered and each point represents the correlation of one of them, within a certain phylogenetic distance, in a particular biome (shapes). Black lines are averages over both taxas and biomes, weighted by abundances in each considered bin. (b) Same data as in (a) above but plotted separately for each biome.

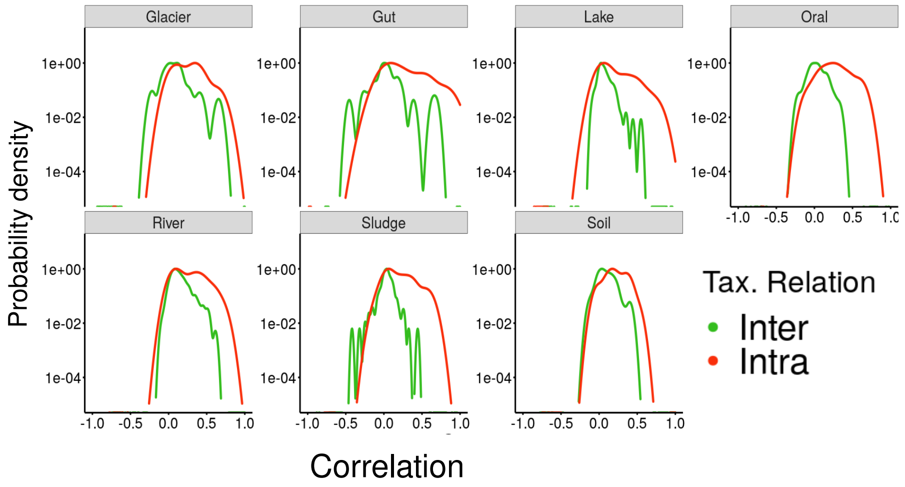


Supplementary Figure S12: **Histogram of correlations for both intra and inter phyla**. As clearly illustrated, inter-taxonomic correlations have a symmetric distribution centered around zero, while intra ones are skewed to positive values.

##### E. Time-series

The analysis of temporal data is analogous to that in Section (S3 A), but instead of studying fluctuations and correlations between different communities, one considers one single community across time. For each host, $h = 1,..,H$, one has different samples from different times (days) $t = 1,..,T$. All the observables are defined as in Sec.(S3 A) but replacing the community average by a time average $\langle..\rangle_t = \frac{1}{T}\sum_{t=1}^{T}(..)$. In particular, the equal time correlation between two species abundance fluctuations $(i, j)$ is defined by:

$$\eta_{kij} = \langle q(t)_{k_i} q(t)_{k_j}\rangle_t = \sum_{t=1}^{T} \frac{q(t)_{k_i} q(t)_{k_j}}{T}; \tag{S16}$$

and the $\Delta t$ delayed correlation as:

$$\eta_{kij}(\Delta t) = \langle q(t+\Delta t)_{k_i} q(t)_{k_j}\rangle_t = \sum_{t=1}^{T-\Delta t} \frac{q(t+\Delta t)_{k_i} q(t)_{k_j}}{T}. \tag{S17}$$

In Fig. S13 and in Table S3, we report the results of fitting the correlations with a stretched-exponential curve as done in Sec.(S3 B). One can see that the correlations follow a stretched exponential of the form of Eq.(S14) with very good approximation, $R^2 \geq 0.9$ in all biomes and hosts, except for the oral biome of the F4 host. Interestingly, there is some variation in the values of $\lambda$ and $\chi$ both across hosts and biomes. In addition, the gut and oral biomes show different but similar parameters in the cross-sectional and longitudinal datasets. Furthermore, the skin biome seems to deviate from the stretched-exponential fit at very small distances coherently across hosts. Nevertheless, observe that the fit of the total pattern to a single stretched exponential is still good, $R^2 > 0.8$, but that the skin biome deviate in totally different way than the oral and gut (see Fig. S14). We leave for future work the investigation of the factors driving such a variability that appears to be larger than in cross-sectional datasets.

Moreover, in Figures S15-S18, we report separately for each biome the behavior of the correlation, for each host and averaging over them, and show that also the delayed correlations can be well fitted by following modification of stretched-exponential function see sec. S4 E 2):

$$\eta(\Delta t, d_G) = \exp\left(-\frac{\Delta t}{\tau} - \lambda d_G^{1/3}\right), \tag{S18}$$

where $\lambda$ and $\chi$ are fixed from the equal-time correlations,see Table S3, and $\tau$ is fixed just once for each biome, $\tau = 1$ for Gut and oral and $\tau = 0.5$ for left and right-palm skin .

| Biome | Host Id | $\chi$ | $\lambda$ | $R^2$ |
|---|---|---|---|---|
| Gut | M3 | $0.31 \pm 0.01$ | $4.18 \pm 0.11$ | 0.98 |
| Gut | F4 | $0.25 \pm 0.01$ | $3.76 \pm 0.04$ | 0.99 |
| Oral | M3 | $0.26 \pm 0.01$ | $3.30 \pm 0.15$ | 0.90 |
| Oral | F4 | $0.20 \pm 0.04$ | $3.18 \pm 0.26$ | 0.54 |
| Skin Left Palm | M3 | $0.38 \pm 0.03$ | $3.77 \pm 0.24$ | 0.96 |
| Skin Left Palm | F4 | $0.39 \pm 0.03$ | $3.81 \pm 0.39$ | 0.92 |
| Skin Right Palm | M3 | $0.47 \pm 0.05$ | $4.08 \pm 0.58$ | 0.89 |
| Total | Total | $0.30 \pm 0.01$ | $3.73 \pm 0.12$ | 0.82 |

Supplementary Table S3: Stretched-exponential fit parameters for each biome for *equal-time* temporal data.

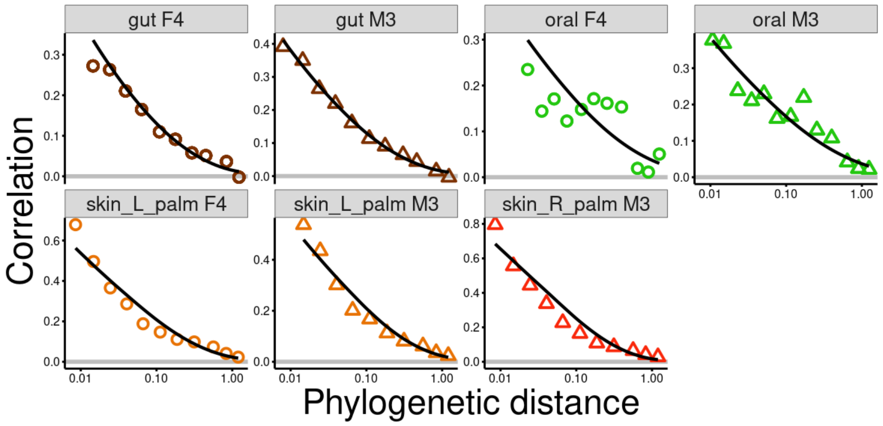


Supplementary Figure S13: **Stretched exponential fit for longitudinal data.** Plots of stretched-exponential fits for different biomes and hosts together with data. Almost all the biomes follow with very good approximation a stretched-exponential decay Eq. (S14), with a the poorest fit in the F4 oral case. Such large deviations are probably due to some unknown conditions of the host. Furthermore, the skin biome consistently deviates from the stretched exponential at small distances.

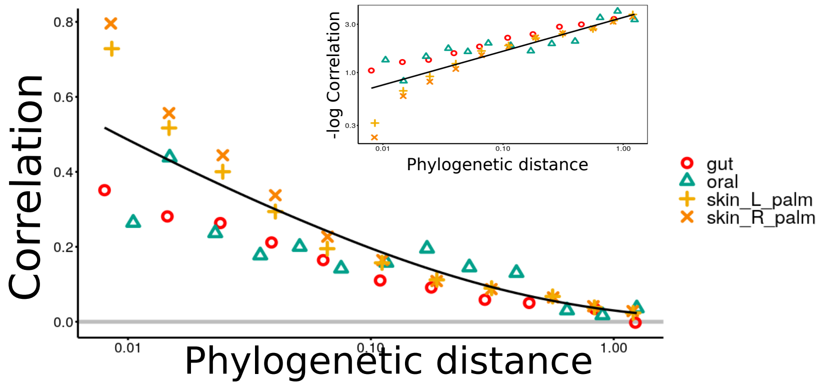


Supplementary Figure S14: **Macroecological law for temporal data: average over hosts and biomes** Correlations as function of phylogenetic distance averaged over hosts for each biomes. Inset: $-log(Corr)$ as function of distance averaged over hosts. Points represent correlations for each biomes averaged over hosts. The black line is the stretched-exponential fit corresponding to the last line reported in the Table S3. Observe that, at small distances, gut and oral biomes have a different behavior than the skin one.

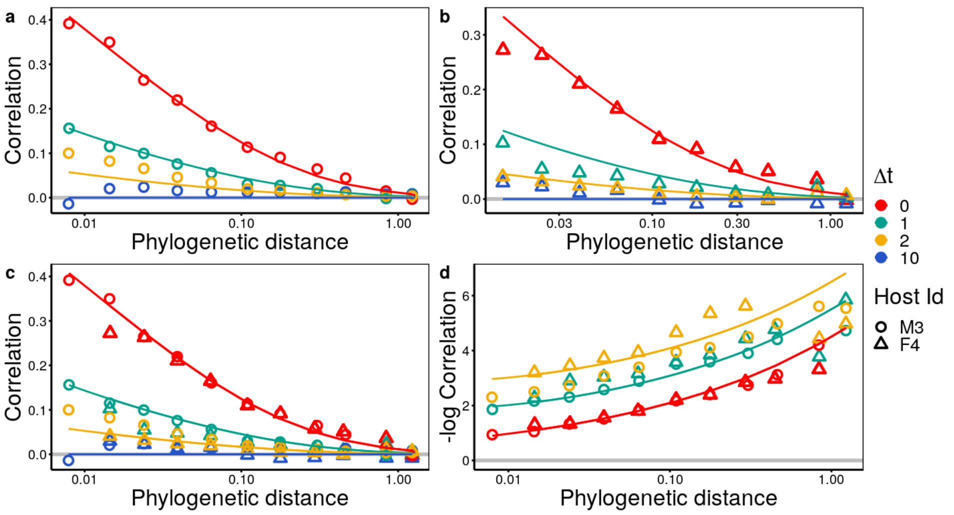


Supplementary Figure S15: **Macroecological law for temporal data: Gut biome.** Correlations as function of phylogenetic distance for each host (a-b), averaged over hosts (c), $-log(Corr)$ as function of distance averaged over hosts (d). Colored points represent correlations at equal time (red) or one, two and ten day delay(green, yellow, blue). Shapes stand for different hosts. Solid lines are the stretched exponential decay from eq.(S18). Notably, the delayed correlations can be predicted using eq.(S18),with the same $\lambda$ and $\chi$ of equal time correlation, just by fixing growth timescale $\tau = 1$. for all the plots. The other parameters for the solid lines are: $(a)\chi = 0.31, \lambda = 4.18; (b)\chi = 0.25, \lambda = 3.76; (c,d)\chi = 0.3, \lambda = 4.$

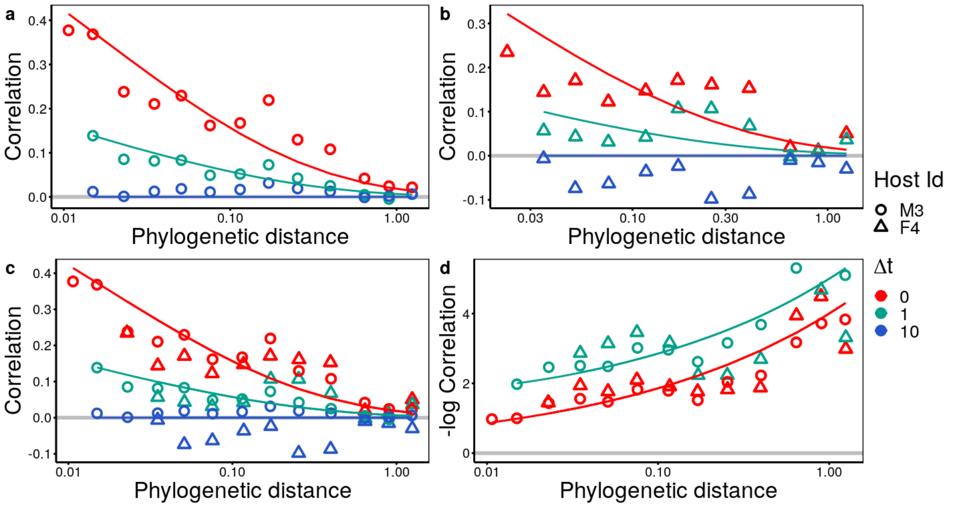


Supplementary Figure S16: **Macroecological law for temporal data: Oral biome.** Symbols, lines and sub-figure legend as in Fig.S15. Observe that the F4 host presents major deviations, that are probably caused by some special health conditions. Nevertheless, the correlations are still decaying from positive to null values, and the delayed pattern follows the same, predictable, tendency of the other biomes. Solid lines are the stretched-exponential decay from Eq.(S18), with $\tau = 1$ in all the plots. The other parameters for the solid lines are: $(a,c,d)\chi = 0.26, \lambda = 3.3; (b)\chi = 0.2, \lambda = 3.18.$

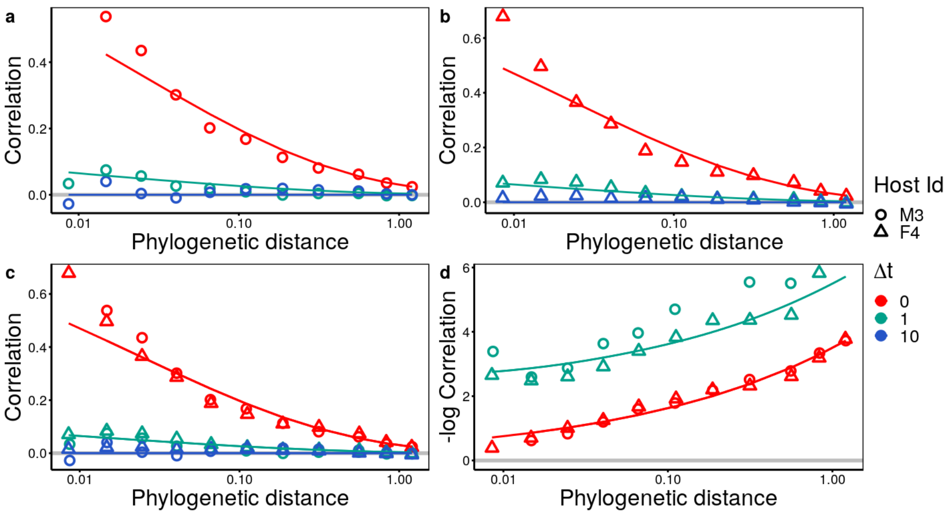


Supplementary Figure S17: **Macroecological law for temporal data: Skin, left- palm biome.** Symbols, lines and sub-figure legend as in Fig.S15. Observe that deviations from the stretched-exponential fit are significant at small phylogenetic distance. Notably, the delayed correlations decay to zero way faster than the other biomes. Both observations are coherent with recent macroecological studies on human microbiome that identified a very rapid dynamics in this skin dataset [9]. Solid lines are the stretched exponential decay from eq.(S18), with $\tau = 0.5$ for all the plots. The other parameters for the solid lines for all plots are: $\chi = 0.38, \lambda = 3.8$.

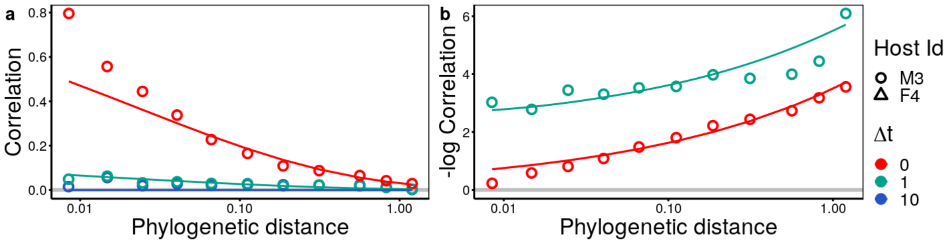


Supplementary Figure S18: **Macroecological law for temporal data: Skin, right- palm biome.** Correlations (a) and $-log(Corr)$versus phylogenetic distance for host M3. Observe that deviations from the stretched exponential and the rapid decay of the delayed correlation are coherent with the left-palm biome, Fig.S17. Solid lines are stretched exponential decay from eq.(S18), with $\tau = 0.5$ The other parameters for the solid lines are: $\chi = 0.47, \lambda = 4.$.

#### S4. MODELS

Consider N species each with an abundance $x_i$, $i \in [1, N]$, growing in a well-mixed environment, exposed to $R$ environmental population-dependent factors(i.e. resources ), $R_\beta$, $\beta \in [1, R]$, and M population-independent factors, $M_\alpha$, $\alpha \in [1, M]$ such as very abundant or rapidly depleted resources or abiotic factors such as temperature and ph. Each species $i$ has different preferences in (i) consuming resources in the case of population-dependent factors like, e.g., sugars and in (b) growing preferentially under certain levels of population-independent ones, as quantified by the $R$ dimensional vector $\boldsymbol{b}^i$ and the M dimensional one $\boldsymbol{a^i}$, respectively. Each of these vectors is constrained on a hypersphere ($M$ or $R$ dimensional) of radius $r_M, r_R$, respectively, i.e.: $|\boldsymbol{b}^i|^2 = \sum_{\beta=1} b^i_\beta = r^2_R$, and $|\boldsymbol{a}^i|^2 = r^2_M$, both of which are fixed to 1 here.

In this way, the per-capita growth rate of each species is influenced both by population-dependent and population-independent factors, weighted by the corresponding preference vector

$$\frac{1}{x_i(t)}\frac{dx_i}{dt} = \sum_{\beta=1}^{R} b^i_\beta R_\beta(t) + \sum_{\alpha=1}^{M} a^i_\alpha M_\alpha(t) - \delta \ , \tag{S19}$$

where $x_i(t)$ is the abundance of species $i$ at time t, $R_\beta(t)$ is the abundance of resource $\beta$ at time $t$, $M_\alpha(t)$ is the value of the population-independent factor $\alpha$, and $\delta$ is the death rate. We assume the population-independent factors are subject to rapid stochastic fluctuations

$$M_\alpha(t) = \bar{M}\left(1 + \sqrt{\nu}\zeta_\alpha(t)\right) \tag{S20}$$

where $\bar{M}$ represents some baseline level and $\zeta_\alpha(t)$ is a Gaussian white noise. The parameter $\nu$ quantifies the strength of population-dependent factors fluctuations. Similarly, the abundance $R_\beta$ of each resource $\beta$ fluctuates in time. In this case, fluctuations are determined by a balance between stochasticity and the consumption by the population:

$$R_\beta(t) = \bar{R}\left(1 + \sqrt{\omega}\varphi_\beta(t) - \gamma\sum_{j=1}^{N} b^j_\beta x_j\right) , \tag{S21}$$

where $\bar{R}$ is a baseline level, $\gamma$ the consumption timescale, and $\varphi_\beta(t)$ a Gaussian white noise with zero mean and variance 1. Similarly to $\sigma$, the parameter $\omega$ quantifies the importance of resources fluctuations. As discussed below this dynamics for consumed resources can be derived from a more-standard consumer resource model.

The model defined in Eq.(S19) can be *effectively* written as a generalized Lotka-Volterra model with fluctuating growth rate and species competition:

$$\frac{dx_i}{dt} = x_i\left(r_i(t) - \sum_{j=1}^{N} C_{ij}x_j\right) \ , \tag{S22}$$

where the entries of the competition matrix are determined by the overlap in resource preference ($C_{ij} = \bar{R}\gamma\boldsymbol{b}^i \cdot \boldsymbol{b}^j$). The time-dependent growth rate can be written as $r_i(t) = \bar{r}_i + \sqrt{\sigma}\xi_i(t)$, where the explicit expressions of $\bar{r}_i$,s and $\xi$ on the original parameters are:

$$\bar{r}_i = \bar{R}\sum_\beta b^i_\beta + \bar{M}\sum_\alpha a^i_\alpha - \delta \tag{S23}$$

$$\sqrt{\sigma}\xi_i(t) = \sqrt{\omega}\bar{R}\sum_\beta b^i_\beta\varphi_\beta + \sqrt{\nu}\bar{M}\sum_\alpha a^i_\alpha\zeta_\alpha \tag{S24}$$

$$\sigma = \nu\bar{M}^2 + \omega\bar{R}^2, \tag{S25}$$

and, additionally, the noise correlations $\langle \xi_i(t)\xi_j(t')\rangle = \rho_{ij}\delta(t-t')$ are given by

$$\rho_{ij} = \frac{\nu \bar{M}^2 \boldsymbol{a}^i \cdot \boldsymbol{a}^j + \omega \bar{R}^2 \boldsymbol{b}^i \cdot \boldsymbol{b}^j}{\nu \bar{M}^2 + \omega \bar{R}^2} \ , \tag{S26}$$

Depending on the geometry of the preference vectors and the type of fluctuations under consideration, one can derive four different general types of models:

- (0)*Fluctuating and non overlapping factors.*

  If all the population-dependent and population-independent factors preference vectors are perpendicular to each other, then species become decoupled and one readily recovers the "stochastic logistic model" (SLM) describing simple environmental fluctuations:

  $$r_i(t) = \bar{r}_i + \xi_i(t), \quad C_{ij} = \bar{R}\gamma\delta_{ij}, \quad \rho_{ij} = \delta_{ij} \tag{S27}$$

- (A)*Shared fluctuating population-dependent factors.*

  If population-independent factor fluctuations are absent, $\nu = 0$, species correlations are determined by population-dependent vectors, which generate a both competition (encoded in the entries $C_{ij}$) and resource fluctuations (encoded in the entries $\rho_{ij}$):

  $$r_i(t) = \bar{r}_i + \sqrt{\omega}\xi_i(t), \quad C_{ij} = \bar{R}\gamma \boldsymbol{b}^i \cdot \boldsymbol{b}^j, \quad \rho_{ij} = \boldsymbol{b}^i \cdot \boldsymbol{b}^j \tag{S28}$$

- (B) *Shared resources and non-overlapping fluctuating population-independent factors.*

  If population-dependent fluctuations are absent, $\omega = 0$, and population-independent factors preferences are all perpendicular to each other, species experience independent growth rate fluctuations and competition for resources.

  $$r_i(t) = \bar{r}_i + \sqrt{\nu}\xi_i(t), \quad C_{ij} = \bar{R}\gamma \boldsymbol{b}^i \cdot \boldsymbol{b}^j, \quad \rho_{ij} = \delta_{ij} \tag{S29}$$

- (C) *Shared fluctuating population-independent factors with fixed non-overlapping resources.*

  If population-dependent fluctuations are absent, $\omega = 0$, and population-dependent factors preferences are all perpendicular to each other, i.e. there are no common population-dependent preferences, species only experience correlated growth rate fluctuations but no inter-specific competition:

  $$r_i(t) = \bar{r}_i + \sqrt{\nu}\xi_i(t), \quad C_{ij} = \bar{R}\gamma\delta_{ij}, \quad \rho_{ij} = \boldsymbol{a}^i \cdot \boldsymbol{a}^j \tag{S30}$$

  Hence this model implements the force called "environmental filtering".

In all these models, we are mostly interested in studying the distribution of the Pearson correlation coefficient, i.e. the correlation between abundance fluctuations:

$$\eta_{i,k} = \frac{\langle x_i x_k\rangle - \langle x_i\rangle\langle x_k\rangle}{\sqrt{Var_i Var_k}}, \tag{S31}$$

as well as its dependence on the species similarity in preferences space, at the stationary state.

Note that in all the considered models just one set or preference vectors is relevant, while the other is perpendicular (population-dependent in A/B population-dependent in 0 and C). Hence, the species dissimilarity can be quantified by the distance in preference space $d_{P,ij}$, as function of the angle $\theta_{ij}$ between the corresponding relevant vector(cosine distance, in the example for population-dependent ones):

$$d_{P,ij} \equiv \frac{2}{\pi}\theta = \frac{2}{\pi}\arccos\left(\frac{\boldsymbol{b^i}\cdot\boldsymbol{b^j}}{|\boldsymbol{b^i}||\boldsymbol{b^j}|}\right) = \frac{2}{\pi}\arccos\left(\frac{\boldsymbol{b^i}\cdot\boldsymbol{b^j}}{r_P}\right). \tag{S32}$$

##### A. Evolutionary algorithms to generate a wide distribution of preference distances.

If vectors in the preference space –identifying the characteristic of each given species in the different models— are generated in a simple random fashion, they have a large probability to be orthogonal to each other; i.e. vectors with small distances are very unlikely to be randomly generated. In particular, as a consequence of the central limit theorem, for sufficiently large numbers of environmental factors, $R$, the random vectors $\boldsymbol{b^i}$ tend to be orthogonal to each other, i.e., $D_{P,ij} \approx 0 \quad \forall i, j)$, hindering the observation of similar species. To paliate this problem and populate all the space of possible pairwise distances we consider two alternative algorithms.

###### 1. *Algorithm 1: high-dimensional preference space*

If the preference space under consideration is high dimensional, i.e. $N \sim R \sim M \ggg 1$, one can employ the following algorithm. We generate the set of $M$ preference vectors $\boldsymbol{b}$ by sampling their component from a Gaussian distribution with mean $m/R$ ($m$ small and positive) and s.t.d. $1/\sqrt{R}$, $\mathcal{N}(m/R, 1/\sqrt{R})$, such that the radius of the overall vector is constant and close to 1 for large $M$: $r_P^2 = \sum_\beta b_\beta^2 = 1 + \frac{m^2}{R} \approx 1$.

Starting from an initial random distribution of vectors $\boldsymbol{b^i}^0$ —and implementing an evolutionary branching process— generates as an outcome a set of vectors $\boldsymbol{b^i}$ which are distributed across all values of possible cosine distances. The algorithm has the following steps:

1. Sample at random two species $i, j$, i reproduces and j dies.
2. Replace the pair of vectors by a new pair, specified by: $\boldsymbol{b^i} = q\boldsymbol{b^i} + (1-q)\boldsymbol{\epsilon}^i$, $\boldsymbol{b^j} = q\boldsymbol{b^i} + (1-q)\boldsymbol{\epsilon}^j$, where the $q \in [0, 1]$ is the "fidelity" and $p = 1 - q$ is the "mutation", and $\epsilon^{i,j}$ vectors sampled from $\mathcal{N}(m/R, 1/\sqrt{R})$.
3. Iterate Z times.

Note that the vectors are automatically kept in the sphere. For large enough values of $Z$ and $q = 0.9$, the population has a small pool of similar individuals, corresponding to a long left tail of the distance distribution (see Fig.S1).

###### 2. *Algorithm 2: low-dimensional preference space*

If the preference space is low dimensional in comparison with the species number, i.e. $N \sim R \ggg M \sim 1$, one can employ an alternative algorithm not relying on the central limit theorem. Consider $N$ vectors of dimension $M$ each generated with just one random non-zero entry sampled by taking the absolute value of a normal distribution $N(m/M, 1/\sqrt{(M)})$, with $m > 0$. Then, the algorithm proceeds as follows:

1. Sample at random two species $i, j$, $i$ reproduces and $j$ dies.
2. Replace both vectors with a mutation of the vector $b_i$ in one random entry, where $q \in [0, 1]$ is the fidelity and $p = 1 - q$ is the mutation. $\boldsymbol{b^i} = q\boldsymbol{b^i} + q\boldsymbol{\epsilon}^i$, $\boldsymbol{b^j} = q\boldsymbol{b^i} + (1-q)\boldsymbol{\epsilon}^j$, with $q \in [0, 1]$ is the fidelity of reproduction and $\epsilon^{i,j} = (0, .., \epsilon_k^{i,j}, 0, .., 0)$ with k chosen at random between 1 and $M$ and $\epsilon_k^{i,j}$ sampled by $\mathcal{N}(m/R, 1/\sqrt{R})$.
3. Iterate Z times.

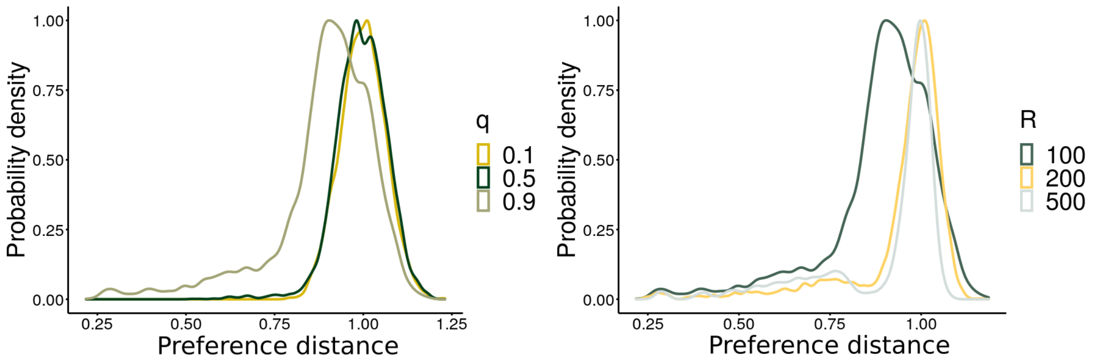


Supplementary Figure S19: **Preference distance distribution generated by the algorithm in high-dimensional space.** Left: distribution obtained for different values of $q$ and $R = 100$. Right: distribution with $q = 0.9$ and different values of R. As a consequence of the central limit theorem, the distribution converges to a Gaussian for small values of $q$ and $R \to \infty$. On the contrary, for high values of fidelity, $q = 0.9$, the distribution develops a long left tail, covering all the preference space. As shown in the right figure for any value of $q$, the variance shrinks by increasing $R$. In both figures we considered $N = 100$, and iterated the algorithm $Z = 50N$.

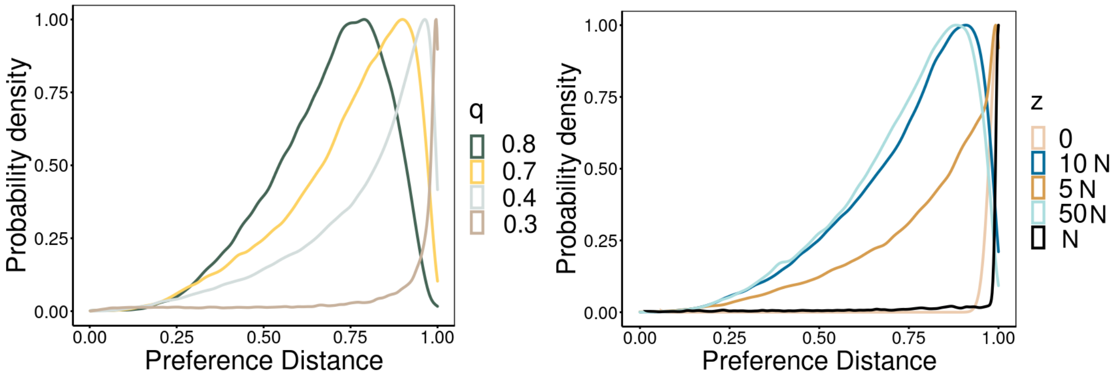


Supplementary Figure S20: **Preference distance distribution generated by the second algorithm for low-dimensional preference spaces** Left: distribution for different values of q and $N = 500, M = 10, Z = 10N$. Right: distribution with $q = 0.7$ and different values of Z.

If the fidelity is moderate but and $Z$ is large the distance distribution develops a long tail independently on the dimension of $M$, see Fig. S20

##### B. Model 0: Fluctuating and non overlapping factors.

As a first analysis, we use the Stochastic Logistic model as a null expectation for the correlations. Such a model is easily derived if both the population-dependent and population-independent factors preference vectors are perpendicular to each other (for simplicity one component for vector with modulus m)

$$\bar{r}_i = (\bar{R} + \bar{M})m - \delta, \quad C_{ij} = m\bar{R}\gamma\delta_{ij}, \quad \sigma_i = \nu\bar{M}^2 + \omega\bar{R}^2, \quad \rho_{ij} = \delta_{ij}. \tag{S33}$$

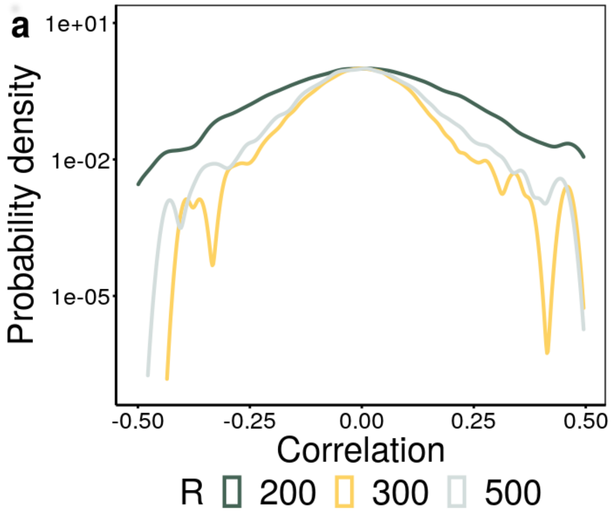


Supplementary Figure S21: **Correlations for Model 0.** (a) Stationary probability distribution (log-scale) of correlations for different values of $R$ (color coded) and fixed $N$. Parameters: $N = 100, K_i = 10.0, \bar{r}_i = 1.0, \sigma_i' = 0.1, q = 0.9, Z = 50N$, and $t_{fin} = 10^4$.

Indeed, such a model describes logistic growth plus environmental fluctuations of N uncoupled species:

$$\dot{x}_i = x_i \left( \bar{r} - \frac{x_i}{K} \right) + x_i \sqrt{\sigma} \xi_i \tag{S34}$$

$$\langle \xi_i(t) \rangle = 0 \tag{S35}$$

$$\langle \xi_i(t) \xi_j(t') \rangle = \delta_{ij} \delta(t - t') \tag{S36}$$

where the carrying capacity is $K = (\bar{R}\gamma)^{-1}$. By considering the following ansatz on the parameters:

$$\bar{r}_i = \frac{1}{\tau_i}, \tag{S37}$$

$$K_i = \tau_i K_i', \tag{S38}$$

$$\sigma_i = \frac{\sigma_i'}{\tau_i}, \tag{S39}$$

one recovers the standard form of the stochastic logistic model:

$$\dot{x}_i = \frac{x_i}{\tau_i} \left( 1 - \frac{x_i}{K_i'} \right) + \sqrt{\frac{\sigma_i'}{\tau_i}} \xi_i(t) x_i \tag{S40}$$

$$\langle \xi_i(t) \rangle = 0 \tag{S41}$$

$$\langle \xi_i(t) \xi_j(t') \rangle = \delta_{ij} \delta(t - t'). \tag{S42}$$

Observe that, by definition, the preference distance is always one, and hence this model gives a symmetric distribution of correlations centered in zero, with spurious variance due to finite size effects that converges to zero in the infinite species and resources limit, see Fig.S21.

##### C. Model A:Shared fluctuating population-dependent factors.

The presence of population-dependent factors (such as, e.g., sugars) in the environment fluctuates due to seasonality, fluxes with external space etc., inducing both competition between species and fluctuating growth. Thus, if population-independent factors are absent, species interactions are determined by a combination of the effect of competition and resources fluctuations:

$$\begin{aligned}\dot{x}_i &= x_i\left(\bar{r}_i - \sum_j C_{ij}x_j\right) + \sqrt{\sigma}\xi_i(t)x_i \\ \langle\xi_i\rangle &= 0, \\ \langle\xi_i(t)\xi_j(t')\rangle &= \delta(t-t')\rho_{ij};\end{aligned} \tag{S43}$$

with:

$$\bar{r}_i = \bar{R}\sum_\beta b^i_\beta + \bar{M}m - \delta, \quad C_{ij} = \bar{R}\gamma \boldsymbol{b}^i\cdot\boldsymbol{b}^j, \quad \sigma_i = \omega\bar{R}^2, \quad \rho_{ij} = \boldsymbol{b}^i\cdot\boldsymbol{b}^j \tag{S44}$$

The population-dependent factors preference vector are generated as described in S4 A, Importantly, in the limit $R \gg 1$, the model does not depend on the preference vector but just on the angles between them. Indeed, in such a limit the growth rate depends just on the mean preference

$$\bar{r}_i = \bar{R}R\langle b^i\rangle + \bar{M} - \delta = (\bar{R}+\bar{M})m - \delta. \tag{S45}$$

Furthermore, the preference vector modulus, i.e. the hyper-sphere radius converges to 1:

$$r_P^2 = \sum_\beta b_\beta^2 = 1 + \frac{m^2}{R} \approx 1, \tag{S46}$$

and hence species interactions depend directly on the preference distance:

$$C_{ij} = \bar{R}\gamma\cos(\theta_{ij}) = \bar{R}\gamma\cos\left(\frac{\pi}{2}d_{P,ij}\right) \tag{S47}$$

$$\rho_{ij} = \cos(\theta_{ij}) = \bar{R}\gamma\cos\left(\frac{\pi}{2}d_{P,ij}\right). \tag{S48}$$

Curiously enough, this model leads to a consistent number of extinctions, and the surviving communities are "effectively neutral", i.e. with species pair at almost all distances but with average correlation zero, see Fig. S22.

##### D. Model B: Shared resources and non-overlapping fluctuating population-independent factors.

We now consider that population-dependent factors do not fluctuate in time, i.e. $\omega = 0$, and that population-independent factors preferences are all perpendicular to each other. Hence species experience independent growth rate fluctuations and competition for resources. Such a model results in competition in preference space for $N$ species and $R$ resources in the form of stochastic Lotka-Volterra equation:

$$\begin{aligned}\dot{x}_i &= x_i\left(r_i(t) - \sum_{j=1}^N C_{ij}x_j\right) \\ &= x_i\left(\bar{r}_i - \sum_{j=1}^N C_{ij}x_j\right) + \sqrt{\sigma_i}\xi_i(t)x_i,\end{aligned} \tag{S49}$$

with:

$$\bar{r}_i = \bar{R}\sum_\beta b^i_\beta + \bar{M}m - \delta, \quad C_{ij} = \bar{R}\gamma\boldsymbol{b}^i\cdot\boldsymbol{b}^j, \quad \sigma = \nu\bar{M}^2 \quad \rho_{ij} = \delta_{ij}. \tag{S50}$$

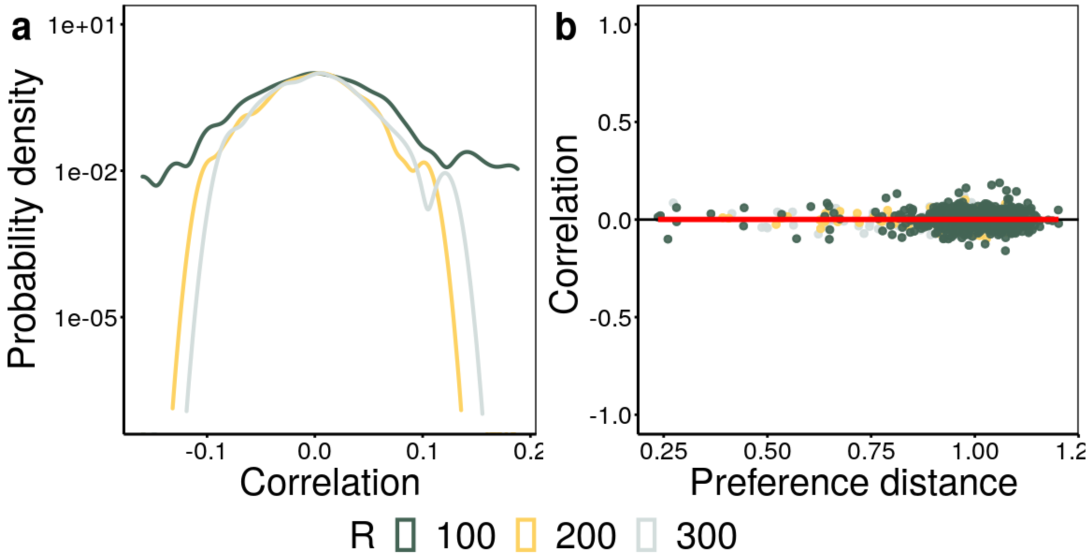


Supplementary Figure S22: **Correlations in Model A.** (a) Stationary probability distribution (log-scale) of correlations, and (b) correlations as function of preference distance for different values of $R$ (color coded) and fixed $N$. The red line in (b) is the correlation averaged over all the presented realizations. As a consequence of limiting similarity various extinction events happen before reaching the stationary state, resulting in the majority of species having distance one, and just a few species in the small distance tail. Note that the correlations do not depend on the species distance, leading to and effective neutral behavior. Parameters: $R = 100, 200, 300$. $q = 0.1$, $Z = 50N$,$N = 100, m = 0.5\bar{R} = \bar{M} = 1, \delta = 0.1, \gamma = 0.3, \omega = 1$.

Furthermore, by considering the limit $R \gg 1$ we obtain here a simplified version of the model depending only on the the preference distance matrix $d_{P,ij}$ and not on the vectors $\boldsymbol{b^i}$. Indeed, the deterministic growth rate reads:

$$\bar{r}_i = \bar{R}R\langle b^i \rangle + \bar{M}m - \delta = (\bar{R} + \bar{M})m - \delta, \tag{S51}$$

$$C_{ij} = \bar{R}\gamma\cos(\theta_{ij}) = \bar{R}\gamma\cos\left(\frac{\pi}{2}d_{P,ij}\right). \tag{S52}$$

We leave for a further work the analytic calculation of the Pearson correlation in this case. Here, we limit ourselves to show numerically that there is a robust dependence of correlations on the preference distance: for small distances the correlation is negative and it increases converging to 0 at large distances, Eq.S23.

##### E. Model C: Shared fluctuating population-independent factors with fixed non-overlapping resources.

If resource fluctuations are absent, $\omega = 0$, and population-dependent factors preferences are all perpendicular to each other, i.e. no common resource preferences, species experience dependent growth rate fluctuations and no inter-specific competition:

$$\dot{x}_i(t) = x_i\left(\bar{r}_i - \frac{x_i}{K_i}\right) + \sqrt{\sigma}\xi_i(t)x_i, \tag{S53}$$

where:

$$\bar{r}_i = \bar{M}\sum_{\alpha} a^i_\alpha + m\bar{R} - \delta, \quad K_i = (\gamma\bar{R})^(-1), \quad \rho_{ij} = \boldsymbol{a}^i \cdot \boldsymbol{a}^j \tag{S54}$$

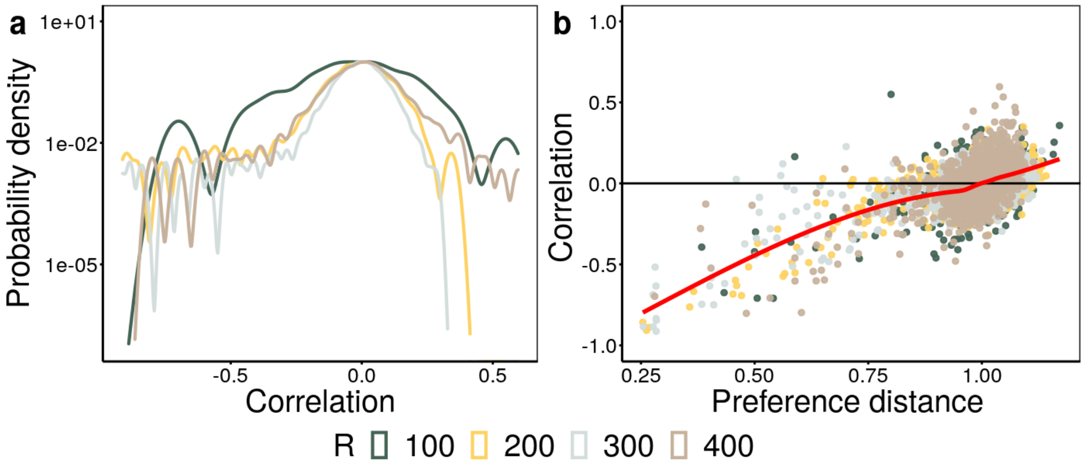


Supplementary Figure S23: **Correlations for Model B** (a) Stationary probability distribution (log-scale) of correlations, and (b) correlations as function of preference distance for different values of $R$ (color coded) and fixed $N$. The red line in (b) is the correlation averaged over all the presented realizations. As a consequence of limiting similarity various extinction events happen before reaching the stationary state, resulting in the majority of species having distance one, and just a few species in the small distance tail. Parameters: $q = 0.1$, $Z = 50N$,$N = 100, m = 0.5\bar{R} = \bar{M} = 1, \delta = 0.1, \gamma = 0.3, \omega = 1$.

Hence this model implements the force called "Environmental Filtering". By taking the limit $M \gg 1$, we can recast the model just in terms of preference distance $d_{P,ij}$. The deterministic growth rate reads:

$$\bar{r}_i = \bar{M}\langle a^i\rangle + m\bar{R} - \delta = (\bar{R} + \bar{M})m - \delta; \tag{S55}$$

while the noise correlation can be rewritten as:

$$\rho_{ij} = \boldsymbol{a}^i \cdot \boldsymbol{a}^j = \cos(\frac{\pi}{2}\theta_{ij}), \tag{S56}$$

cause

$$|a|^2 = \frac{m^2}{M} + 1 \approx 1. \tag{S57}$$

Finally, by rescaling the parameter with the growth rate as follow:

$$\bar{r}_i = \frac{1}{\tau_i} \tag{S58}$$

$$K_i' = K_i\tau_i^{-1} = \frac{(\bar{R} + \bar{M})m - \delta}{\gamma\bar{R}} \tag{S59}$$

$$\sigma_i = \frac{\sigma_i'}{\tau_i}, \tag{S60}$$

one obtains the “correlated stochastic logistic” model (CSLM):

$$\frac{dx_i}{dt} = \frac{x_i}{\tau_i}\left(1 - \frac{x_i}{K_i}\right) + \sqrt{\frac{\sigma}{\tau_i}}\xi_i(t)x_i \tag{S61}$$

$$\langle\xi_i\rangle = 0 \tag{S62}$$

$$\langle\xi_i(t)\xi_j(t')\rangle = \cos\left(\frac{\pi}{2}d_{P,ij}\right), \tag{S63}$$

(where the notation has been simplified).

In the following section S4 E 1, we show that in the linear approximation around the fixed point, the stationary Pearson correlation reads:

$$\eta_{ij}(d_{P,ij}) = \frac{\exp\left(\cos(\frac{\pi}{2}d_{P,ij})\frac{\sigma}{2-\sigma}\right) - 1}{\exp\left(\frac{\sigma}{2-\sigma}\right) - 1} \approx \cos\left(\frac{\pi}{2}d_{P,ij}\right), \tag{S64}$$

in the case $\sigma_i = \sigma_j, \tau_i = \tau_j$.

The decay of Pearson correlation coefficients with preference distance, Eq. (S64), is confirmed by numerical simulations, see Fig.(S24. Note, in particular, that the deterministic mean preference $m$ divided by the resource number, sets the average correlation, but its decrease implies a large number of extinctions. Furthermore, increasing the number of resources for a constant number of species decreases the mean correlation, its variance (by decreasing the distance distribution one).

In all the models considered here, the effective carrying capacities are constant across species. Here we show how to generalize the calculations to include a carrying capacity distribution. To generalize this it is sufficient to consider the consumption parameter timescale $\gamma_i$, to be species dependent. In the present model case each resource is consumed exclusively by one species ($N = R$), hence:

$$R_i(t) \ = \ \bar{R}\left(1 - m\gamma_i x_i\right), \tag{S65}$$

leading to an effective carrying capacity (rescaled by the growth rate):

$$K_i \ = \ \frac{\left(\bar{R} + \bar{M}\right)m - \delta}{m^2\gamma_i\bar{R}} \tag{S66}$$

Finally, let us note that the decay of correlation applies also to communities with only two species: if we consider a pair of species with a given level of similarity, we obtain a correlation which is identical to the one observed in the full community (see Fig. S26). To obtain such a pattern, we fixed that the modulus of all preference vectors to one, i.e. $|\boldsymbol{a}_i| = 1$ for $i = 1, 2$ and also fixed preference vector the first species as $\boldsymbol{a}_1 = (1, 0)$. In each simulation (i.e. for each gray point in the figure), we choose the desired preference distance between the two species and derived the component of the second vector as:

$$\begin{aligned} a_2^1 \ &= \ \cos\left(\frac{\pi}{2}d_P\right) \\ a_2^2 \ &= \ \sqrt{1 - (a_2^1)^2}. \end{aligned} \tag{S67}$$

In the next section (S4 E 1) we present a more careful explanation on why the pattern is also valid only for two species (at tunable distances).

##### *1. Linear approximation around the fixed point*

Here we derive the stationary correlation of a couple of species by employing a linear expansion around the fixed point. This approximation assumes that stochastic fluctuations are small and hence the dynamics is localized nearby the deterministic fixed point. By looking at Eq. (S61), it is clear that the deterministic fixed point for each species is $x_i^* = K_i$. Nevertheless, owing to the presence of multiplicative fluctuations it is not possible to perform the analysis in these variable, but it is necessary to linearize the noise-correlation matrix and recast the equations into an independent

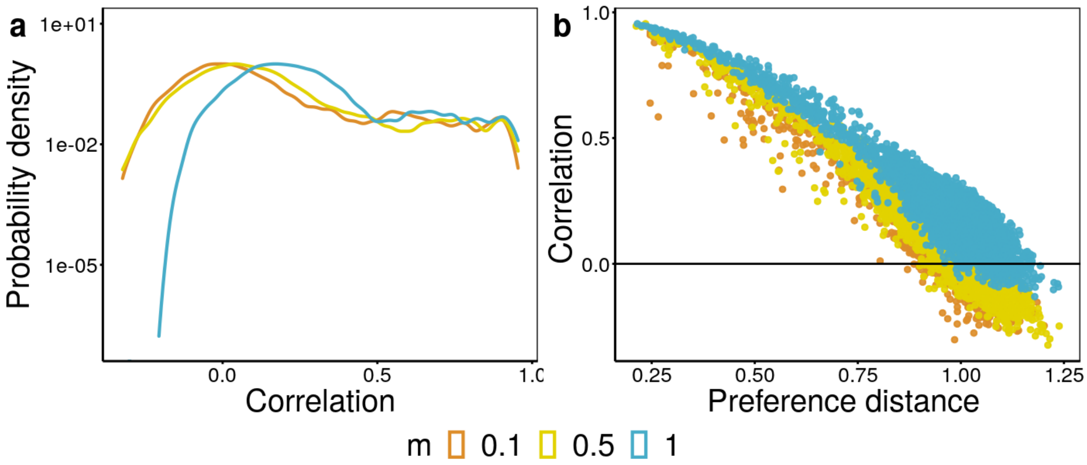


Supplementary Figure S24: **Correlation in Model C: dependence on m.** Pearson correlation distribution (log-scale) (a) and correlations as function preference distance (b) for different values of $m$ (color coded), for $m = 0.1, 0.5, 1$. Parameters: $N = M = 100, q = 0.9, S = 50N, \bar{R} = \bar{M} = 1, \gamma = 0.1, \nu_i = 0.5, t_{fin} = 10^4$.

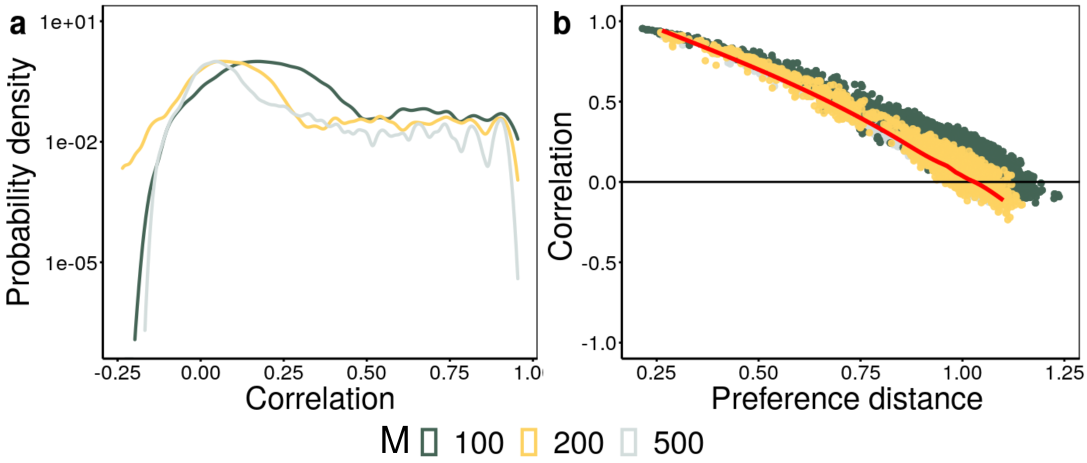


Supplementary Figure S25: **Correlation in Model C: dependence on the number of resources.** Pearson correlation distribution (log-scale) (a) and correlations as function preference distance (b) for different values of $M$ (color coded), $M = 100, 200, 500$. The red line in (b) stands for eq.$S64$. As expected, by increasing the number of resources, the correlation distribution variances decreases, and the decay is well-approximated by the linear noise approximation. Parameters: $N = 100, q = 0.9, S = 50N, m = 1, \bar{R} = \bar{M} = 1, \gamma = 0.1, \nu_i = 0.5$, and $t_{fin} = 10^4$.

additive noise form. In these new variables one can easily perform a linear expansion and, afterward, come back to the original ones. As a first step, one can apply a logarithmic change of variables to make the noise additive:

$$u_i = log(x_i), \tag{S68}$$

by using the Ito formula Eq. S61 becomes:

$$\dot{u}_i = \frac{\dot{x}_i}{x} = \frac{1}{\tau_i}\left(1 - \frac{\sigma}{2}\right) - \frac{e^{u_i}}{\tau_i K_i} + \sqrt{\frac{\sigma_i}{\tau_i}}\zeta_i(t). \tag{S69}$$

As a second step, one needs to find eigenvalues and eigenvectors of the correlation matrix. Given that we are interested in the two-species correlations, we restrict ourselves here, without loss of generality, to an arbitrary couple of species

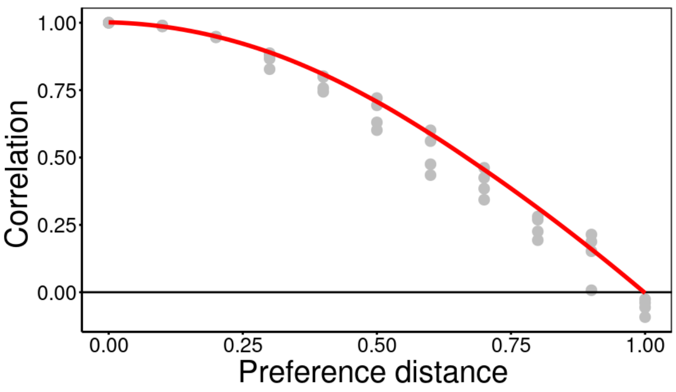


Supplementary Figure S26: **Correlation in Model C for couple of species** Model C of environmental filtering at the stationary state where just two species with a given preference distance per realization are considered. Each gray point represent the stationary Pearson correlation of the couple, while the red line in stands for eq.$S64$. As expected, the pattern is still valid. Parameters: $N = 2, \boldsymbol{a}_1 = (1., 0), \gamma = 0.1, \nu = 0.5$, and $t_{fin} = 10^4$

$(i, j)$. Furthermore, for sake of notation let us rename the parameters as:

$$W_i = \sqrt{\frac{\sigma_i}{\tau_i}}, \tag{S70}$$

$$\rho_{ij} = \rho = \cos\left(\frac{\pi}{2} d_{P,ij}\right), \tag{S71}$$

and include these factors in the noise correlation matrix:

$$C_{ij} = \begin{pmatrix} W_i^2 & W_i W_i \rho_{ij} \\ W_i W_j \rho_{ij} & W_j^2 \end{pmatrix}. \tag{S72}$$

Hence, the eigenvalues of the reduced correlation matrix are:

$$\lambda_{i,j} = \frac{1}{2}(W_i^2 + W_j^2 \pm \sqrt{W_i^4 + 2(2\rho^2 - 1)W_i^2 W_j^2 + W_j^4}), \tag{S73}$$

and the corresponding eigenvectors

$$\nu_{i,j} = \left( -\frac{W_j^2 - W_i^2 \pm \sqrt{W_i^4 + 2(2\rho^2 - 1)W_i^2 W_j^2 + W_j^4}}{2 W_i W_j \rho}, 1 \right). \tag{S74}$$

Note that we have to impose $\rho \neq \pm 1$ in order to avoid having a degenerate transformation. For the sake of simplicity let us assume the following ansatz on the parameters:

$$W_i = W_j \to \tau_i = \tau_j = \tau, \sigma_i = \sigma_j = \sigma. \tag{S75}$$

In this case, the eigenvalues and eigenvectors reduce to:

$$\lambda_{i,j} = W^2(1 \pm \rho), \tag{S76}$$

and

$$\nu_{i,j} = \frac{1}{\sqrt{2}}(1, \pm 1), \tag{S77}$$

where we have also normalized the vectors. Now, it is possible to decompose the noise as:

$$\zeta_i = \sum_\mu \sqrt{\lambda_\mu} v_{\mu,i} \eta_\mu(t), \tag{S78}$$

with

$$\begin{aligned} \langle \eta_\mu \rangle &= 0, \\ \langle \eta_\mu(t)\eta_\gamma(t') \rangle &= \delta(t-t')\delta_{\mu,\gamma}; \end{aligned} \tag{S79}$$

and the dynamical variable as

$$p_\mu = \sum_i u_i \nu_{\mu,i}. \tag{S80}$$

In our simple case the variables are:

$$\begin{aligned} p_i &= \frac{u_i + u_j}{\sqrt{2}}, \\ p_j &= \frac{u_i - u_j}{\sqrt{2}}. \end{aligned} \tag{S81}$$

Hence, the new Langevin equations are

$$\begin{aligned} \dot{p}_i &= \frac{\sqrt{2}}{\tau} - \frac{1}{\sqrt{2}\tau}\left(\frac{e^{\frac{p_i+p_j}{\sqrt{2}}}}{K_i} + \frac{e^{\frac{p_i-p_j}{\sqrt{2}}}}{K_j} + \sigma\right) + \sqrt{\frac{\sigma}{\tau}(1+\rho)}\eta_i(t), \\ \dot{p}_j &= -\frac{1}{\sqrt{2}\tau}\left(\frac{e^{\frac{p_i+p_j}{\sqrt{2}}}}{K_i} - \frac{e^{\frac{p_i-p_j}{\sqrt{2}}}}{K_j}\right) + \sqrt{\frac{\sigma}{\tau}(1-\rho)}\eta_j(t), \\ \langle \eta_i(t) \rangle &= 0, \\ \langle \eta_i(t)\eta_j(t') \rangle &= \delta_{ij}\delta(t-t'). \end{aligned} \tag{S82}$$

Note that the deterministic force admits a scalar potential

$$V(p_i, p_j) = -\frac{\sqrt{2}p_i}{\tau} + \frac{1}{\tau}\left(\frac{e^{\frac{p_i+p_j}{\sqrt{2}}}}{K_i} + \frac{e^{\frac{p_i-p_j}{\sqrt{2}}}}{K_j} + \frac{\sigma}{\sqrt{2}}p_j\right), \tag{S83}$$

and the minimum of the potential gives the fixed point of the dynamics, namely:

$$(p*_i, p_j^*) = \frac{1}{\sqrt{2}}\left(ln\left[\left(1-\frac{\sigma}{2}\right)^2 K_i K_j\right], ln\left(\frac{K_i}{K_j}\right)\right). \tag{S84}$$

Importantly, note that the fixed point, when converted to the original variables, is different from the fixed point of the deterministic dynamics in eq.(S61):

$$(x_i^*, x_j^*) = (K_i(1-\frac{\sigma}{2}), K_i(1-\frac{\sigma}{2})). \tag{S85}$$

If one now considers $\sigma^2 \ll 1$ it is interesting to study the linear fluctuations around the fixed point, like in the small-noise approximation:

$$p_i = p_i^* + \delta p_i, \tag{S86}$$

the Jacobian evaluated at the fixed point reads:

$$J_{p^*} = -\frac{1}{2\tau}(1-\frac{\sigma}{2})\begin{pmatrix} \frac{e^{\frac{p_i+p_j}{\sqrt{2}}}}{K_i} + \frac{e^{\frac{p_i-p_j}{\sqrt{2}}}}{K_j} & \frac{e^{\frac{p_i+p_j}{\sqrt{2}}}}{K_i} - \frac{e^{\frac{p_i-p_j}{\sqrt{2}}}}{K_j} \\ \frac{e^{\frac{p_i+p_j}{\sqrt{2}}}}{K_i} - \frac{e^{\frac{p_i-p_j}{\sqrt{2}}}}{K_j} & \frac{e^{\frac{p_i+p_j}{\sqrt{2}}}}{K_i} + \frac{e^{\frac{p_i-p_j}{\sqrt{2}}}}{K_j} \end{pmatrix}_{p^*} \tag{S87}$$

$$= -\frac{1}{\tau}\left(1-\frac{\sigma}{2}\right)\begin{pmatrix} 1 & 0 \\ 0 & 1 \end{pmatrix} \tag{S88}$$

Therefore, one ends up with the following linearized dynamics

$$\dot{\boldsymbol{\delta p}} = J\boldsymbol{\delta p} + \boldsymbol{\eta}, \tag{S89}$$

with the Jacobian above and (rescaled) diffusion matrix:

$$D = \begin{pmatrix} D_i & 0 \\ 0 & D_j \end{pmatrix} = \frac{W^2}{2}\begin{pmatrix} 1+\rho & 0 \\ 0 & 1-\rho \end{pmatrix}, \tag{S90}$$

such that

$$\langle \eta_\mu(t)\eta_\gamma(t')\rangle = 2D_{\mu,\gamma}\delta(t-t'). \tag{S91}$$

We use the following notation for the Jacobian:

$$J = -\begin{pmatrix} J_i & -J_{ij} \\ -J_{ji} & J_j \end{pmatrix}. \tag{S92}$$

In this scenario, it is know that the stationary distribution is a bivariate Gaussian with covariance matrix[7]

$$\Sigma = \frac{-1}{Tr(J)det(J)}\begin{pmatrix} D_jJ_{ij}^2 - D_iJ_{ij}J_{ji} + D_iJ_j(J_i+J_j) & D_jJ_iJ_{ij} + D_iJ_{ji}J_j \\ D_jJ_iJ_{ij} + D_iJ_{ji}J_j & D_iJ_{ji}^2 - D_jJ_{ij}J_{ji} + D_jJ_i(J_i+J_{ij}) \end{pmatrix} \tag{S93}$$

$$= \frac{\sigma}{2-\sigma}\begin{pmatrix} 1+\rho & 0 \\ 0 & 1-\rho \end{pmatrix},$$

$$P^*(\delta p_1, \delta p_2) = \frac{2-\alpha\sigma}{2\pi\sigma\sqrt{1-\rho^2}}e^{-\left(\frac{1}{\sigma}-\frac{\alpha}{2}\right)\left(\frac{\delta p_1^2}{1+\rho}+\frac{\delta p_2^2}{1-\rho}\right)}. \tag{S94}$$

Now it is necessary to go back to the original variables $x_i$, taking care of the modulus of the determinant of the Jacobian of the transformation $H$:

$$\begin{aligned} \boldsymbol{\delta p} &= H(\boldsymbol{x}), \\ \delta p_i = p_i - p_i^* &= \frac{1}{\sqrt{2}}ln\left(\frac{x_ix_j}{\left(1-\frac{\sigma}{2}\right)K_iK_j}\right), \\ \delta p_j = p_j - p_j^* &= \frac{1}{\sqrt{2}}ln\left(\frac{x_1K_2}{K_1x_2}\right), \end{aligned} \tag{S95}$$

$$
\begin{aligned}
P(\boldsymbol{x}) &= P(\boldsymbol{\delta p}(\boldsymbol{x}))|det(dH)|) = \frac{1}{x_i x_j} P(\boldsymbol{\delta p}(\boldsymbol{x})) \qquad \text{(S96)}\\
&= \frac{2-\sigma}{2\pi x_i x_j \sigma \sqrt{1-\rho^2}} \\
&\quad e^{-\left(\frac{2-\sigma}{2\sigma(1-\rho^2)}\right)\left[ln^2\left(\frac{x_i}{(1-\sigma/2)K_i}\right)+ln^2\left(\frac{x_j}{(1-\sigma/2)K_2}\right)-2\rho ln\left(\frac{x_j}{(1-\sigma/2)K_i}\right)ln\left(\frac{x_j}{(1-\sigma/2)K_j}\right)\right]}.
\end{aligned}
$$

Note that, for the implicit assumptions of the approximation, this distribution is a log-normal one, while we know that the marginal distribution needs to be a Gamma. This can be appreciated from the moments:

$$
\begin{aligned}
\langle x_i \rangle &= K_i \left(1-\frac{\sigma}{2}\right) e^{\frac{\sigma}{2(2-\sigma)}} &\text{(S97)}\\
\langle x_i x_j \rangle &= K_i K_j \left(1-\frac{\sigma}{2}\right)^2 e^{\frac{(1+\rho_{ij})\sigma}{2-\sigma}} &\text{(S98)}\\
\langle x_i^2 \rangle &= K_i^2 \left(1-\frac{\sigma}{2}\right)^2 e^{\frac{2\sigma}{2-\sigma}} &\text{(S99)}\\
Var_i &= K_i^2 \left(1-\frac{\sigma}{2}\right)^2 (e^{\frac{2\sigma}{2-\sigma}} - e^{\frac{\sigma}{2-\sigma}}). &\text{(S100)}
\end{aligned}
$$

Nevertheless, one can calculate the covariance

$$
Cov_{ij} = \langle x_i x_j \rangle - \langle x_i \rangle \langle x_j \rangle = K_i K_j \left(1-\frac{\sigma}{2}\right)^2 (e^{\frac{(1+\rho_{ij})\sigma}{2-\sigma}} - e^{\frac{\sigma}{2-\sigma}}), \qquad \text{(S101)}
$$

and finally the Pearson correlation coefficient:

$$
\begin{aligned}
\eta_{ij} &= \frac{Cov_{ij}}{\sqrt{Var_i Varx_j}} = \frac{e^{(1+\rho_{ij})\frac{\sigma}{2-\sigma}} - e^{\frac{\sigma}{2-\sigma}}}{e^{\frac{2\sigma}{2-\sigma}} - e^{\frac{\sigma}{2-\sigma}}} &\text{(S102)}\\
&= \frac{e^{\rho\frac{\sigma}{2-\sigma}} - 1}{e^{\frac{\sigma}{2-\sigma}} - 1} \simeq \rho_{ij} = cos(\frac{\pi}{2} d_{P,ij}). &\text{(S103)}
\end{aligned}
$$

##### *2. Temporal behavior*

Eq. (S61) can be formally solved exactly, leading to:

$$
\begin{aligned}
x_i(t) &= \frac{K_i \tau x_i(0) e^{(1-\frac{\sigma}{2})\frac{t}{\tau}+\sqrt{\frac{\sigma}{\tau}}W_i(t)}}{x_i(0) I_i[0,t] + 1}, &\text{(S104)}\\
W_i(t) &= \int_0^t ds \zeta_i(s), &\text{(S105)}
\end{aligned}
$$

where $I_i[0,t]$ is the integral of the associated "geometric Brownian motion":

$$
I_i[0,t] = \int_0^t ds \exp\left((1-\frac{\sigma}{2})\frac{s}{\tau} + \sqrt{\frac{\sigma}{\tau}} W_i(s)\right). \qquad \text{(S106)}
$$

The exact integral eq.(S104) can be used to understand the effect of delay in the dynamics. Namely, assuming that the system is in its stationary state ($t \to \infty$), one can compute heuristically the relation between the abundance at time $t$ and at a later time $t + \Delta t$:

$$
\begin{aligned}
x_i(t+\Delta t) &\approx K_i \tau e^{(1-\frac{\sigma}{2})\frac{t+\Delta t}{\tau}+\sqrt{\frac{\sigma}{\tau}}W_i(t+\Delta t)} I_i[0, t+\Delta t]^{-1} \\
&= x_i(t) \exp\left((1-\frac{\sigma}{2})\frac{\Delta t}{\tau}\right) \kappa(t, \Delta t), &\text{(S107)}\\
\kappa(t, \Delta t) &= \exp\left(\sqrt{\frac{\sigma}{\tau}} \int_t^{t+\Delta t} \zeta_i(s)\right) \left(\frac{I_i[0,t]}{I_i[0, t+\Delta t]}]\right); &\text{(S108)}
\end{aligned}
$$

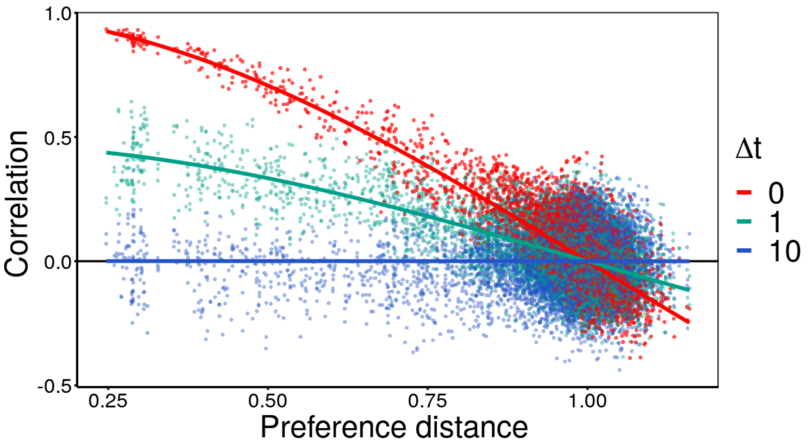


Supplementary Figure S27: Delayed correlations as function of preference distance. Points represent species person correlation coefficient at stationary state at equal time (red) and with delay 1 (green), and 10 (blue) for one realization of the model. Solid lines represent the analytically-derived formula Eq.(S111). Parameters are equal for all species, the variability is given to the variability in preferences distribution, as plotted in Fig.S27. Parameters: $N = 100, M = 100, q = 0.9, Z = 50N, m = 0.5, \omega = 0.1, \bar{r} = 1, K = e^2$, and $t_{fin} = 10^4$.

the function $\kappa$ in the limit $t \gg \Delta t$ converges to 1, such that

$$x_i(t + \Delta t) \approx e^{(1-\frac{\sigma}{2})\frac{\Delta t}{\tau}} x_i(t). \tag{S109}$$

By inserting it in the Pearson coefficient formula we obtain:

$$\eta_{ij}(\Delta t) \approx e^{(1-\frac{\sigma}{2})\frac{\Delta t}{\tau}} \eta_{ij}(0), \tag{S110}$$

that combined with the linear approximation, Eq.(S102), finally leads to:

$$\eta_{ij}(\Delta t) \approx e^{(1-\frac{\sigma}{2})\frac{\Delta t}{\tau}} \cos\left(\frac{\pi}{2} d_{P,ij}\right) \tag{S111}$$

which quantifies analytically the time-delayed correlations.

##### F. Model D: Purely abiotic fluctuations

To explicit consider population-independent factors such as abiotic factors like temperature or Ph, we propose here a modification of the previous models where such factors affect the growth rate in a “multiplicative way”. The per-capita growth rate of each species is, as above, influenced both by population-dependent and population-independent factors, weighted by the corresponding preference vector

$$\frac{1}{x_i(t)} \frac{dx_i}{dt} = M_i(t) \sum_{\beta=1}^{R} b^i_\beta R_\beta(t) - \delta \ , \tag{S112}$$

As above, we assume the population-independent factors to be subject to rapid stochastic fluctuation

$$M_i(t) = \bar{M} + \sum_{\alpha=1}^{M} \sqrt{\nu} a_{i\alpha} \zeta_\alpha(t), \tag{S113}$$

where $\bar{M}$ represents some baseline level and $\zeta_\alpha(t)$ is a Gaussian white noise and $\nu$ quantifies the strength of abiotic factors fluctuations. Similarly, the abundance $R_\beta$ of each resource $\beta$ fluctuates over time. In this case, fluctuations are determined by a balance between stochasticity and the consumption by populations:

$$R_\beta(t) = \bar{R}\left(1 + \sqrt{\omega}\varphi_\beta(t) - \gamma\sum_{j=1}^{M} b^j_\beta x_j\right), \tag{S114}$$

where $\bar{R}$ is a baseline level, $\gamma$ the consumption timescale, and $\varphi_\beta(t)$ a Gaussian white noise with zero mean and variance 1. Similarly to $\sigma$, the parameter $\omega$ quantifies the importance of resources fluctuations. The model defined in Eq. (S19) can be *effectively* written as a generalized Lotka-Volterra model with fluctuating growth rate and species competition matrix:

$$\begin{aligned}\frac{1}{x_i}\frac{dx_i(t)}{dt} &= \bar{M}\sum_{\beta=1}^{R} b^i_\beta\left(\bar{R} + \sqrt{\omega}\varphi_\beta(t) - \gamma\sum_{j=1}^{N} b^j_\beta x_j\right) + \sqrt{\nu}\bar{R}\sum_\beta b_{i\beta}a_{i\alpha}\zeta_\alpha(t) - \delta \\ &+ \sqrt{\nu\omega}\sum_{\alpha,\beta} a_{i\alpha}b_{i\beta}\zeta_\alpha(t)\varphi_\beta(t) - \sqrt{\nu}\gamma\sum_{\alpha,\beta}\sum_j a_{i\alpha}b_{i\beta}b_{j\beta}x_j\zeta_\alpha(t).\end{aligned} \tag{S115}$$

First, let us note that if the fluctuation parameters $\nu, \omega$ and the consumption timescale $\gamma$ are small, all the terms on the second line are sub-leading and the models is equivalent to the general case defined in the first part of section(S4).

For the sake of simplicity, here we study a reduced version of the model, that we call Model D, where resources do not fluctuate and their preferences vectors are all perpendiculars:

$$\frac{1}{x_i}\frac{dx_i(t)}{dt} = \left(\bar{M} + \sqrt{\nu}\sum_{\alpha=1}^{M} a_{i\alpha}\zeta_\alpha(t)\right)\left(\bar{r}_i - \frac{x_i}{K_i}\right) \tag{S116}$$

with

$$\bar{r}_i = \sum_{\beta=1}^{R} b_{i\beta} \tag{S117}$$

$$K_i^{-1} = |\boldsymbol{b}_i|^2\gamma. \tag{S118}$$

To realistically model abiotic factors they need to appear in a number smaller than the number of species, i.e. $N \sim R \ggg M$. In order to obtain a distance distribution with the desired properties in such a scaling regime, we use the evolutionary algorithm exposed in sec.S4 A 2. Based on a linear-noise perturbation we expect the stationary pattern of correlations to not deviate much from model C. Indeed, Fig.S28 shows the numerical solution for the average correlation as function of preference distance still decays in good numerical agreement with the Eq. (S64.

##### G. Model derivation from a consumer-resource framework

In this section we provide a sketch of how the general models introduced in previous sections can be derived from a more-general and standard consumer-resource setting. We leave for a future publication the full presentation and derivation [8].

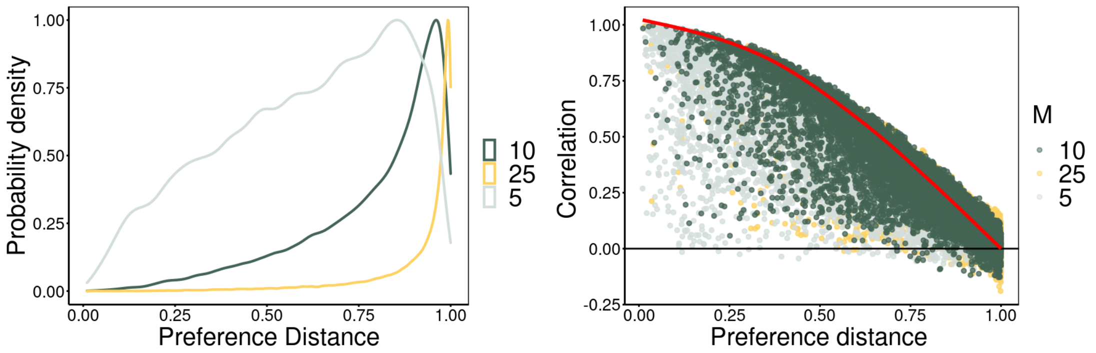


Supplementary Figure S28: **Correlation in Model D: dependence on the number of abiotic factors** Model D of environmental filtering at the stationary state. Left: preference distance distribution for differente values of M (color coded), $M = 5, 10, 25$ and $N = 300$. Right: Pearson correlation distribution (log-scale) and correlations as function preference distance for different values of M (color coded), $M = 5, 10, 25$ and $N = 300$. The red line stands for Eq.S64. As expected, by increasing the number of resources, the correlation distribution variances decreases and the decay is well-approximated by the linear noise approximation. Parameter values: $N = 300, q = 0.4, Z = 50N, m = 1, \bar{R} = \bar{M} = 1, \gamma = 0.1, \nu_i = 0.5$, and $t_{fin} = 10^4$

Consider $N$ species consuming $R$ fluctuating resources, :

$$\dot{x}_i = x_i M_i(t) \sum_{\beta=1}^{R} R_j b^i_\beta - \delta_i x_i \tag{S119}$$

$$\dot{R}_\beta = \lambda_\beta - \mu_\beta R_\beta - \gamma R_\beta \sum_{j=1}^{N} b^j_\beta x_j + \sqrt{\omega}_\beta \varphi_\beta(t), \tag{S120}$$

$$\langle \varphi_\beta(t) \rangle = 0, \tag{S121}$$

$$\langle \varphi_\beta(t) \varphi_\alpha(t') \rangle = \delta_{\alpha\beta}(t - t'), \tag{S122}$$

where species $i$ preference for resource $\beta$ is represented by the component of the vector $\boldsymbol{b}^i$, $b^i_\beta$, $\delta_i$ is the species death rate, $\lambda_\beta$ and $\mu_\beta$ the resource entrance and exit rate, $\gamma$ the consumption timescale and $\omega$ the fluctuation amplitude of the resource noise $\varphi_{beta}$. The influence of abiotic factors is modeled by the $M_i(t)$ as:

$$M_i(t) = \bar{M} + \sum_{\alpha=1}^{M} \sqrt{\nu} a_{i\alpha} \zeta_\alpha(t), \tag{S123}$$

where $\bar{M}$ represents some baseline level and $\zeta_\alpha(t)$ is a Gaussian white noise. The parameter $\nu$ quantifies the strength of abiotic factors fluctuations. If the resource-generation rate is larger than the typical consumption and that exit rates, i.e. if the resources dynamics is *faster* than the species one, it is possible to use a timescales separation technique to remove the explicit dynamic of resources and obtain the following effective model:

$$\dot{x}_i = x_i \left( M_i(t) \sum_{\beta=1}^{R} b^i_\beta \frac{\lambda_\beta + \sqrt{\omega_\beta} \varphi_\beta}{\mu_\beta + \gamma \sum_{j=1}^{N} b^j_\beta x_j} - \delta_i \right). \tag{S124}$$

By considering $R \ggg 1$,$\gamma \lll \mu$ and Taylor expanding Eq. (S124) around $\gamma = 0$ up to first order in $\gamma$ and in $1/R$, one obtains a generalized Lotka-Volterra model equivalent to the one presented in sec.S4 and S4 F:

$$\dot{x}_i = x_i M_i(t) \left( \sum_{\beta=1}^{R} b^i_\beta \left( \frac{\lambda_\beta}{\mu_\beta} + \frac{\sqrt{\omega_\beta}}{\mu_\beta} \varphi_\beta \right) - \gamma \sum_{j}^{N} C_{ij}(t) x_j \right) - \delta_i x_i, \tag{S125}$$

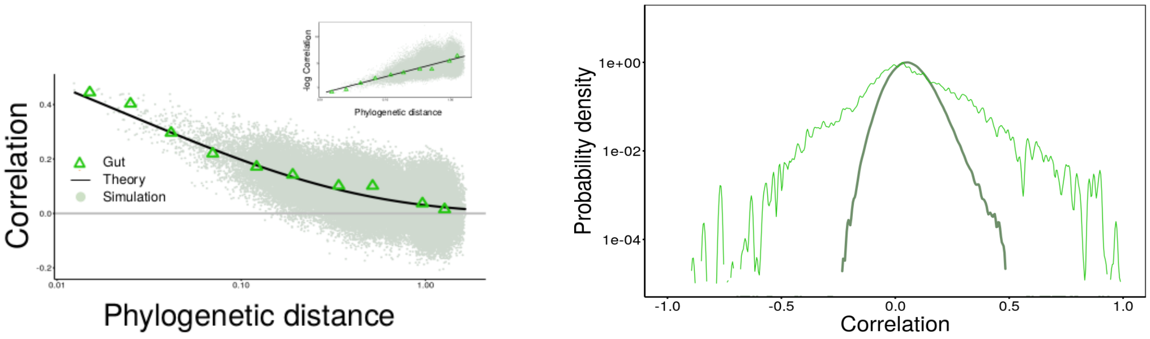


Supplementary Figure S29: **CSLM in phylogenetic space for the Gut microbiome** Left: Correlations versus the phylogenetic distance, both for the gut biome data (green triangles), the model simulation (green cloud of points), and analytic formula eq.(S134)(black line) $\lambda = 3.5$. The model has been simulated 10 times with $N = 300$ species, using as input the empirical phylogenetic distance matrix of the gut, randomly sampling from it N species. Inset: -log Correlations vs phylogenetic distance in log-log scale, empirically and from the model, same data as the main figure. Dark-green points are the Pearson correlation coefficient at the stationary state. Right: Correlation distribution for data and model (log-scale). Light green line is the empirical probability density of correlation for the gut microbiome, while the dark green one corresponds to the correlation distribution of the CSLM (dark green points in left figure.) Carrying capacities are generated log-normally by taking the exponential of random variables sampled by a Gaussian $N(\bar{K}, \sigma_K)$, $\tau_i = \tau$ and $\sigma_i = \sigma$ for $i = 1, .., N$. Parameters: $\tau = 1, \bar{K} = 16.1, \sigma_K = 3.8, \sigma = 1.42, \lambda = 3.5, t_f = 10^4$.

with the competition matrix specified by:

$$C_{ij} = \sum_{\beta=1}^{R} b_\beta^i b_\beta^j \frac{\lambda_\beta}{\mu_\beta^2}. \quad \text{(S126)}$$

##### H. CSLM in phylogenetic space

To compare the CSLM with empirical data it is necessary to connect the preference distance with the phylogenetic one. Let us $d_{G,ij}$ and $d_{P,ij}$ the phylogenetic and phenotypic distance of the couple of species $ij$, and $d_G = \langle d_{G,ij} \rangle_{d_{G,ij} \in b}$, $d_P = \langle d_{P,ij} \rangle_{d_{P,ij} \in b}$ the average genetic and preference distance within a given bin. Note that the empirical relationship we aim at reproducing is between the average correlation and the averaged phylogenetic distance in each bin, i.e.:

$$\eta(d_G) = \langle \eta_{ij} \rangle_{d_{G,ij} \in b} = \exp(-\lambda d_G^{1/3}), \quad \text{(S127)}$$

Indeed, we are not interested in the full probability distribution of correlation in one bin, but just in its mean value. Hence, it is sufficient to find a relation between the average distance $d_P$ and $d_G$, and not between the the full matrices. On the other hand, the CSLM relates the species correlation with their preference distance:

$$\eta_{ij} = \cos(\frac{\pi}{2} d_{P,ij}); \quad \text{(S128)}$$

hence, the average preference distance can be calculated by inverting the formula for the correlation as

$$d_{P,ij} = \frac{2}{\pi} arcos(\eta_{ij}), \quad \text{(S129)}$$

and by taking averages over the couples in the considered bin:

$$d_P(b) \;=\; \left\langle d_{P,ij} \right\rangle_b = \left\langle \frac{2}{\pi} \arccos(\eta_{ij}(d_{G,ij})) \right\rangle_b \tag{S130}$$

To evaluate the last term it would be necessary to know the exact distribution of phylogenetic distance in each bin, but such a distribution is highly non-universal and difficult to quantify. To circumvent the problem, we apply a "mean-field approximation", i.e. we neglect the variance in each bin and consider just its mean:

$$\begin{aligned} d_P(b) \;&=\; \left\langle \frac{2}{\pi} \arccos(\eta_{ij}(d_{G,ij})) \right\rangle_b \approx \frac{2}{\pi} \arccos\left( \left\langle \eta_{ij}(d_{G,ij}) \right\rangle_b \right) \\ &=\; \frac{2}{\pi} \arccos(\eta(d_G)) = \frac{2}{\pi} \arccos\left( e^{-\lambda d_G^{1/3}} \right). \end{aligned} \tag{S131}$$

To incorporate such a relation in the CSLM let us note once more that we are interested in average pattern of correlation as function phylogeny, and hence it is sufficient to assume Eq. (S131) true also for the full phylogenetic matrix, i.e.:

$$d_{P,ij} \approx \frac{2}{\pi} \arccos\left( e^{-\lambda d_{G,ij}^{1/3}} \right). \tag{S132}$$

Putting together all these ingredients, one finally obtains the CSLM in phylogenetic space:

$$\begin{aligned} \frac{dx_i}{dt} \;&=\; \frac{x_i}{\tau_i}\left(1 - \frac{x_i}{K_i}\right) + \sqrt{\frac{\sigma_i}{\tau_i}} x_i \xi_i(t) \\ \langle \zeta_i(t) \rangle \;&=\; 0 \\ \langle \zeta_i(t) \zeta_j(t') \rangle \;&=\; e^{-\lambda d_{G,ij}^{1/3}}. \end{aligned} \tag{S133}$$

By applying Eq.(S111) to this specific setting, one finally obtains a theoretical prediction relating correlation and phylogeny:

$$\eta_{ij} \;=\; e^{-\lambda d_{G,ij}^{1/3}}, \tag{S134}$$

$$\eta_{ij}(\Delta t) \;=\; e^{-(1-\frac{\sigma}{2})\frac{\Delta t}{\tau}} e^{-\lambda d_{G,ij}^{1/3}}. \tag{S135}$$

In Fig(S29) we show that the model correctly predicts the decaying pattern of correlation versus phylogenetic distance. However, it cannot reproduce the full correlation distribution. The correlations are obtained from 10 realizations with $N = 300$ species, the averaged are over $10^3$ abundances sampled during the stationary time series every $\delta_t = 10\tau$. To generate the noise correlations, in each realization we sampled $N$ species from the phylogenetic distance matrix of random community of considered biome. Parameters are set to reproduce also the species marginal properties and delayed correlations, see Figs.(S15-S18).

---


[1] R. Ambrosini, F. Musitelli, F. Navarra, I. Tagliaferri, I. Gandolfi, G. Bestetti, C. Mayer, U. Minora, R. S. Azzoni, G. Diolaiuti, C. Smiraglia, and A. Franzetti. Diversity and assembling processes of bacterial communities in cryoconite holes of a karakoram glacier. *Microbial ecology*, 73(4):827–837, 05 2017.

[2] J. G. Caporaso, C. L. Lauber, E. K. Costello, D. Berg-Lyons, A. Gonzalez, J. Stombaugh, D. Knights, P. Gajer, J. Ravel, N. Fierer, J. I. Gordon, and R. Knight. Moving pictures of the human microbiome. *Genome Biology*, 12:R50 – R50, 2011.

[3] P. J. McMurdie and S. Holmes. phyloseq: An r package for reproducible interactive analysis and graphics of microbiome census data. *PLOS ONE*, 8(4):1–11, 04 2013.

[4] A. L. Mitchell, M. Scheremetjew, H. Denise, S. Potter, A. Tarkowska, M. Qureshi, G. A. Salazar, S. Pesseat, M. A. Boland, F. M. I. Hunter, P. ten Hoopen, B. Alako, C. Amid, D. J. Wilkinson, T. P. Curtis, G. Cochrane, and R. D. Finn. EBI Metagenomics in 2017: enriching the analysis of microbial communities, from sequence reads to assemblies. *Nucleic Acids Research*, 46(D1):D726–D735, 10 2017.

[5] J. P. NiñoâGarcía, C. RuizâGonzález, and P. A. Giorgio. Interactions between hydrology and water chemistry shape bacterioplankton biogeography across boreal freshwater networks. *The ISME Journal*, 10:1755–1766, 2016.

[6] C. Quast, E. Pruesse, P. Yilmaz, J. Gerken, T. Schweer, P. Yarza, J. Peplies, and F. O. Glockner. The SILVA ribosomal RNA gene database project: improved data processing and web-based tools. *Nucleic Acids Research*, 41(D1):D590–D596, 11 2012.

[7] H. Risken and H. Haken. *The Fokker-Planck Equation: Methods of Solution and Applications Second Edition.* Springer, 1989.

[8] M. Sireci and M. Muñoz. The stochastic chemostat model. *In preparation.*

[9] S. Zaoli and J. Grilli. A macroecological description of alternative stable states reproduces intra- and inter-host variability of gut microbiome. *Science Advances*, 7(43):eabj2882, 2021.
